# Chemoarchitectural studies of the rat hypothalamus and zona incerta. *Chemopleth 1.0* – A downloadable interactive *Brain Maps* spatial database of five co-visualizable neurochemical systems, with novel feature- and grid-based mapping tools

**DOI:** 10.1101/2024.10.02.616213

**Authors:** Vanessa I. Navarro, Alexandro Arnal, Eduardo Peru, Sivasai Balivada, Alejandro R. Toccoli, Diana Sotelo, Olac Fuentes, Arshad M. Khan

**Author notes:** Address correspondence to: Arshad M. Khan, Ph.D. or Vanessa I. Navarro.

## Abstract

The hypothalamus and zona incerta of the brown rat (*Rattus norvegicus*), a model organism important for translational neuroscience research, contain diverse neuronal populations essential for survival, but how these populations are structurally organized as systems remains elusive. With the advent of novel gene-editing technologies and artificial intelligence, there is an apparent research need for high-spatial-resolution maps of rat hypothalamic neurochemical cell types to aid in their gene-directed targeting, to validate their expression in transgenic lines, or to supply precious ground-truth training data for machine learning algorithms. Here, we present *Chemopleth 1.0* [available at: https://doi.org/10.5281/zenodo.15788189], a chemoarchitecture database for the rat hypothalamus (HY) and zona incerta (ZI), which features downloadable interactive maps featuring the census distributions of five immunoreactive neurochemical systems: (1) vasopressin (as detected from its gene co-product, copeptin); (2) neuronal nitric oxide synthase (<u>EC 1.14.13.39</u>); (3) hypocretin 1/orexin A; (4) melanin-concentrating hormone; and (5) alpha-melanocyte-stimulating hormone. These maps are formatted for the widely used *Brain Maps 4.0* (BM4.0) open-access rat brain atlas (<u>RRID:SCR_017314</u>). Importantly, this dataset retains atlas stereotaxic coordinates that facilitate the precise targeting of the cell bodies and/or axonal fibers of these neurochemical systems, thereby potentially serving to streamline delivery of viral vectors for gene-directed manipulations. The maps are presented together with novel open-access tools to visualize the data, including a new workflow to quantify cell positions and fiber densities for BM4.0. The workflow produces “heat maps” of neurochemical distributions from multiple subjects: 1) *isopleth maps* that represent consensus distributions independent of underlying atlas boundary conditions, and 2) *choropleth maps* that provide distribution differences based on cytoarchitectonic boundaries. The database files, generated using the Adobe® Illustrator® vector graphics environment, can also be opened using the free vector graphics editor, Inkscape. We also introduce a refined grid-based coordinate system for this dataset, register it with previously published spatial data for the HY and ZI, and introduce novel grid-based annotation of experimental observations. This database provides critical spatial targeting information for these neurochemical systems unavailable from mRNA-based maps and allows readers to place their own datasets in register with them. It also provides a space for the continued buildout of a community-driven atlas-based spatial model of rat hypothalamic chemoarchitecture, allowing experimental observations from multiple laboratories to be registered to a common spatial framework.

## 1 Introduction

For the past century, the laboratory rat has been a mainstay model organism to understand the neural bases of motivated behaviors. But it is only recently that gene-editing tools (D. Li et al., 2013; W. Li et al., 2013) using viral vector delivery have been used in rats to manipulate neurotransmitter systems to better understand their various functions (Homberg et al., 2017; Ma et al., 2017; Rosas-Vidal et al., 2018; Bäck et al., 2019; Neff, 2019; Zallar et al., 2019; Sun et al., 2020; Iwasaki et al., 2023; Yu et al., 2024). Despite such advances, targeting viral constructs to natively expressed chemical transmitter systems in the rat brain remains hampered by a lack of reliable maps of such systems at high spatial resolution. Moreover, there has been a dearth of tools to map and measure the densities of spatial patterns of these systems from multiple experimental subjects. In this report, we describe *Chemopleth 1.0*, a downloadable database of brain atlas maps, with targetable stereotaxic coordinates, for five neurochemical systems in the rat hypothalamus and zona incerta: vasopressin, neuronal nitric oxide synthase (nNOS), hypocretin 1/orexin A (H_1_/ O_A_), melanin concentrating hormone (MCH), and alpha melanocyte stimulating hormone (αMSH). These systems are of great translational significance for understanding feeding control, body weight regulation, fluid balance, arousal and vascular regulation. Two sets of scientific developments – one, a set of careful labeling experiments, and the other, a set of rigorous technical innovations – form the basis of this database.

First, experiments described over twenty-five years ago (Broberger et al., 1998; Elias et al., 1998), identified spatial interactions between neuropeptide-expressing cell populations in the medial hypothalamus (e.g., αMSH) and lateral hypothalamus (e.g., H_1_/O_A_, MCH). These seminal findings, which included mRNA-labeled cell body distributions and select photographs of cellular interactions between some of these populations, remain a high-water mark for hypothalamic chemoarchitectural studies, providing critical structural evidence linking first-order leptin-sensitive neuronal populations to downstream, second-order communication nodes (Sawchenko, 1998; Elmquist et al., 1999). Second, the introduction by Swanson of digital atlas templates in the form of vector-formatted graphics files (Swanson, 1993) as a companion software package to his *Brain Maps* rat brain atlas (Swanson, 1992), marked a quiet but profound turning point for neuroscientists seeking to create electronically formatted maps of neuroanatomical data they obtained from their own experiments for this model organism (reviewed in Swanson, 2001). One of the groups cited above (Broberger et al., 1998) used the *Brain Maps* digital atlas templates to map the distributions of cell bodies for medial and lateral hypothalamic neuronal populations, and additional groups have since provided cell body maps of these populations in more recent versions of the Brain Maps reference space (Swanson et al., 2005; Hahn, 2010).

The *Chemopleth 1.0* database (Navarro et al., 2025; https://doi.org/10.5281/ <u>zenodo.15788189</u>), detailed in the present report, extends these developments – identifying hypothalamic cell groups and mapping their distributions to the *Brain Maps* atlas – in several important ways to better facilitate, for contemporary usage, the precise targeting of these systems and their anatomical registration with other data. First, unlike previous maps, the immunoreactive protein (i.e., immunohistochemical) maps in this database chart not only the cell bodies but also the axons and, in some cases, the terminal fields specific to these neurochemical systems in a standardized format. This is important for those interested in delivering retrogradely transportable vectors into axonal terminal fields (e.g., retro-AAV vectors) into the rat brain, the locations for which are not provided by mRNA-based maps. Second, the cell bodies and axons charted in our database are mapped with respect to stereotaxic space, coordinates for which should be valuable to any experimentalist seeking to have a basic starting point to target vectors in the rat brain. At the time of this writing, such stereotaxically guided information is often lacking in 2-D models at the spatial resolutions mapped here, let alone 3-D models of rodent hypothalamus produced by light sheet microscopy or other methods. Third, we provide computational tools that permit visualization and analysis of spatial information for each system from more than one subject and in more than one graphical representation. This has been carried out to showcase the extent of variability for the distribution patterns observed among several subjects, and to identify for the community the denser areas for each neurochemical system that can be experimentally targeted in stereotaxic space. Fourth, we present a new take on an older conceptual framework – grid-based annotation – that refines the existing stereotaxic coordinate system of the rat brain atlas and leverages its use not only for mapping but also for the gentle entry of outside datasets from the published literature into the database for the community to extend its contents.

Importantly, this study highlights the utility of the *Brain Maps* reference atlas as a *data repository*, taking advantage of the Adobe® Illustrator® (<u>Ai</u>) vector graphics workspace for the current *Brain Maps 4.0* digital atlas templates (“BM4.0”; Swanson, 2018; <u>RRID:SCR_017314</u>), and the ability of users to use the free vector graphics editor, Inkscape (www.inkscape.org) as another way to access this database. The foundation of the *Chemopleth 1.0* database available in the Zenodo data repository (Navarro et al., 2025; https://doi.org/10.5281/zenodo.15788189) consists of downloadable <u>Ai</u>- and Inkscape-compatible Scalable Vector Graphics (SVG) files containing atlas templates for eight atlas levels of what have been formally designated as the *hypothalamus (Kuhlenbeck, 1927)* and *zona incerta (>1840)* (see *Section 2.1*) that form base maps in BM4.0. Building upon these base maps, we provide the following tools/resources:

1. *data layers* above the base map containing the mapped distributions of five chemically identified neuronal populations (both their cell bodies and their labeled neurites). These populations include the vasopressin gene product, copeptin; nNOS (<u>EC 1.14.13.39</u>), H_1_/O_A_, MCH, and αMSH;
2. *choropleth maps*, which sort the relative abundance of distributions for these five neurochemical systems, in 3–7 subjects, by brain region;
3. *isopleth maps*, which sort the relative abundance of distributions for the same neurochemical systems, in 3–7 subjects, independent of brain region; and
4. *a grid-based coordinate system*, including the introduction of an ephemeris table organization for the distributions of these neuronal populations within this grid.

The <u>Ai</u> / Inkscape workspaces allow for the interactive visualization of data layer overlays over each base map. Each data layer, which can be accessed using the *Layers* panel in <u>Ai</u> (*Layers and Objects* panel in Inkscape), contains its own set of spatial data visualizable in relation to data present in other layers “in front of/above” or “in back of/under” the layer being examined.

After describing how the *Chemopleth 1.0* database and its tools were built, we describe the mapping innovations it contains and how to navigate the digital overlays within it. We then discuss the need for these tools, highlight the contents of the database in relation to previous visualization efforts, and discuss the design features of the database, including our conceptual framework for grid-based annotation. Because we present visualization tools (downloadable vector-formatted maps with digital overlays; open-source code for spatial averaging) for a structural model (vector-graphics atlas) of a complex object in nature (brain), our narrative draws from an eclectic set of disciplines (cartography, geosciences, computing, the humanities and arts) – all of which include scholarly and artistic efforts to visualize, register, and annotate complex information in time and space.

## 2 Materials and Methods

### 2.1 : Naming conventions

*Chemopleth 1.0* contains data populating brain regions named according to lexical guidelines set forth in Swanson (2015), as applied to the rat brain in *Brain Maps 4.0*, an open-access rat brain atlas (Swanson, 2018; <u>RRID:SCR_017314</u>), and includes standard terms. Each standard term is set in italics and includes the named neuroanatomical structure and the associated (author, date) citation that first uses the term as defined, e.g., “*lateral hypothalamic area (Nissl, 1913)*”. For those terms where assignment of priority was not possible, they are assigned the annotation “*(>1840)*”, i.e., “defined sometime after the year 1840”, a year which roughly marks the introduction of cell theory in biology, e.g., “*dorsomedial hypothalamic nucleus (>1840)*”. Standard terms for *gray matter regions (Swanson & Bota, 2010)* can be found in “Table C: Rat CNS Gray Matter Regions 4.0 (Topographical Histological Groupings)” from the Supporting Information available online for Swanson (2018), which is based on the scientific literature and cytoarchitectonic features of Nissl-stained tissue sections. Similarly, *white matter tracts (Bell & Bell, 1826)* are listed in “Table D: Rat CNS White Matter Tracts 4.0 (Topographic Histological Groupings)” from the same Supporting Information, e.g., “*hypothalamic postcommissural fornix (Swanson, 2015)*”. Importantly, any citation included within the standard term is included in the list of cited references in this study. A list of the standard terms and the abbreviations used in this study is provided in the Abbreviations section.

### 2.2 : Workflow to create Chemopleth 1.0

#### 2.2.1 : Data sources

In principle, as with other transformation workflows (e.g., see Oguchi et al., 2011; Khan et al., 2018b) a variety of data sources, analog or digital, could potentially serve as input data to our database with the appropriate transformations. To populate *Chemopleth 1.0*, we assembled data types from a variety of sources; these types are numbered 1–5 in **Figure 1**. The most basic input data consisted of single-subject maps drawn by our laboratory in the commercially available workspace, Adobe® Illustrator® (<u>Ai</u>), on individual data layers over a base map. These were produced within the vector graphics application itself and registered to the *Brain Maps 4.0* base map (Swanson, 2018) which is the bottom layer (“base maps”; **Fig. 1**) making up the database (**Inputs 1 + 2 + 3, Fig. 1**). A second, more derived set of data populating our database is in the form of “heat maps” of spatially averaged data that are created by first exporting several individual subject maps (of a single neurochemical system at a single atlas level) as Javascript-converted Portable Network Graphics (PNG) raster-graphics files or a Scalable Vector Graphics (SVG) vector-graphics files. These are then processed by a Python script to create the heat maps using a Gaussian filtering form of kernel density estimation, after which these maps are re-imported into the database (**Inputs 4 and 5, Fig. 1**). Note that we used the <u>Ai</u> workspace to create these files, but they can, in principle, be generated within the free vector graphics editor, Inkscape (<u>www.</u> inkscape.org; version 1.4.2), which we used to open/test the files in that system at the time of this writing. Guidance for downloading, installing and using Inkscape can be found online at the <u>Inkscape Beginner’s Guide</u>. All procedures described below were performed within or in relation to the use of <u>Ai</u> workspace, and hence will be discussed mainly in that context, but readers using Inkscape should be able to utilize the database just as readily.

**Figure 1.**
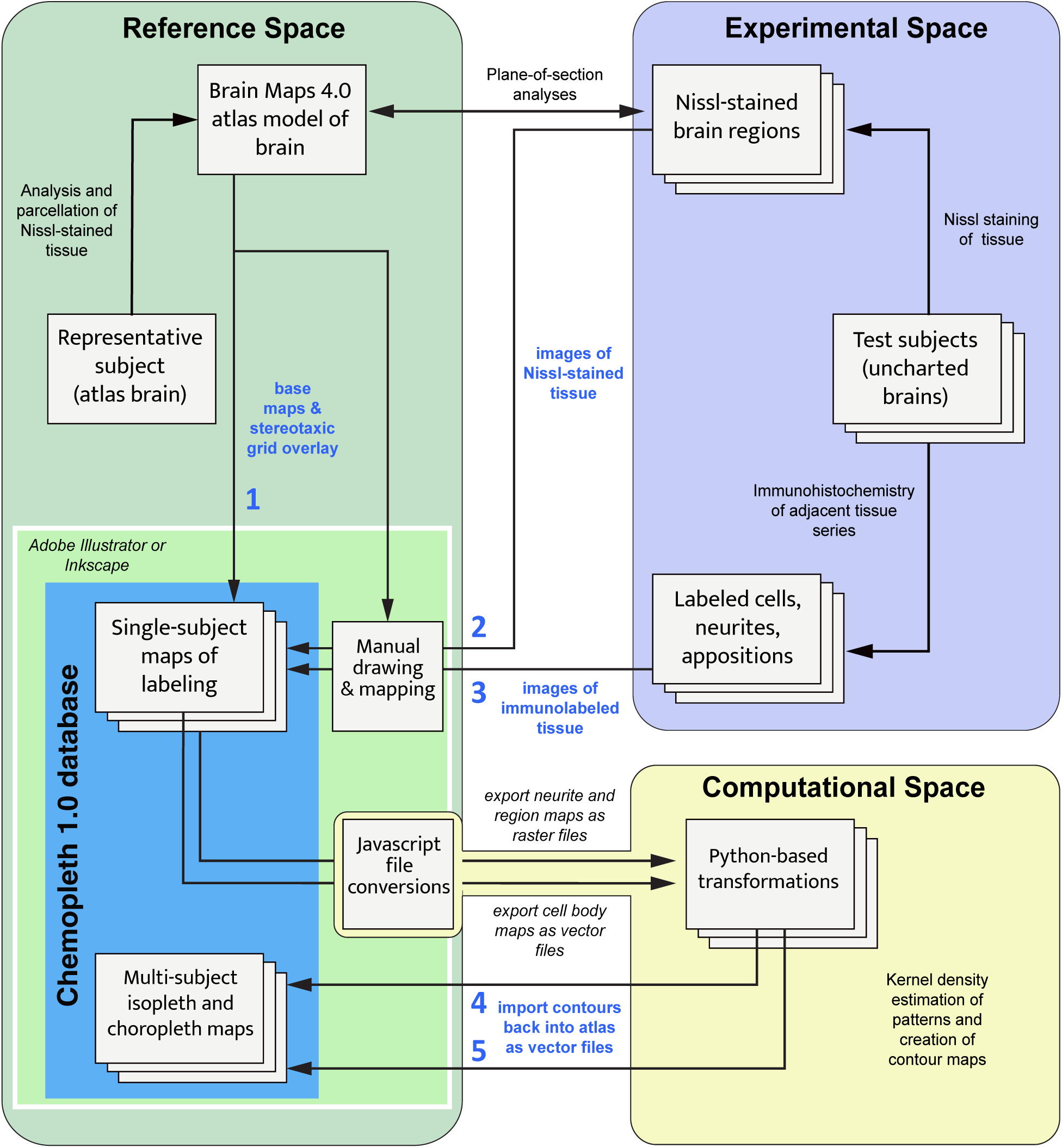
Workflow to generate content for the *Chemopleth 1.0* database. Five input sources (numbered **1–5** in bold within the schema above) feed data into the database (dark blue box): **(1)** base maps and stereotaxic grids from the *Brain Maps 4.0* rat brain atlas in *Reference Space*, for which **(2)** images of Nissl-stained tissues from experimental brains (*Experimental Space*) are used to manually draw critical context-of-plane boundaries for brain regions, that, in turn, allow us to manually map **(3)** immunoreactive cells and neurites from adjacent series of tissues from these subjects onto the base maps. These single-subject maps can collectively be exported, after appropriate Javascript-enabled file conversions, into a Python-based *Computational Space*. Within that space, a custom program allows for their spatial averaging using kernel density estimation and their import back into *Chemopleth 1.0* (Steps **4** and **5**) as heat maps of multi-subject data on a single base map. These heat maps can either be *choropleths* (based on atlas territories), or *isopleths* (based on distributions independent of the underlying base map). See text for details.

#### 2.2.2 : Workflow details

Early workflow attempts are noted in our *Supplemental Methods* (see Supplementary Materials accompanying this article). Here, we describe the workflow we settled upon to generate the data and populate the database. It operates within three major conceptual spaces: *Reference*, *Experimental*, and *Computational* (**Figure 1**). The *Reference Space* includes the *Chemopleth 1.0* database, which provides new content for the rat brain atlas reference, the *Brain Maps 4.0* (BM4.0) rat brain atlas (Swanson, 2018; <u>RRID:SCR_017314</u>). Included in this *Reference Space*, therefore, is the actual animal subject which forms the basis of the BM4.0 atlas, an adult male 315-g Sprague Dawley rat (“representative subject”), along with its Nissl-stained brain tissue which was studied by Swanson to produce cytoarchitectonic boundaries for all major cell groups of the rat brain (“atlas brain”); these were drawn as vector objects to form atlas maps (“Brain Maps 4.0 atlas model of the brain”). The BM4.0 atlas maps, in turn, supply both the *base maps* and *stereotaxic grid* overlay for the *Chemopleth 1.0* database (**Input 1** in **Fig. 1**).

The *Experimental Space*, in contrast, contains the experimental subjects for which we wanted to map chemoarchitectural labeling within the BM4.0 atlas, and which would form the basis of *Chemopleth 1.0*. In this space, we sectioned brain samples, collecting 30 μm-thick tissue sections into adjacent series of fixed-interval tissue sets, one of which was stained with thionine to generate a Nissl-stained reference series, and another stained for a neuropeptide or neural enzyme of interest using immunohistochemistry (detailed in Navarro, 2020; Peru, 2020). Photomicrographs of the Nissl-stained series displayed labeling patterns that revealed the overall cellularity of the tissue, that allowed us to draw boundaries around the stained cell clusters (parcellating them into nuclei and areas) or around white-matter tracts. These drawn boundaries, in turn, were overlaid onto the images of tissue sections from the adjacent immunostained tissue series for which (aside from the immunolabeled material) overall cellularity was not visible. These sections were near enough in the brain to the Nissl-stained tissue series to lawfully permit the transfer of its drawn boundaries.

In **Figure 1**, we have noted these processes in the *Experimental Space* with the photomicrographs (digital images) of the Nissl-stained tissues and immunoreactive cells and neurites serving as **Inputs 2** and **3**, respectively, which were each used to draw boundaries and labeling patterns onto BM4.0 base maps to assemble the single-subject maps in the *Choropleth 1.0* database within the *Reference Space*. Thus, the single-subject maps of chemoarchitecture in *Chemopleth 1.0* were a product of: (**Input 1**) BM4.0 digital-format base maps, together with their stereotaxic grid; overlaid with drawn Nissl-delimited boundaries from images of the Nissl-stained tissue (**Input 2**) to guide the mapper in taking their drawings from images of labeling patterns of immunoreactive cells and neurites for the neurochemical systems we examined (**Input 3**) over the right locations on the base map. The biological significance of the immunohistochemical staining patterns in the native tissue, together with their photomicrographic documentation, will be discussed in separate articles elsewhere.

Finally, we sought to develop cartographic methods that allow us to visualize immunoreactivity patterns in the same atlas level for more than one subject, so that consensus locations on the map could be identified across subjects where labeling was observed. To achieve this goal, we wrote instructions in Javascript to convert cell and fiber maps from single subjects into file formats that could be exported out of the database and processed in a Python-based (A. Arnal, Mar 2021) *Computational Space*, to then re-enter the *Reference Space* as multi-subject “heat maps” showcasing the concordance of distributions of immunoreactivity for any given neurochemical we labeled. Two types of “heat maps” were generated (**Inputs 4 and 5** in **Fig. 1**).

The first type of heat map was a *choropleth map* (Dixon, 1972), which scores the density of labeling as a color-coded gradient over bounded areas. In our study, there were two types of choropleth map: (1) a *brain-region choropleth map*, which is a base map of defined brain regions with vector-object boundaries (these boundaries first had to be ‘closed’ to form discrete polygons; see *Section 2.4.2.1*); and 2) a *grid-region choropleth map,* based on rectilinear boundaries formed by the stereotaxic grid overlay in the digital BM4.0 atlas.

The second type of heat map was an *isopleth map* (Mackay, 1951; Schmid & MacCannell, 1956; Barnes, 1978), which is not a product based on binned densities over cytoarchitectonically-bounded regions on the base map or the rectilinear zones of the overlying stereotaxic grid. Instead, the isopleth map is a computationally produced pattern based on kernel density estimation, which allows for distributions that are not bounded by or quantified to fit the cytoarchitectonic regions of the base map underneath them, but reflect, by color gradients, the relative abundances of the distributed label in relation to the total distribution densities across atlas levels.

### 2.3 : Tissue processing, parcellations, alignment, mapping, and semi-quantitative analysis

These procedures, which were required for us to produce atlas maps of the chemoarchitecture, will be described in detail elsewhere, and have been described thus far within our student dissertations forming the basis of this project (Navarro, 2020; Peru, 2020; also see Wells, 2017 and pp. 6–9 of Arnal, 2022). Below, we describe our spatial analysis of these maps to produce our final dataset that forms the *Chemopleth 1.0* database. We also show how to navigate the dataset and import/register user-generated data.

### 2.4 : Preparing the dataset for spatial analysis

Our maps of immunoreactive perikarya and neurites, present within the <u>Ai</u> vector graphics workspace as digital overlays, were spatially analyzed outside of this workspace (*Computational Space* in **Fig. 1**) to produce novel data visualizations and tabulated measurements using Python programming language (Van Rossum, 1993; see Van Rossum (2009–2018) for a history of Python; we used Python 3.0 (Van Rossum, 2009) for this effort). To prepare the data for use in Python, scripts were written in JavaScript (JS) programming language (Wirfs-Brock & Eich, 2020) that could be accessed and run within the <u>Ai</u> workspace to export the maps into usable formats. The JS scripts, *<u>Swanson Data Export.js</u> and <u>Swanson Shape Export.js</u>*, are available in <u>GitHub</u>, along with Python scripts for generating isopleth and choropleth maps and counts.

#### 2.4.1 : Exporting data

Individual data layers from the BM4.0 dataset were exported from the template files within the <u>Ai</u> workspace by running the JS script, *<u>Swanson Data Export.js</u>*. Perikarya or appositions mapped on the electronic atlas templates were exported as SVG files. Each SVG file contained the mapped antigen or apposition from one animal on one level of the atlas and the <u>Ai</u> artboard of the atlas level, which served as the spatial reference frame across files.

Neurite maps on the atlas were exported as PNG files. Unlike the perikarya or apposition maps that were represented by coordinates, neurite paths were represented by raster images, with discrete bins or pixels marking the neurite trajectory. The artboard also served as a spatial reference frame across these files, and the stroke width of all paths was set to approximately 5 µm before exporting.

*Rationale for our export methods*. To prevent us from being limited by the size of the circle glyphs in vector space used to mark the perikarya or appositions, we exported the SVG vector format of the map rather than the raster form of it. Once it was imported as an SVG into the Python program, a raster was then generated internally to represent those cells or appositions, which was then computed upon to generate counts and/or to perform Gaussian blurring. For neurite maps, the entry into Python of the PNG (raster) format obviated the need for any further processing to perform the same analysis.

#### 2.4.2 : Exporting regions of the BM4.0 base map for choropleth analysis

Brain region and stereotaxic grid boundaries were exported (*Swanson Shape Export.js*) to compute region-dependent measurements for each atlas level to produce choropleth maps (as described in *Section 2.5.3*), where densities are delimited by *brain region* or by *stereotaxic grid region*, respectively.

##### 2.4.2.1 : Brain-region choropleth maps

For each *brain-region choropleth map*, some regional boundaries in *Brain Maps 4.0* are only visually implied and had first to be fully closed to enable proper data binning by the Python programs. Therefore, before exporting them, we converted these visually implied regions to closed polygons in <u>Ai</u> **(Fig. 2)**. We found that the easiest way to obtain these polygons without manually redrawing boundaries was to use the *LivePaint* feature in <u>Ai</u>, which interpolates between existing paths that are close to each other. The feature takes a user-defined “gap” parameter to automatically close paths that may not be fully intersecting or touching. After using this feature, we cleaned up the final regions by merging polygons that belonged to a single region such that the number of polygons equaled the number of brain regions. Then, after closing visually implied regions, we addressed the brain regions in the atlas that shared a continuous space with other regions, such as the *internuclear hypothalamic area (Swanson, 2004)* (I). For these cases, our team members (AA, VIN, SB, AMK) conferred to draw new boundaries provisionally. In this manner, all regions had discrete boundaries, which enabled region-dependent quantification of perikarya and neurite profiles. For *Chemopleth 1.0*, these boundaries are a necessary starting point for our analyses, but we anticipate that the boundary assignments will change as new data about the regions come to light or if further discussion warrants a redrawing of the boundaries based on new information. Readers need to be aware that any boundaries they wish to redraw would likely necessitate a re-analysis of choropleth scores, since such scores are computed over the total area occupied by each polygon.

**Figure 2.**
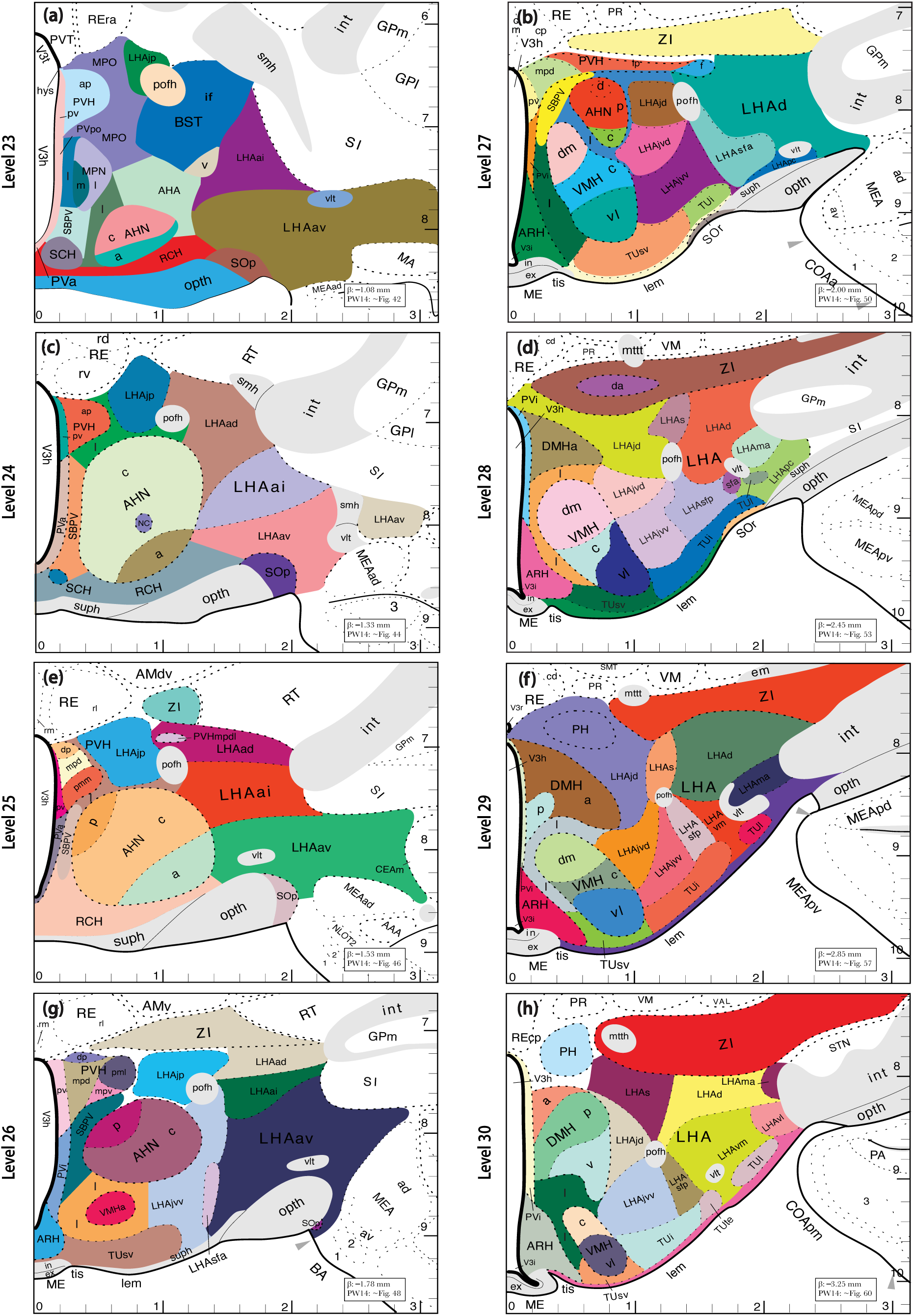
Summary of bounded regions provisionally created for the *Chemopleth 1.0* database within the *Brain Maps 4.0* atlas templates (Swanson, 2018) for levels 23–30. Regions with visually apparent cytoarchitectonic boundaries were provisionally assigned closed boundaries to allow us to compute across them to produce areal densities for choropleth mapping. See text for details. The boxes in the lower right of each level show the anteroposterior coordinate, in millimeters, with respect to the bregma skull suture (labeled ‘β’) which correspond to the position of that plane of section. Below each coordinate is the figure number in the Paxinos & Watson (2014) rat brain stereotaxic atlas (labeled ‘PW14’) that the atlas level most closely corresponds to in the anteroposterior position (as estimated by Khan et al., (2018b) by alignment of the atlases along the anteroposterior axis). Please consult **Table 1** for an explanation of the abbreviations for these regions.

##### 2.4.2.2 : Grid-region choropleth maps

We also generated *grid-region choropleth maps*, which computed the densities of cells, fibers or appositions based on bounded *grid regions* from the stereotaxic grid embedded within the atlas templates. The *grid regions* were just that portion of the stereotaxic grid that overlaid the areas mapped for this study (hypothalamus and zona incerta) and were collected into a separate layer (called “gridRegions”, see Results). The grid-region counts were used to produce our *FMRS* readouts (see *Sections 2.5.3 & 2.6.2* of the Methods and *Section 3.5.4* of the Results for details).

### 2.5 : Spatial analysis of data

#### 2.5.1 : Importing data into Python for further computational analysis

The raster approach described in *Section 2.4.1* allows binning the perikaryon/apposition centroid points and neurite profiles to a uniform grid of pixels (“control areas”) to facilitate cell and neurite counts for choropleth mapping. A useful reference to guide readers on the general principles of setting control areas is provided by Barnes (1978). The approach also allows for density estimation of perikaryon, apposition, and neurite profiles by Gaussian blurring to produce isopleth maps.

#### 2.5.2 : Generating isopleth maps for mapped immunoreactive perikarya, appositions, and neurites

To visualize spatial trends of immunoreactivity in a manner that considers subtle differences in the exact atlas map placements of filled circles for perikarya or appositions that would statistically occur because of the mapper, we decided to represent each instance of an immunoreactive perikaryon/apposition and neurite segment as a normal distribution curve in two dimensions. The normal curve has a long history of describing error distributions, dating back to its mathematical formulation a few decades before the emergence of cell theory in the mid-19th century (Adrain, 1808; Gauss, 1809/1857; see Stahl, 2006). Suitably, the normal curve is an appropriate representation of a mapped perikaryon, apposition, or neurite segment since it embodies the distribution of errors, with the peak of the curve assigned as the centroid point of the plotted perikaryon or apposition and the possible alternative positions on the plot assumed to be within a normal distribution. Specifically, the sigma value (σ), which determines the spread of the normal curve, can be considered a proxy for errors incurred when mapping a particular perikaryon, apposition, or neurite to an atlas template. Based on sampling of various hypothalamic regions and measuring the diameters of perikarya (10–15 perikarya/region), we found the average diameter of perikarya to usually range between 10–20 μm. We therefore set σ to ∼14 μm, which would place any perikaryon’s mapped circle centroid point at ∼28 μm from the first standard deviation of the Gaussian curve (third column (10 pixels) in **Fig. 3**), thus conservatively blurring the location information for 1–3 mapped perikarya. Now, if a perikaryon is ∼28 μm in diameter, this means that a region with a diameter of ∼28 μm can physically only contain information about one perikaryon’s position. Put another way, the only time a region of 28 µm can contain information about more than one perikaryon is if there is an overlap of perikaryon positions from multiple experimental subjects. We found the probability value, or the height of the curve, at the first standard deviation and used multiples of that height as thresholds to draw contours. This effectively gave us a contour for regions where densities were similar. Of course, depending on the needs of the mapper and the sizes of the cells to be mapped, one could adjust the sigma value to reflect the dimensions of other perikarya. We suggest that readers try different values based on their region of interest, the size of the perikarya in that region, and the desired granularity of the isopleth map needed. **Fig. 3** provides a view of how the maps change as a function of changing the value for σ.

**Figure 3.**
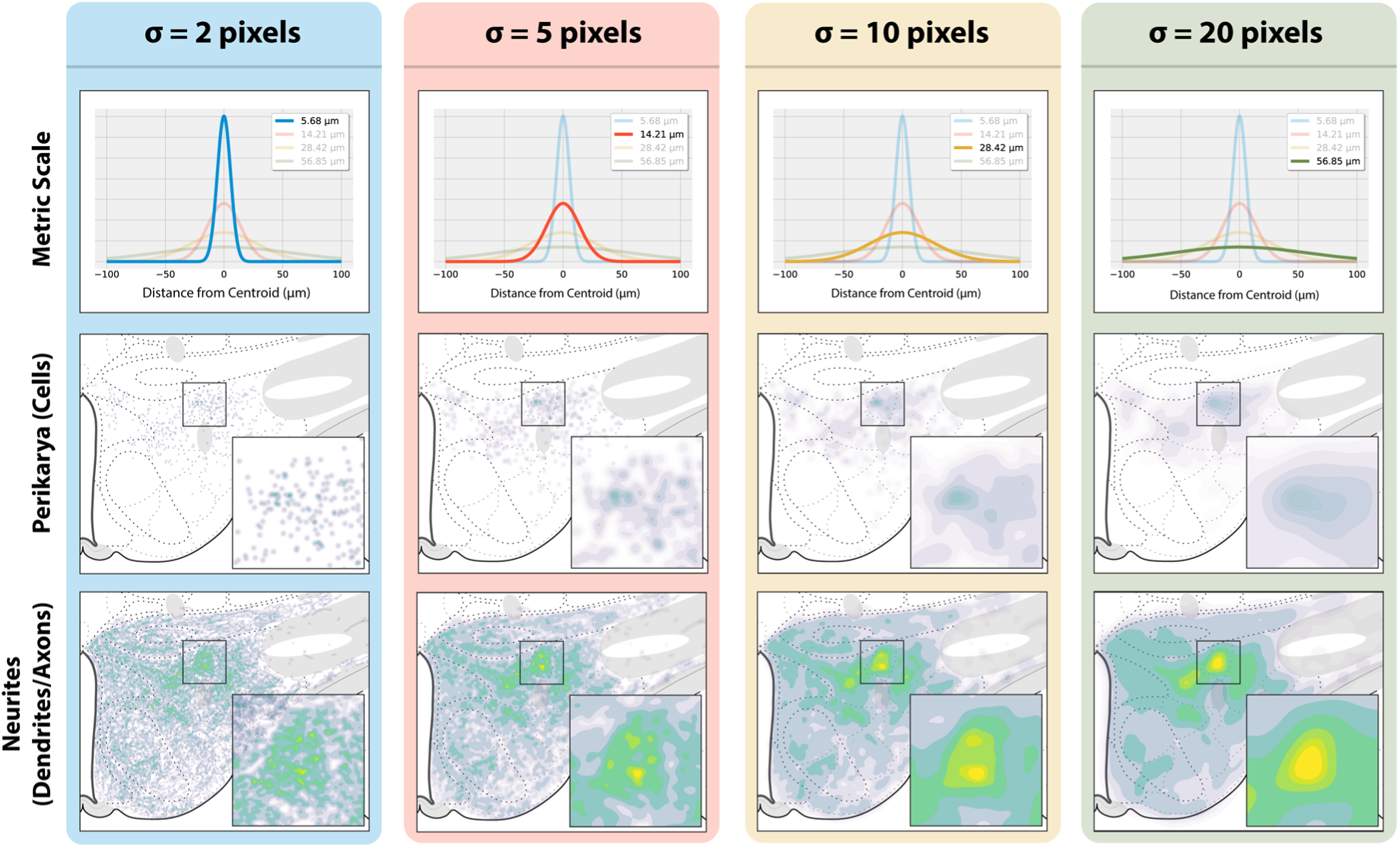
Comparison of kernel density estimation parameters and their resulting isopleth maps. The columns from left to right are sorted by increasing values for sigma (in pixels), which is a value that signifies the spread of the normal distribution curve for any mapped perikaryon. **Top row:** The normal distribution curves resulting from the sigma value assignments, as measured in microns. **Middle row:** Isopleth maps of the mapped perikaryon profiles immunoreactive for hypocretin 1/orexin A (H_1_/O_A_) from multiple subjects averaged together and plotted on the base maps of the *Brain Maps 4.0* rat brain atlas (Swanson, 2018), with the box outlining the enlarged area containing a portion of the *lateral hypothalamic area (Nissl, 1913)*. **Bottom row:** Isopleth maps of the neurite profiles from the same subjects, with the box outlining the same region as for the middle row (cell body) maps. For the *Chemopleth 1.0* database, all isopleth maps were generated with a sigma value of 10 pixels (*third column from the left*). See text for details.

In our implementation, we used the <u>gaussian_filter</u> method of the Python library, <u>SciPy</u> (v1.7.0), which takes as input, 1) the raster image of the mapped distributions from a BM4.0 atlas template and, 2) the σ value, and produces a blurred version of the input image or raster. Mathematically, the blurring occurs after computing a 2-dimensional normal curve of pixel intensities at every pixel position. Moreover, this Gaussian blur method is faster than the traditional stepwise kernel-density estimation method of iterating over all perikarya, appositions and neurite segment centroids and computing the density of every point while progressively adding them together (Gray & Moore, 2003; Virtanen et al., 2020). Once the scripts (cells: *<u>cellIsopleths.py</u>*; appositions: *<u>appositionIsopleths.py</u>*; fibers: *<u>fiberIsopleths.py</u>*) computed the density estimation maps (utilizing the <u>NumPy</u> Python library), the maps were passed to a contouring function (utilizing the Python plotting library, <u>matplotlib</u>, v3.5.1) to draw contours at 10–100% of maximum density in 10% increments. Each script then saved the contour as an SVG file which we then placed back on the map in the <u>Ai</u> workspace **(Fig. 4)**.

**Figure 4.**
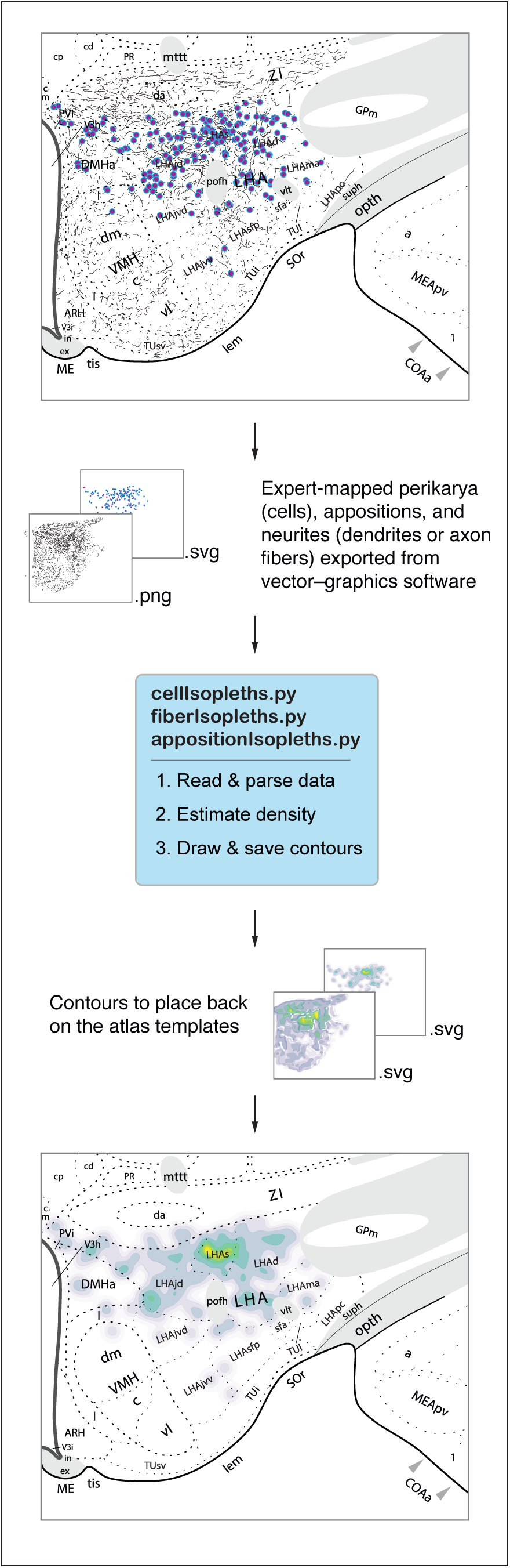
Process of generating isopleth maps of chemoarchitecture from multiple subjects based on individual single-subject maps exported and computed upon by our custom Python workflow. See text for details.

We also extended this process to visualize spatial trends of cell and fiber distributions for a given neurochemical system from multiple cases. To achieve this, we computed each density map, and before passing each of them to the contouring function, we combined the density maps by averaging. Then, for multiple levels, we looped through the final density maps of each level and recorded the minimum and maximum values across all levels. Finally, we generated a contour map for each level. However, this time, we provided the minimum and maximum values to the contour function, allowing our selected color scheme to capture the relative distribution among levels.

#### 2.5.3 Generating choropleth maps for mapped immunoreactive perikarya and neurites

To streamline the process of counting perikarya or estimating neurite density per region, we developed Python scripts that utilize raster brain regions or stereotaxic grid regions to automatically calculate the number of perikarya, appositions, and approximate area of neurites (to produce two types of choropleth map: one type based on brain region and another based on stereotaxic grid region; see *Section 2.4.2*). The data loading process for this method is identical to that used for the isopleth analysis, except that in the case of brain regions, these were first prepared from the BM4.0 atlas for export as described earlier in *Section 2.4.2.1*. Alternatively, grid regions from the stereotaxic grid overlay in BM4.0 were used instead. Once the raster versions of the data were ready, we loaded the region files and iterated through all the brain or grid regions across the corresponding level. For cells, appositions and fibers, the scripts used, respectively, were: *<u>cellChoropleths.py</u>*, *<u>appositionChoropleths.py</u>*, and *<u>fiberChoropleths.py</u>* (all available in <u>GitHub</u>).

At each step within each script, we calculated the number of pixels in each data raster that overlapped with the region raster. For this purpose, our Python scripts for choropleth map generation utilized <u>NumPy</u> (v1.21.0) and <u>matplotlib</u> (v3.5.1). For perikaryon profiles, we tabulated the number of overlapping pixels representing the direct perikaryon profile count and applied a sampling correction (Abercrombie, 1946). For neurite profiles, we tabulated their approximate area by scaling the count of overlapping pixels by the metric conversion factor of approximately 8 × 10^−6^ mm^2^/pixel (i.e., 1 pixel represents that much space in vector space). We then calculated the averages of counts or areas per level across cases of the same marker. Subsequently, we utilized the tabulated information to generate choropleth maps with brain regions or grid regions serving as the bins of each type of choropleth map, respectively. However, instead of simply using the counts of perikarya or the area of neurites per region to color the choropleth regions, we normalized all values by the areas of their respective brain regions or grid regions. The final choropleth maps were saved as raster (PNG) files and then placed on a data layer within the <u>Ai</u> atlas workspace. Next, we iterated across the averaged levels to record the lowest and highest values, which were used to assign colors that captured the relative distribution among levels of a given marker.

### 2.6 : Spatial referencing of profile counts

#### 2.6.1 : Feature-based spatial referencing

As described previously (*see* Fig. 6 of Khan et al., 2018a), the spatial distributions of the data plotted in the maps generated in this study were determined by superimposing the drawn boundaries of *gray matter regions (Swanson & Bota, 2010)* [see *Section 2.1* for naming conventions] of the HY and ZI, determined by cytoarchitectural criteria revealed by adjacent Nissl-stained tissue sections. The cytoarchitectural criteria used in this study are based on those used by Swanson (2004), which draw upon a wide variety of studies for the ZI and each of the HY *gray matter regions (Swanson & Bota, 2010)* presented. In summary, these criteria include the cellular morphology, staining intensity, orientation, packing density and distribution, as revealed by Nissl-stained sections. The tissue range studied in the present report includes that which falls within BM4.0 atlas templates (levels) 23–30, which comprise two of the four major rostrocaudal divisions of the *hypothalamus (Kuhlenbeck, 1927)* (HY) (anterior and tuberal). For each of these levels, the entire mediolateral and dorsoventral extents of the HY were analyzed, and many of these levels also included the *zona incerta (>1840)* (ZI) within the *ventral thalamus (His, 1893)* (TH) (see *Section 3.2*). Cytoarchitectural boundaries of tissue sections were used to identify them as corresponding to one of the atlas levels observed in this study (plane-of-section analysis), which then allowed for the transfer of data from immunohistochemically stained sections to atlas templates. These same cytoarchitectural boundaries were overlaid onto the photomicrographs of immunohistochemically stained sections using <u>Ai</u> software to organize them into transparent layers (digital overlays). Overlays in <u>Ai</u> also allowed for the migration of data from tissue sections to approximate locations of all mapped perikarya and neurites onto atlas templates in this study. We distributed our profile counts across the hierarchical list of brain regions found in Table C: Rat CNS Gray Matter Regions 4.0 (Topographical Histological Groupings) from the Supporting Information available online for Swanson (2018). Counts were made from the mapped glyphs denoting immunoreactive neuronal profiles and tabulated.

#### 2.6.2 : Grid-based spatial referencing

Complementing our feature-based spatial referencing, we reasoned that indexing the quantitative data to an atlas-independent spatial reference system would allow the data to be readily migratable to other atlas reference spaces indexed to the same format, as we have discussed previously (see Section 4.6.2.2 of Khan, 2013; Khan et al., 2018b; Perez et al., 2018). Accordingly, we are proposing a system to tabulate the distributions of our profile counts within a stereotaxic, grid-based spatial referencing system. This was facilitated by Swanson’s inclusion of the stereotaxic coordinate system of Paxinos and Watson (1986) as a data layer in the electronic vector-formatted templates of his atlas. The *FMRS annotation system* presented in the Results (*Section 3.5.4.2*) is essentially a refinement of that existing atlas feature and modifies the existing data layer containing the original atlas stereotaxic grid to include labels designated for each grid region.

### 2.7 : Validation of methods

Several checks were made of our processes for generating the datasets or use of the tools presented in *Chemopleth 1.0*. First, for counting procedures, our automated quantitative analysis was checked against expert, manually annotated counting (e.g., comparisons between the two methods for counting H_1_/O_A_-ir cell bodies). Second, for our isopleth map constructions, we aligned the isopleth generated for a given chemical system with the underlying single-subject maps to see if higher densities in any given subject generally reflected the trends marked by the spatial averaging of the isopleth corresponding to that chemical system **(Fig. 5)**. Finally, though this was not intended to be a validation of our maps, we found that registering published legacy datasets, replotted onto base maps and aligned to our isopleths (*described next*), showed a marked correspondence between datasets of the same mapped neurochemical systems from the pairs of studies (see *Results*).

**Figure 5.**
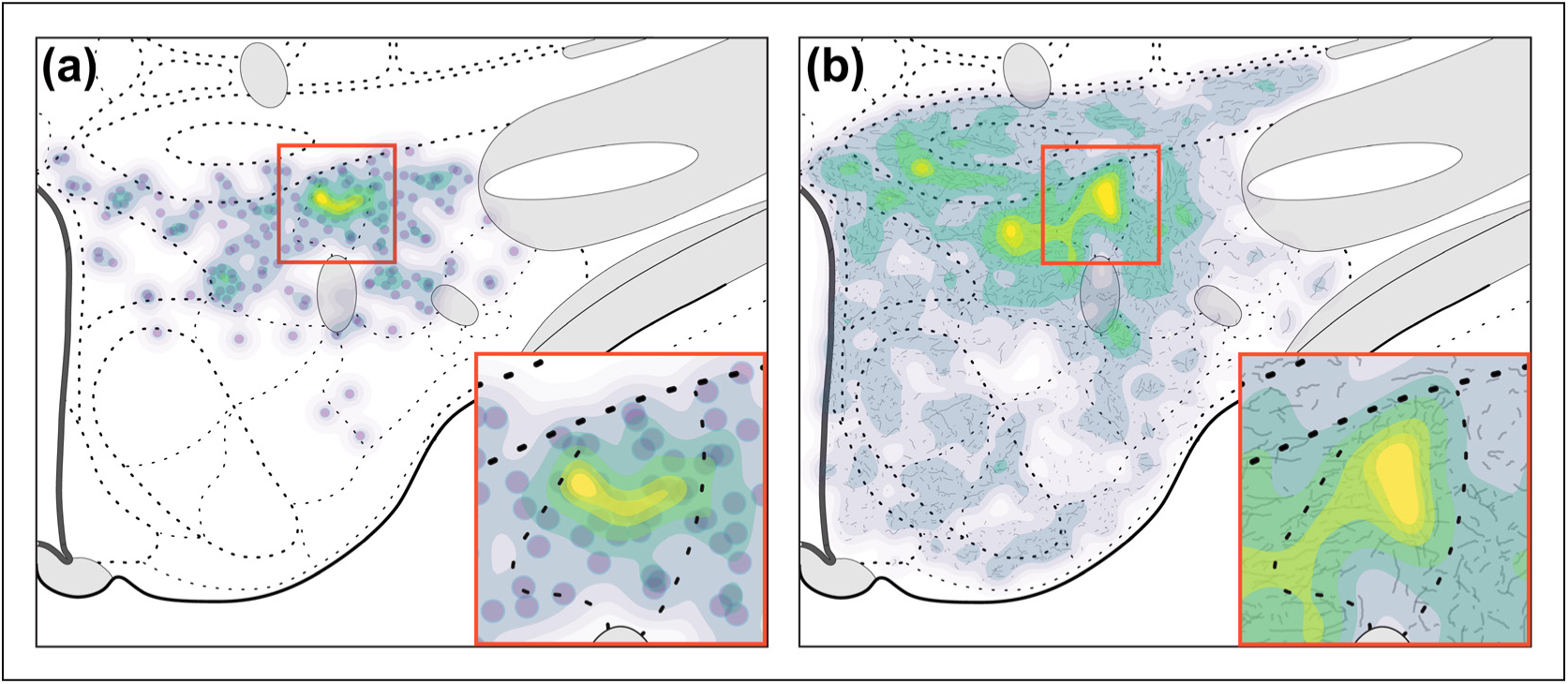
Qualitative validation of isopleth accuracy. Isopleth maps were overlaid over **(a)** perikarya maps or **(b)** neurite maps from single subjects to compare the distribution patterns between the maps. Notice that the brighter areas, which reflect greater average density, tend to be over single-subject maps that also show areas of higher density, thus accurately reflecting those patterns.

**Figure 6.**
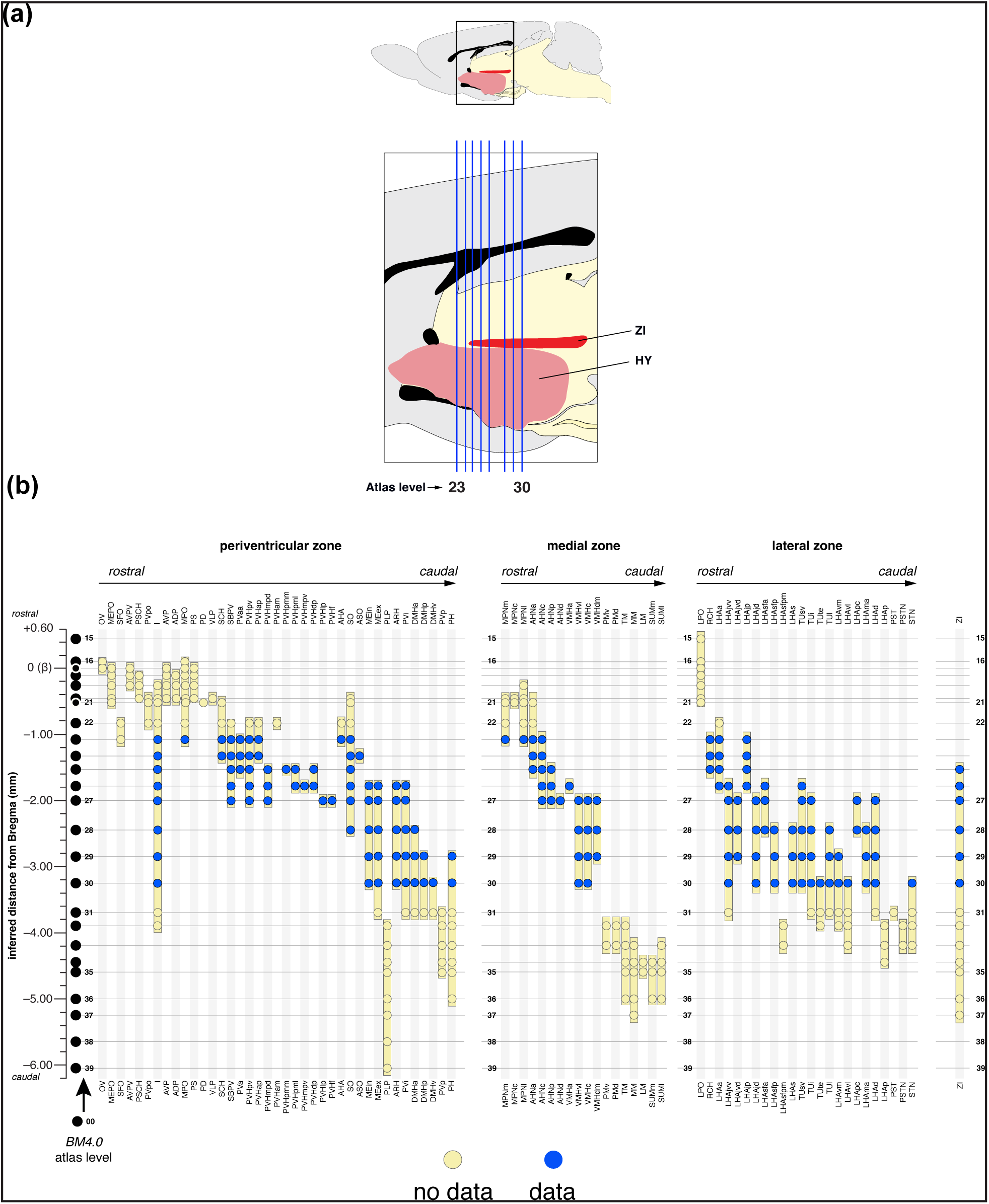
Coverage of *hypothalamus (Kuhlenbeck, 1927)* (HY) and *zona incerta (>1840)* (ZI) within this dataset. (**A**). (*top*) sagittal-plane silhouette of a rat brain from the Swanson (2018) *Brain Maps 4.0* (BM4.0) atlas, with the box outlining the enlarged area containing the approximate extents of the HY and ZI. *Blue lines* indicate the planes of section through both structures representing levels 23–30 in the atlas (designated as S^23–30^). (**B**). A more granular view of the coverage of the HY and ZI in our dataset, with the vertical axes to the left and right marking the bregma coordinate (in mm). *Filled black circles* at the left mark individual atlas levels in BM4.0 with positions calibrated to bregma. The top and bottom axes show, in approximate rostral to caudal order, major brain regions (marked by their atlas abbreviations) that are included in the HY and ZI. From left to right, the clusters of regions are organized by the longitudinally arranged *periventricular*, *medial* and *lateral zones (Nauta & Haymaker, 1969)* of the HY (Swanson, 2018; see Bedont et al., 2015; Xie & Dorsky, 2017), followed by the ZI. *Filled blue circles* indicate those regions in the BM4.0 base maps, at their particular atlas levels, that are populated within *Chemopleth 1.0*. Note that the most rostral (S^15–22^) and caudal (S^31–39^) levels are not populated (*pale yellow circles*) at the time of this writing.

### 2.8 : Migration and registration of legacy datasets to Chemopleth 1.0

To migrate the datasets from Yao et al., (2005) and Kerman et al., (2007), we cropped digital photos of the older maps from the publications and placed each of them as a transparent overlay over the base map of BM4.0 that corresponded to the atlas level we imported. These were then overlaid with an additional transparent layer to mark the locations of the perikarya by placing a glyph over the underlying location where the original was mapped, taking care that the maps were aligned precisely. So re-plotted, the data of the imported study, along with the relevant overlay for our isopleth map of the same chemical system, were prepared as figures for this study in <u>Ai</u>.

## 3. Results

### 3.1 : Chemopleth 1.0 dataset files, source code, documentation, and data management

*Chemopleth 1.0* is accessible from the online data repository, Zenodo, which holds a stable version of the dataset that enables further versioning (Navarro et al., 2025; https://doi.org/10.5281/ <u>zenodo.15788189</u>). The base maps forming *Chemopleth 1.0* are digitally cropped portions of electronic files from the open-access *Brain Maps 4.0* (BM4.0) rat brain atlas (the unpopulated base maps can be found in the Supplementary files accompanying Swanson, 2018; <u>RRID:SCR_017314</u>). The mapped datasets populating these base maps were produced by our laboratory and have been reported in Peru (2020) and Navarro (2020). The original staining patterns, together with their photomicrographic documentation, are being prepared separately for publication. In the present report, the maps of these staining patterns have been prepared for viewing within <u>Ai</u>/Inkscape as eight individual SVG files, each representing a unique atlas level. Each file contains a BM4.0 base map of a specific atlas level and digital overlays of our maps, produced for multiple experimental subjects, for these neurochemical systems. Source code for our workflow is available in <u>GitHub</u>. The source code contains the JavaScript (JS) and Python programs to analyze shapes and data in BM4.0. The JS scripts are used to export data and shapes (brain regions or stereotaxic grid regions) from BM4.0 base maps in Adobe Illustrator (<u>Ai</u>), and the Python scripts are used to generate summaries, choropleth maps, and isopleth maps of the data, described in greater detail in the sections that follow. A *Supplemental Video* also documents how to navigate the files (see Supplementary Materials).

Importantly, *Chemopleth 1.0* complies with ‘FAIR’ community guidelines for ‘Findability, Accessibility, Interoperability, and Reusability’ of datasets to enable them to be read by human- and machine-actionable processes (Wilkinson et al., 2016; Martone, 2024). First, the datasets in *Chemopleth 1.0* (Navarro et al., 2025) are findable in the Zenodo data repository with a unique and persistent digital object identifier (DOI: https://doi.org/10.5281/zenodo.15788189). Moreover, rich metadata are associated with each atlas file using XMP (eXtensible Metadata Platform) labeling technology, an ISO standard (<u>ISO 16684-1:2019</u>) that enables embedding of the metadata within each file itself. In each of our atlas files in <u>Ai</u>, a user can access these metadata by selecting *File* > *File Info*. Metadata, including the DOI, that are associated separately with the Zenodo record are searchable using their search engine. These metadata are also indexed by Zenodo on <u>DataCite</u> servers and are compliant with the non-profit’s metadata schema (https://schema.datacite.org/). Second, *Chemopleth 1.0* metadata are freely available apart from the dataset and harvestable using the <u>OAI-PMH protocol</u> of the Open Archives Initiative. Third, the database is intrinsically interoperable, since the files in the database are released in SVG (<u>version 1.1</u>) format, which can be opened by a variety of open platforms, including Inkscape. Thus, as a database for use within the *Brain Maps* atlas environment, *Chemopleth 1.0* also has features that dovetail with the minimum requirements set forth by Kleven et al., (2023b; 2023c) for atlas environments to meet FAIR standards, which include machine readable digital components in open file formats. Finally, the datasets in *Chemopleth 1.0* are also currently reusable under a <u>Creative Commons CC-BY-NC</u> 4.0 <u>license</u>.

### 3.2 : Summary of database contents

*Chemopleth 1.0* (https://doi.org/10.5281/zenodo.15788189) contains a full dataset for five immunoreactive neuronal populations through a circumscribed portion of the *hypothalamus (Kuhlenbeck, 1927)* (HY) and a portion of the *zona incerta (>1840)* (ZI) within the *ventral thalamus (His, 1893)* (TH) in multiple subjects. The database includes the profile distributions of perikarya (cell bodies) and neurites (axons and/or dendrites) that were observed to be immunoreactive for one of the following macromolecules: (1) the pre-pro-vasopressin gene product, copeptin; (2) neuronal nitric oxide synthase (nNOS; <u>EC 1.14.13.39</u>); (3) hypocretin 1/orexin A (H_1_/O_A_); (4) melanin concentrating hormone (MCH); or (5) alpha-melanocyte-stimulating hormone (αMSH). Within the BM4.0 reference atlas (Swanson, 2018), the HY is represented by 22 coronal-plane atlas levels (abbreviated here as ‘S’ with superscripted level numbers), containing boundaries derived directly from Nissl-stained sections (from 22 companion image plates) in the same atlas. These levels, S^15–36^, are samples of a 5.45 mm-long anteroposterior expanse of tissue located between +0.45 and –5.00 mm from bregma (β) (from a rat inferred to be in the flat-skull position). Similarly, the portion of the TH containing the ZI is represented through a 3.72 mm-long anteroposterior expanse of tissue by 13 atlas levels (S^25–37^) located between –1.53 and –5.25 mm from β.

In this study, we elected to create standardized maps for the distributions of immunoreactive perikarya and neurites through eight levels of BM4.0 (S^23–30^), ranging from –1.08 to –3.25 mm (from β), which represents 39.8% coverage (2.17 mm) of the HY, and 46.2% coverage of the ZI (1.72 mm, between –1.53 and –3.25 mm from β) (**Figure 6**). This selected expanse of tissue falls approximately within that which has been apportioned as the anterior and tuberal regions of the HY (Swanson, 1987; after Clark, 1938; also see Simerly, 2015; Xie & Dorsky, 2017). In the narrative that follows, we use “anterior” to refer to HY represented within S^23–25^ and “tuberal” to refer to HY represented within S^26–30^.

### 3.3 : Major data visualizations in Chemopleth 1.0

The *Chemopleth 1.0* database is a collection of different maps of immunoreactive chemoarchitecture for the rat *hypothalamus (Kuhlenbeck, 1927)* (HY) and *zona incerta (>1840)* (ZI). This chemoarchitecture includes immunoreactive cell bodies (perikarya) and neurites (axons and/or dendrites). We classify axons and/or dendrites as “neurites” and do not distinguish between them for three main reasons. First, the immunoreactivity we observe does not always allow us to distinguish between the two protoplasmic extensions based on morphological assessment. Second, since there are also occasional instances reported in the HY where an axon can extend from a primary dendrite rather than the cell body (Armstrong & Stern, 1997), some of the neurites, as drawn, may be portions of both (hence, the “and/or” designation; also, see Goaillard et al., (2020) for examples of certain hypothalamic peptidergic systems, not studied here, that also have hybrid extensions). Finally, a protoplasmic extension from the cell body immunoreactive for our labeled proteins is not assumed to be an axon, since there is strong evidence for instances of dendritic compartmentalization of neuropeptides specifically in subregions of the HY (e.g., see Ludwig et al., 2002; C. Chen et al., 2019). More generally, there are compelling data emerging that demonstrate a role for local protein synthesis in dendritic compartments (e.g., Wu et al., 2016; Das et al., 2018). There are two main classes of maps, either for a single subject or for multiple subjects.

#### 3.3.1 : Single-subject maps

These maps are of two kinds: 1) a *single-label map* of immunoreactive cells and fibers for a neurochemical system (e.g., nNOS), representing data from the study by Peru (2020); and 2) a *dual-label map,* where two neurochemical systems were stained and mapped in the same tissue section (e.g., nNOS and αMSH), representing data from the study by Navarro (2020). Each type of single-subject map includes separate maps for cell bodies (perikarya) and neurites (axons and/ or dendrites). Thus, for the single-label map, we have separate maps for cell bodies and neurites (two maps). For the dual-label maps involving two neurochemical systems, we have four maps (two for cell bodies and two for neurites). Additionally, we went further with the dual-label maps as described in Navarro (2020) and added an additional map which marks locations where putative synaptic appositions between the two neurochemical systems were checked under bright-field illumination at high power. **Figure 7a** provides an example of *several single-label, single-subject maps* (each for MCH-immunoreactive neurites) visualized together on a base map (these single-subject maps were superimposed on a single base map, with each map on its own transparent data layer; read more about data layers in *Section 4.4*).

**Figure 7:**
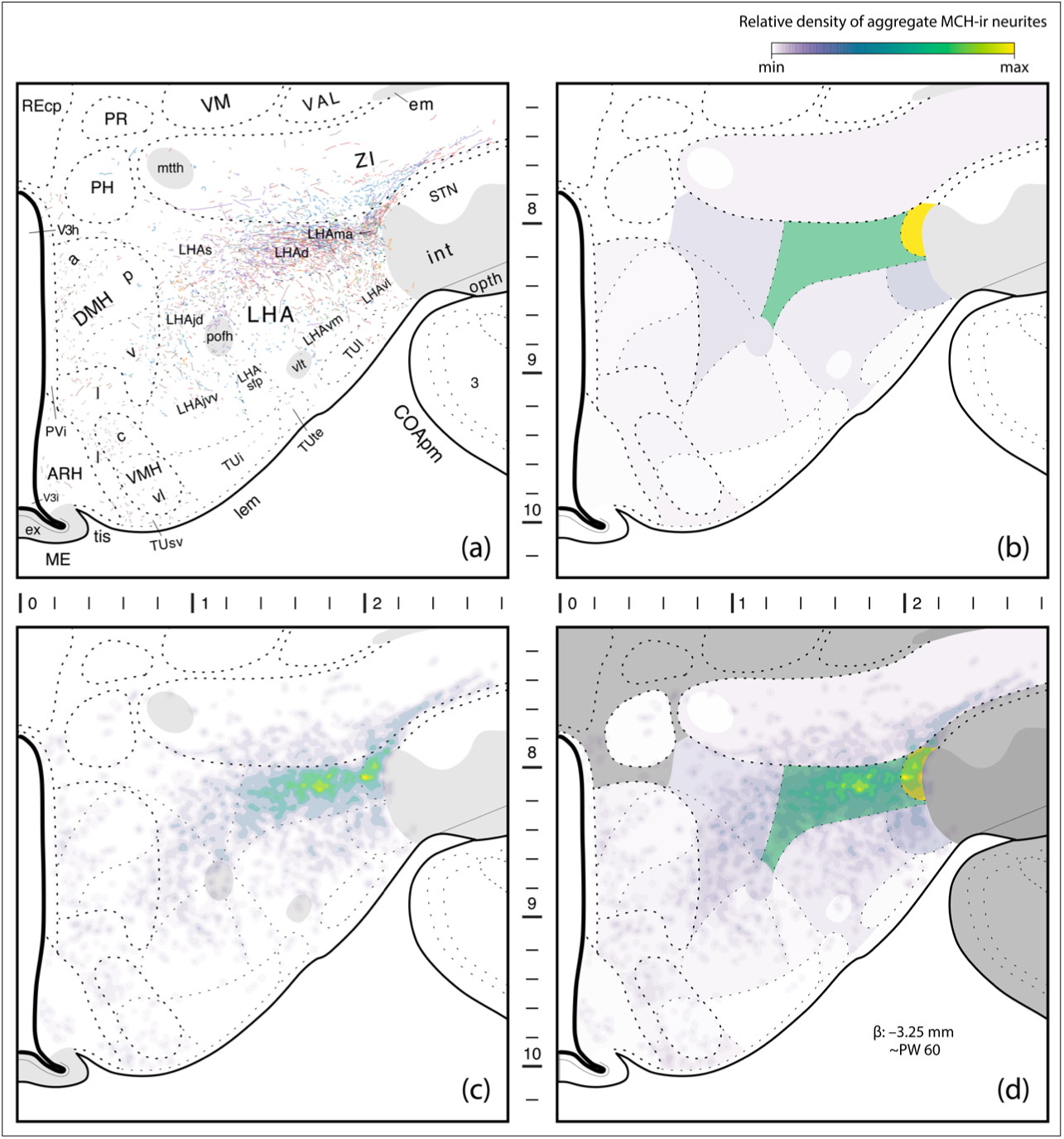
Examples of map types in the *Chemopleth 1.0* database. These maps are for neurites (axons/dendrites) immunoreactive for melanin-concentrating hormone (MCH). Single-subject distributions are superimposed from multiple animals onto a base map to produce a profile of immunoreactive populations in **(a)**, which are scored for areal density within cytoarchitectonically defined territories of the *Brain Maps 4.0* base map to produce choropleth maps **(b)**. The distributions shown in **(a)** are also shown as isopleth maps **(c)**, which are consensus distributions based on kernel density estimation of local spatial conditions of individual distributions, independent of the underlying boundaries of the base map. These two types of “heat map” are superimposed in **(d)** to illustrate how the “hottest” territories scored in the choropleth map (LHAd, LHAma) overlap with those in the isopleth map. The color bar indicates that the scores reflect relative density of the neurites and the scale in between the panels marks inferred stereotaxic coordinates, in millimeters.

#### 3.3.2 : Multi-subject maps

These maps are also of two main kinds. First, *choropleth maps* are area-delimited “heat maps” providing color-coded magnitudes for distribution densities of cell bodies or neurites for each neurochemical system we investigated. **Fig. 7b** shows an example of how multiple single-subject MCH-immunoreactive neurite representations shown in **Fig. 7a** become represented as areal densities across subjects. Each atlas region on the base map is coded by a different color depending on the magnitude of areal density of the immunoreactive fibers within that bounded region. These regions were first prepared for choropleth analysis as described in *Section 2.4.2* and then the analysis was performed on these regions as described in *Section 2.5.2*. Thus, choropleth maps provide heat map assignments confined by cytoarchitectonic region and divided over the areal limit of that region. Another type of choropleth map divides the signal over the areal limit of *stereotaxic grid regions* rather than brain regions (this is described in *Section 2.5.3*).

Second, a separate type of multi-subject representation within *Chemopleth 1.0* is the *isopleth map*, which is a heat map providing color-coded locations of sparse versus dense distributions of cell bodies or neurites *independent* of the underlying boundaries for regions on the base map (i.e., the density is *not* divided over the area of a specific bounded brain region). An example of an isopleth map of MCH-immunoreactive neurites is shown in **Fig. 7c**. This map is a collection of contours which mark the *average densities* of perikaryon or neurite locations on the base map across multiple subjects. Areas that are color-coded in yellow in the figure are denser and reflect greater spatial consensus of the immunoreactive elements across all experimental subjects; those areas coded lighter and duller in shading are areas of lower density, where the presence of immunoreactive elements was found in only one or a few experimental subjects. While the overall isopleth map reflects the *consensus* of mapped immunoreactive elements from multiple subjects, it must be borne in mind that these maps should not be considered as “probabilistic” maps, since the sample sizes are low and therefore the relative contributions of any single subject’s distributions to skew the overall distribution patterns is greater when samples are fewer in number (in our study, n=3–7 subjects).

Keeping this caveat in mind, and consistent with the appearance of a bright region on the choropleth map shown in **Fig. 7b**, the isopleth map in **Fig. 7c** reflects bright or more saturated (i.e., denser) consensus distributions on the base map that, when superimposed on the choropleth map, agree with it (**Fig. 7d**). Notice that while the bright yellow color-coding of the isopleth distributions falls within the bounded territory color-coded as bright green in the choropleth map, the yellow distributions do not completely occupy this green territory but, rather, mark a subset of immunoreactive elements in this area, illustrating how the isopleth distributions provide finer spatial resolution of the distributions at the σ value we assigned (see *Section 2.5.2*). Having understood these types of data visualization, we next turn to how readers can utilize the Adobe® Illustrator® (<u>Ai</u>) vector graphics platform to access and use our database files. Users can also visualize *Chemopleth 1.0* files in the freely downloadable software, Inkscape.

### 3.4 : Navigating Chemopleth 1.0 files using the Adobe® Illustrator® (Ai) graphical user interface

To accurately use the master files of the database, it is important to keep the dimensions of the original <u>Ai</u> files downloaded from the *Brain Maps 4.0* (BM4.0) publication (Swanson, 2018). There are eight atlas levels (numbered 23–30) from BM4.0 covering the anterior and tuberal *hypothalamus (Kuhlenbeck, 1927)* (HY), and portions of the *zona incerta (>1840)* (ZI), of an adult male rat. An example of a database file opened in <u>Ai</u> is provided in **Figure 8**, where BM4.0 level 23 was opened (using Adobe® Illustrator® 2020). Note that the layout might look slightly different depending on what workspace option you choose. Here we have the workspace set in “Essentials Classic mode”; however, any workspace will have the same tools, just with a different layout. To change your workspace, go to Window > Workspace > [Essentials Classic]. In **Figure 8**, three panels are highlighted in yellow, within which key control are outlined in cyan:

**Figure 8.**
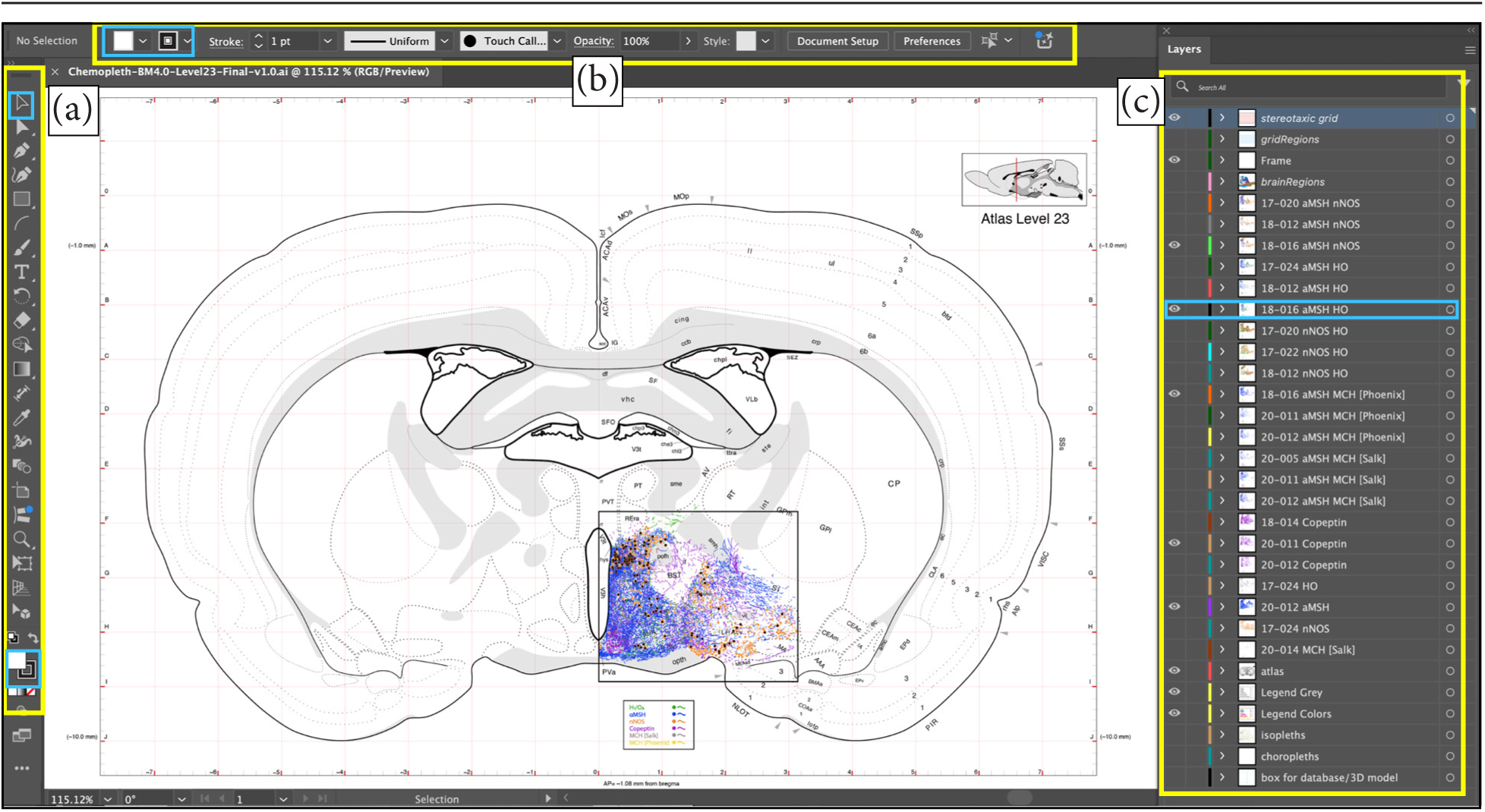
An example of an atlas level with mapped data, which together are part of the *Chemopleth 1.0* database, opened within the Adobe® Illustrator® (<u>Ai</u>) workspace. The artboard displays a base map of a transverse section of the rat brain from the open-source *Brain Maps 4.0* (BM4.0) atlas (Swanson, 2018). In this example, BM4.0 atlas level 23 is shown, with the contents of the database shown in the boxed outline on the base map (*black rectangle*), which consist of patterns of immunoreactivities drawn from photomicrographs of several experimental subjects directly onto the base map (in different colors for each neurochemical label). (**a**). *Tools panel* (left side, vertical) for <u>Ai</u> outlined in *yellow*, with the *Direct Selection Tool* outlined in *cyan* toward the top of the panel and the *Fill and Stroke Tool* outlined in *cyan* toward the bottom of the panel. (**b**). *Control panel* (top, horizontal) for <u>Ai</u> outlined in *yellow* with the *Stroke Tool* outlined in *cyan*. (**c**). *Layers Panel* (right side, vertical) for <u>Ai</u> outlined in *yellow*, with data for one of the specific experimental subjects shown in the map outlined in *cyan*. See the text for an explanation of these panels.

Explanation of select layers:

*stereotaxic grid*, stereotaxic grid lines;

*gridRegions*, regions covered in this study by grid lines;

*Frame*, square surrounding the area of the *hypothalamus (Kuhlenbeck, 1927)*;

*brainRegions*, enclosed brain regions enabling the tabulation of cell counts and fiber area estimates by regions.

*atlas*, base map from BM4.0; *isopleths*, trends by average counts; *choropleths*, average counts by region;

*Box for database/3D model*, frame embedded in the database for future registration of additional data in 2-D or 3-D.

<u>Abbreviations</u>: aMSH, alpha-melanocyte-stimulating hormone; nNOS, neuronal nitric oxide synthase; HO, hypocretin 1 / orexin A; MCH, melanin-concentrating hormone. *Chemopleth 1.0* data files consist of eight atlas levels, with each level saved as a separate SVG file that can be opened in Adobe Illustrator or Inkscape. Please refer to the text for details.

*Tools panel* **(Fig. 8a)**: contains all the tools to use <u>Ai</u>. By hovering over the icon tools, you can see the name of the tool and a keystroke or keystroke sequence in parentheses which is the shortcut for the tool. Two tools are highlighted in cyan: (1) the *Direct Selection Tool* (at the top), the icon is a black arrow. This tool will help you select your object on the artboard/canvas. (2) the *Fill and Stroke Tool* (at the bottom) represents the fill color (top black open square) and stroke color (background white square) of any selected text or object.

*Control panel* **(Fig. 8b**): according to the tool you are using from the *Tool panel*, this panel will display additional specific tools for objects or text you have selected. In this example, the *Stroke Tool* is outlined in cyan, which lets you choose the size of your stroke (outer thickness).

*Layers panel* **(Fig. 8c)**: This is where all the objects in your artboard are listed. Each row is a data layer (transparent digital overlay), and users can toggle on or off the visibility of the data for that layer on the base map by toggling on the *eye icon* to the left of each layer or by leaving it toggled off, respectively. The *lock icon* is to lock the layer, preventing any moving or deleting of data on that data layer.

### 3.5 : Detailed contents of the database

*Chemopleth 1.0* data files (Navarro et al., 2025; available at: https://doi.org/10.5281/ <u>zenodo.15788189</u>) can be viewed more closely for their layer organization via the *Layers panel* in the <u>Ai</u> or the *Layers and Objects* panel in Inkscape. There are at least three experimental subjects per pair of immunoreactive antigen, and each has sublayers with their corresponding antigen, with additional nested sublayers for appositions, cell bodies, and fibers **(Fig. 9)**. Layers contain artwork (i.e., the data) and are arranged in a stack. As shown in **Figure 10**, content on <u>Ai</u> layers located at the top of the *Layers panel* (e.g., “Frame”, “brainRegions”) appear *on top of* content on layers that are lower in the *Layers panel* (e.g., “atlas”, “isopleths”, “grid”). Isopleths show spatially averaged density maps of the distributions of immunoreactive elements, showing regions of consensus (and discordant) distributions across multiple subjects independent of cytoarchitectural boundaries marked on the underlying base map. In contrast, choropleth maps provide distribution gradients normalized and bounded by the cytoarchitectural boundaries of the underlying base map. See Peru (2020) and Navarro (2020) for details concerning the distributions of these immunoreactive biomarkers and their biological significance.

**Figure 9.**
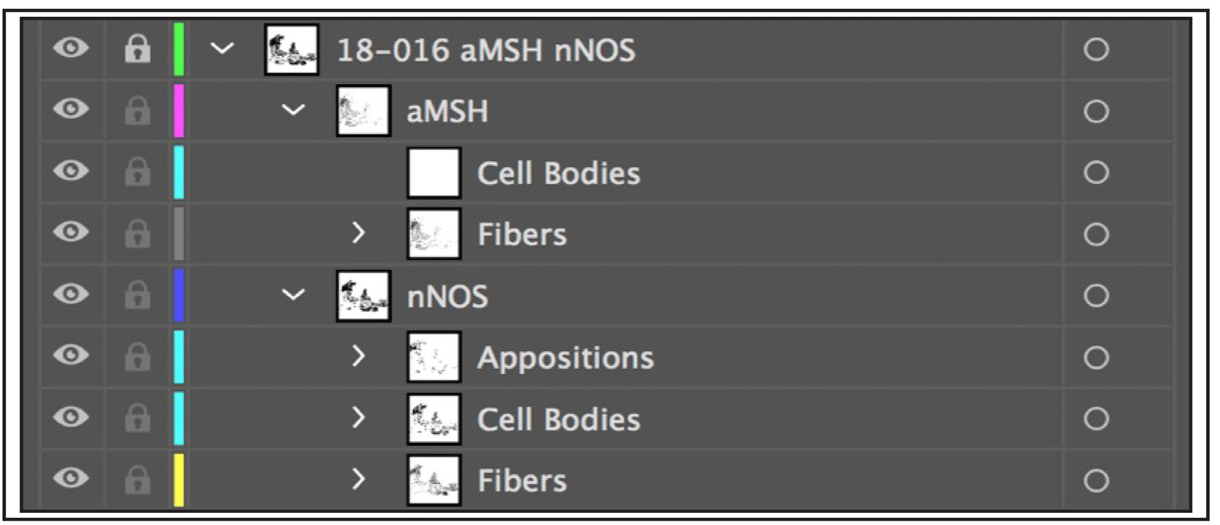
An enlarged view of the *Layers panel* in the <u>Ai</u> workspace, listing the data layers for subject #18-106 within the *Chemopleth 1.0* spatial database. The label for the main layer (*top row*) shows that the data are for dual immunoreactivity labels for the neuropeptides alpha-MSH (aMSH) and nNOS. Clicking on the *arrowhead* (>) to the left of the main layer label opens sublayers for each biomarker, under which additional nested sublayers appear if the *arrowheads* (>) to the left of each sublayer label are toggled in turn: those for cell bodies (perikarya) and neurites (axonal fibers and/or dendrites), and in some cases, appositions. See text for details.

**Figure 10.**
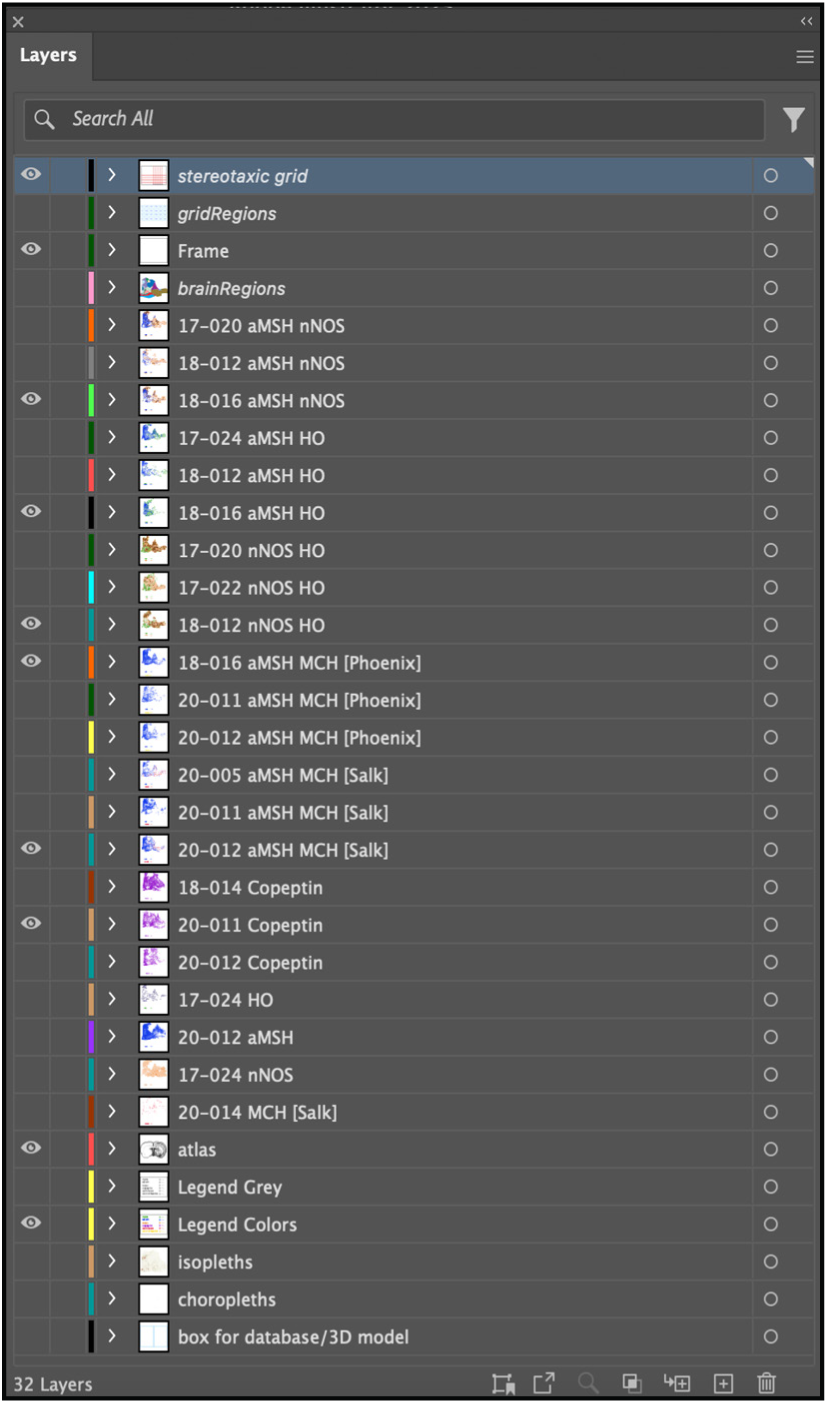
Detail of the *Layers panel* in <u>Ai</u> workspace, showing various layers and their arrangement within the *Choropleth 1.0* database.

Figure 11 provides an illustrative example of how the data overlay in each layer of the *Layers panel* in the <u>Ai</u> workspace can be toggled to visible sequentially to assemble a composite map of multiple antigens simultaneously for any given atlas level (this can likewise be achieved using Inkscape). In the example shown, level 26 of the BM4.0 atlas serves as the base map (Fig. 11a), upon which nNOS (Fig. 11b), H_1_/O_A_ (Fig. 11c), αMSH (Fig. 11d), MCH (Fig. 11e), and copeptin (Fig. 11f) immunoreactive signals mapped to this space as separate overlays are revealed, stepwise. The final composite map displaying all five antigens assembled co-spatially is shown in Fig. 11f.

**Figure 11.**
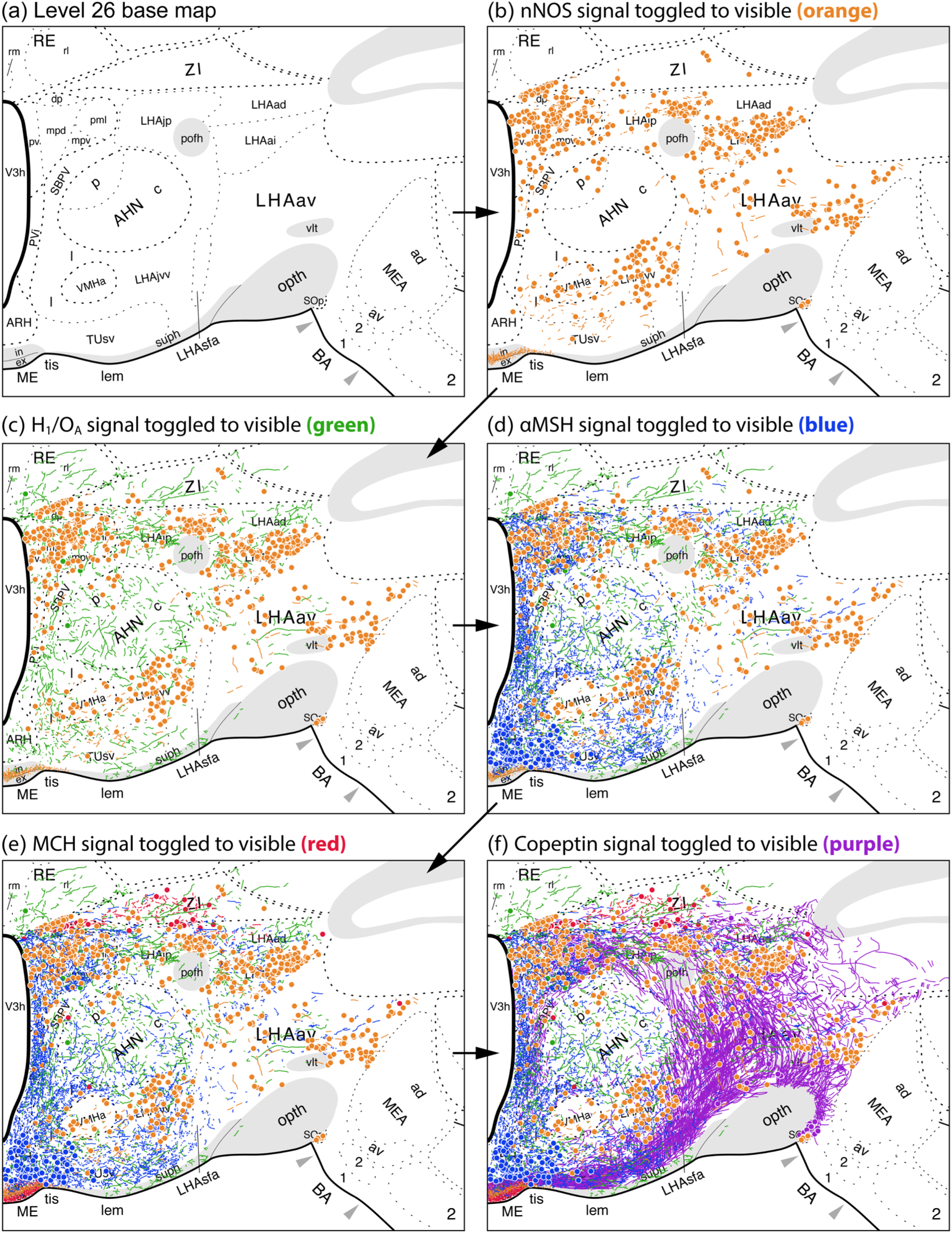
An example of successive reveals of different immunoreactive patterns for the five neuronal antigens mapped in the *Chemopleth 1.0* database. **(a).** A cropped portion of the Level 26 base map from *Brain Maps 4.0* used in the database. Toggling on the visibility for the nNOS immunoreactive signal displays the perikarya (*circle glyphs*) and small neurite extensions (*tiny strokes or lines*) of the mapped elements for this antigen (**b,** *in orange*). Additional toggling of the visibility of additional layers reveals, in succession superimposed signal for hypocretin 1 / orexin A (**c,** *in green*), alpha-MSH (**d,** *in blue*), melanin concentrating hormone (**e,** *in red*) and copeptin (**f,** *in purple*). See the list of abbreviations for an explanation of the atlas region symbols.

#### 3.5.1 : Single-subject maps of immunoreactive perikarya and neurites

Fig. 12 provides a summary of how *Chemopleth 1.0* contains a base map for each BM4.0 atlas level (see **1** in Fig. 12), upon which single-subject maps were produced for each immunoreactive system we labeled, using copeptin-immunoreactive neurites as the example (Fig. 12, **2**) (in this instance, only the neurites immunoreactive for copeptin are made visible by toggling off the visibility for the layer with copeptin-immunoreactive perikarya).

**Figure 12.**
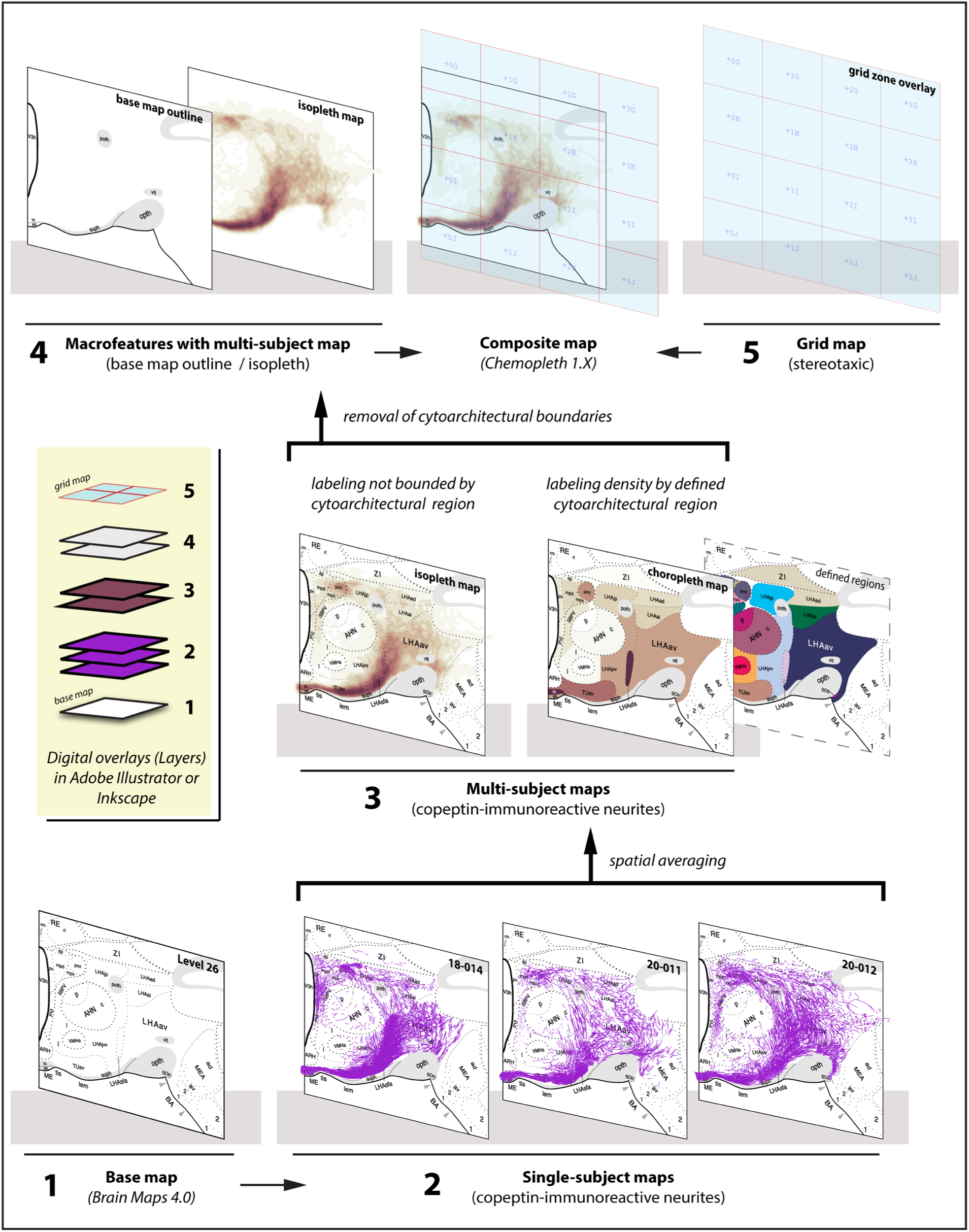
Illustration of the relationship among the base map, single-subject maps, and multi-subject maps of the *Chemopleth 1.0* database. The base map (**1**) from the *Brain Maps 4.0* atlas forms the base layer in the Layers stack shown (*yellow outlined stack, middle left*). Stacked as transparent overlays on this base map are single-subject maps (**2**) and multi-subject maps (**3**). Macrofeatures of the data (**4**) along with a stereotaxic grid map (**5**) form the final composite map of the database. See text for details.

#### 3.5.2 : Multi-subject isopleth and choropleth maps

Fig. 12 also shows two types of multi-subject map, which were produced by exporting the single-subject maps (as described in the Fig. 1 workflow) and computing across them to produce averaged “heat maps”. The color-coded density scorings of these maps were binned by cytoarchitectonic region (to produce a choropleth map) or sorted into a distribution that was independent of the cytoarchitectonic boundaries of the base map (thereby producing an isopleth map) (Fig. 12**, 3**).

#### 3.5.3 : Interaction maps

For certain experimental subjects (see Figure 9) we also performed a spatial analysis of the locations for putative synaptic appositions between two neurochemical systems. For these cases, an additional map was placed in a Layer called “Appositions”.

#### 3.5.4 : Grid-based coordinate system

##### 3.5.4.1 : Definition of the coordinate system

This coordinate system is displayed as an 8.0 × 12.0 mm grid overlay (layer) within each <u>Ai</u>-formatted electronic BM4.0 atlas level template used in this study, with the origin located on the ventromedial corner of the coronal plane hemi-section **(Figure 13a)**. The *x*-axis of this grid (which marks the mediolateral (ML) stereotaxic coordinates of each atlas map) is 8.0 mm in length, and the *y*-axis of the grid (which marks the dorsoventral (DV) stereotaxic coordinates of the map) is 12.0 mm in length. Note, however, that the maximal dimensions vary depending on the position of the brain section within each specific atlas level (i.e., some levels will have tissue expanses spanning fewer lengths).

**Figure 13.**
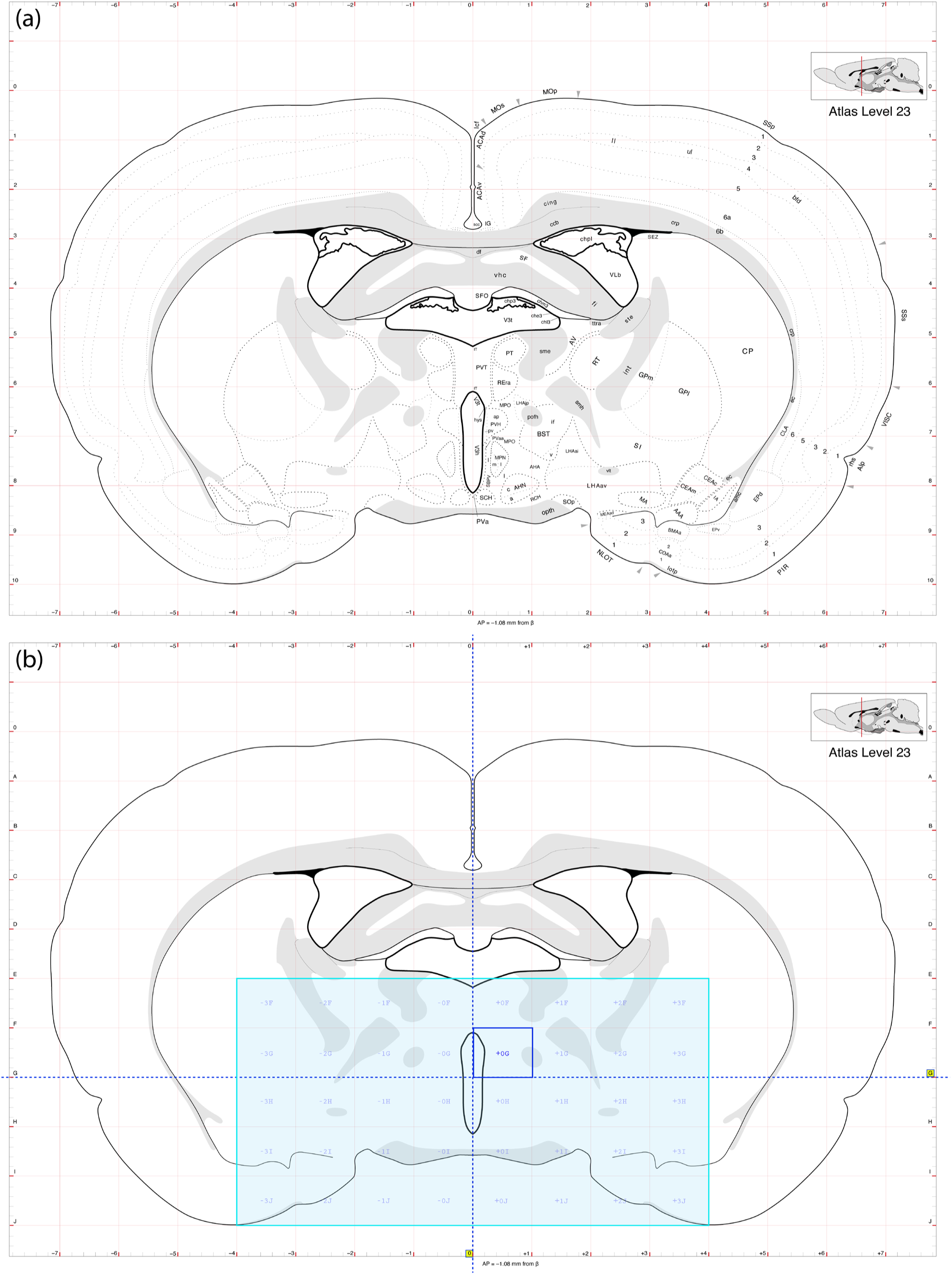
(**a**). Feature-based map with stereotaxic coordinates. Detail of the *Chemopleth 1.0* data layers showing the BM4.0 base map for atlas level 23 and feature-based boundaries (such as parceled brain regions). Also shown is the digital overlay for the sterotaxic grid included in the original BM4.0 atlas template (*red-lined grid*). (**b**). Grid-based map with stereotaxic coordinates. Detail of *Chemopleth 1.0* data layers showing the basic outline of the BM4.0 base map for atlas level 23 along with white matter tracts. A new grid region system (*light blue tiles and labels*) is shown over the base map, dividing portions of it according to a new naming system for each grid region, based on sterotaxic grid (*red-lined grid*) position of the tile (see text for details).

We define the lines parallel to the *y*-axis (sagittal lines, or *sagittals*) by the numerals 0–7, with 0 marking 0.0 mm ML (the ML coordinate at the midline origin), 1 marking 1.0 mm ML, and so on, until the final numeral 7, which marks the penultimate sagittal at 7.0 mm ML. Sagittals located on the right-hand hemisphere of the map are denoted by a plus sign (+) before their respective numeral, and those on the left-hand hemisphere are denoted by a minus sign (–). For example, +5 marks the sagittal located +5.0 mm along the ML stereotaxic axis in the right-hand hemisphere map of a given BM4.0 atlas map. In similar fashion, we assigned alphabetical designations (A–L) for the lines parallel to the *x*-axis (horizontal lines, or *horizontals*), which correspond to numerical values of the dorsoventral coordinate (–1.0 to –12.0 mm). Note that these numbers grow more negative in value from dorsal to ventral along the map. Thus, J denotes the position of the horizontal that is 10.0 mm ventral (i.e., –10.0 mm DV) to the dorsal edge of the map (which is located at position 0.0).

We now describe how we use the sagittals and horizontals to create our grid annotations, much as “eastings” and “northings” of the Universal Transverse Mercator (UTM) grid system are used to annotate grids on geographic maps (Langley, 1998; also see chapter 4 of Headquarters, Department of the Army, 2005). First, the sagittal and horizontal designations can be used to identify specific *grid regions* or *tiles* within the map, using the sagittal as the first position listed for any coordinate pair, followed by the horizontal (thus, *x,y* = *sagittal*, *horizontal = ML, DV*). The notation for *grid regions* for our database is based on the 1.0 × 1.0 mm squares created by the stereotaxic lines and thus follows the format: ± (numeric signifier (0.00 – 8.00 mm) of the ML (sagittal) coordinate; alphabetical signifier (A–L) of the DV (horizontal) coordinate), with + marking coordinates on the right-side of the atlas map and – marking coordinates for the left side of the atlas map.

Second, we adopt the convention to name each tile based on the <u>ventromedial</u> corner intersection of each sagittal and horizontal. For example, tile +8E denotes the tile that has its *lower left corner* marking the intersection of sagittal +8 with horizontal E. Note that ventromedial positioning means that the left side of the atlas map would have the corresponding tile (–8E), with the *lower right corner* marking the intersection of sagittal –8 with horizontal E, since the lower right corner would be more medial than the lower left corner **(Figure 13b)**. We opted to designate the ventromedial corner rather than an absolute direction (e.g., “lower left” corner), since selection of the medial corner for corresponding locations on both hemispheres ensures that each grid region in one hemisphere has the identical but opposite-sign grid region name in the contralateral hemisphere. Thus –4G and +4G would mark grid regions in the left and right hemispheres for bilaterally corresponding territories of brain tissue space. These grid regions have been labeled in the grid digital overlay with a duplexed (uniwidth) TrueType typeface (Courier) at 35% opacity to allow for visualization of the elements in layers underneath, and so that alignment and spacing of the labels is uniform (Lupton, 2024, pp 52–53) even when the typeface is adjusted for preference by users of the database to be of a different weight (i.e., boldface). Finally, note that all levels of the BM4.0 atlas follow the plane of section of Paxinos & Watson (2014) in terms of coordinates being calibrated with respect to a “flat skull” position, so users of these coordinates should keep that in mind when targeting their regions of interest.

##### 3.5.4.2 : Populating stereotaxic space: Creation of the ‘FMRS’ annotation system

Using this refined grid-based stereotaxic coordinate system, we tabulated our counts for immunoreactive perikarya and/or neurites within this grid using a new annotation system we call the *FMRS annotation system* (FMRS = “<u>F</u>requencies <u>M</u>apped with <u>R</u>eference to <u>S</u>tereotaxy), which essentially are counts of these immunoreactive elements *binned by grid region* as opposed to brain regions (Figure 14), thereby generating a *grid-region-based choropleth map*, which we call an *FMRS*. An *FMRS*, an acronym we use as a noun and pronounce like the word *ephemeris* (see *Section 4.2.2.2* for rationale), is a table of positions of spatial data points (in this case, profile counts) in stereotaxic coordinate space obtained from a single observation or a few observations (i.e., a single subject or a few subjects), with each entry in the *FMRS* consisting of the following values: anteroposterior (bregma) coordinate (in mm), grid region or tile defined by a specific sagittal and horizontal intersection, and profile count (frequency of perikaryon profiles or a pixel-density score for neurites) in that grid region.

**Figure 14.**
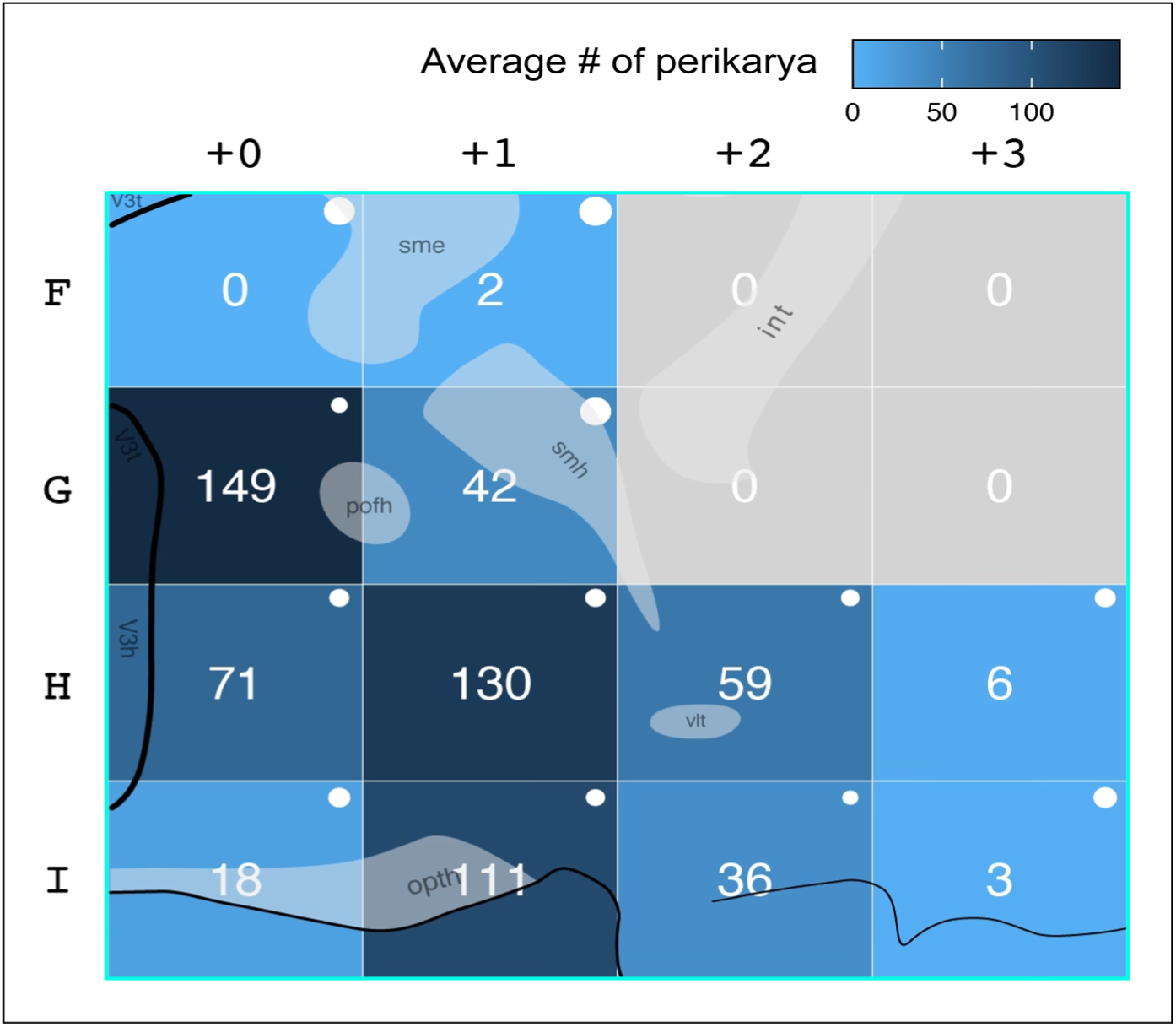
An FMRS for nNOS-expressing perikarya counted by grid region. Note that each region is color-coded according to the density gradient shown (*top right*), and that the macrofeatures of the brain are visible under the grid regions as a reference to the underlying features from the Atlas Level 23 BM4.0 base map (*see text for details*). The sizes of white dots mark relative values for the standard error of the mean (SEM).

We generated several FMRS, each keyed to a single bregma coordinate. To construct each FMRS from any data-populated BM4.0 atlas level within <u>Ai</u>, the visibility of the data layer featuring the vector-formatted cytoarchitectonic boundaries was toggled off, and the visibilities of the data layers with glyphs for immunoreactive perikarya or neurite profiles and the Paxinos & Watson-derived stereotaxic grid were toggled on, allowing the profile counts to be parsed by grid region within Swanson (2018) reference space.

#### 3.5.5 : Legacy datasets in register with new data

Figures 15–17 show the results of placing legacy data in register with our current maps in *Chemopleth 1.0*. Specifically, we migrated the data from Kerman et al., (2007), which was mapped in BM3.0 (Swanson, 2004) reference space, into BM4.0 reference space and placed their data in register with our datasets for MCH- and H_1_/O_A_-immunoreactive perikarya distributions (Fig. 15). We also migrated spatial data for nNOS- and H_1_/O_A_-immunoreactive perikarya distributions reported by Yao et al., (2005) from BM2.0 space (Swanson, 1998) to BM4.0 space (Swanson, 2018) **(Figs. 16, 17)**. In all instances, there were striking concordances of spatial overlap between legacy datasets and our own mapped datasets, providing a powerful convergence of evidence supporting the accuracy of the mapping.

**Figure 15.**
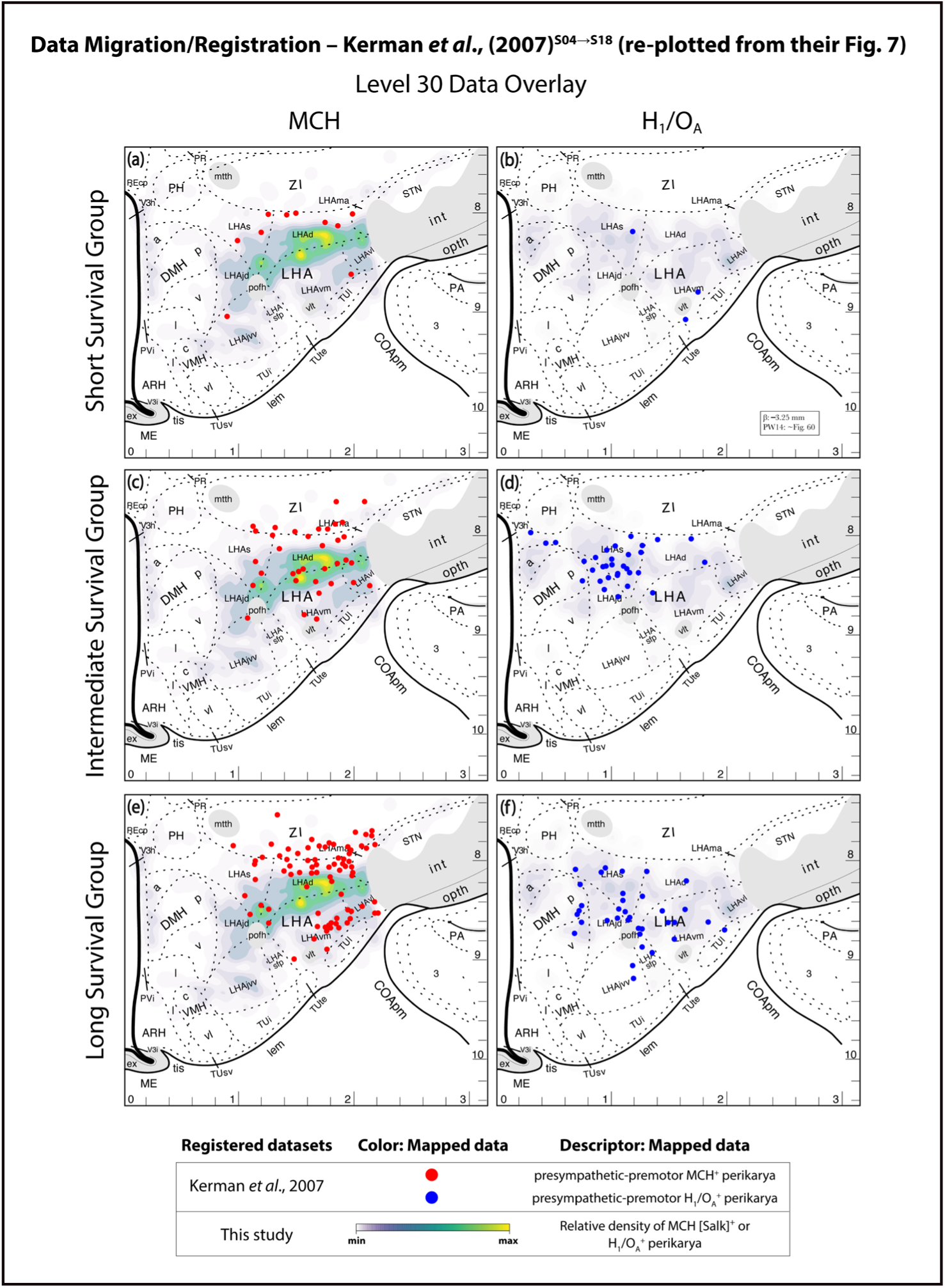
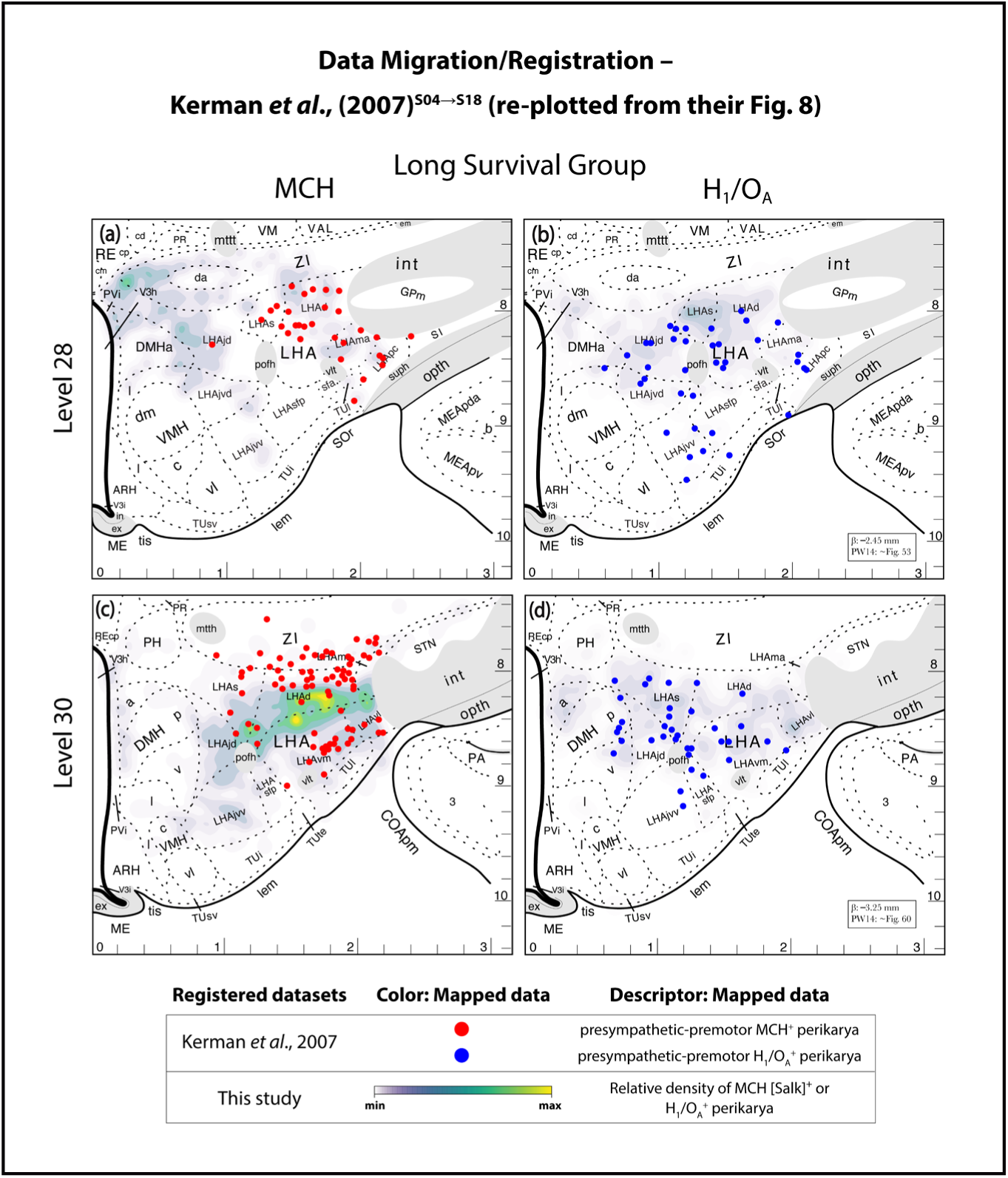
Example of data migrated and redrawn from Kerman et al., (2007) into *Chemopleth 1.0* and placed in registration with the database’s relevant datasets.

**Figure 16.**
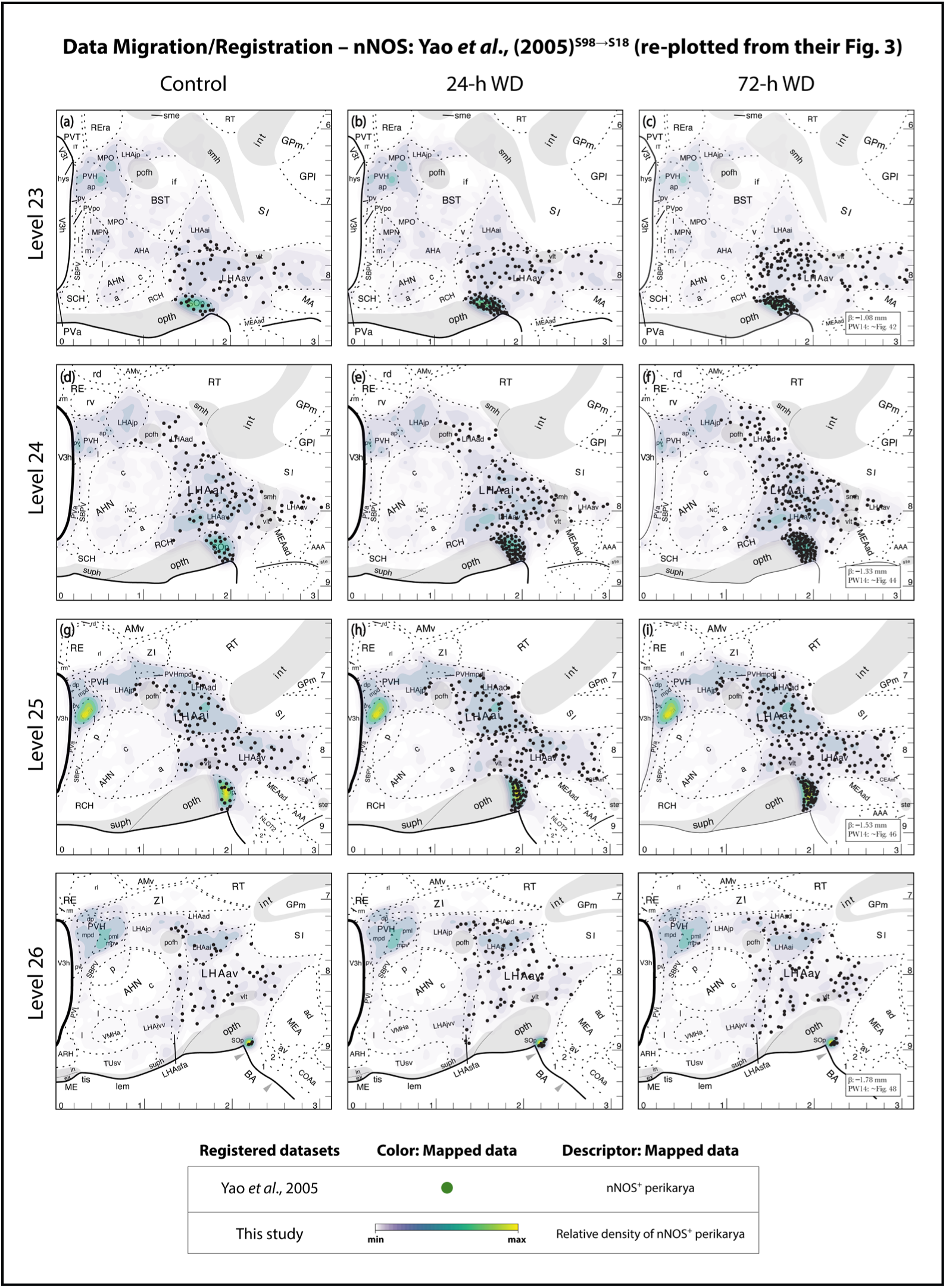
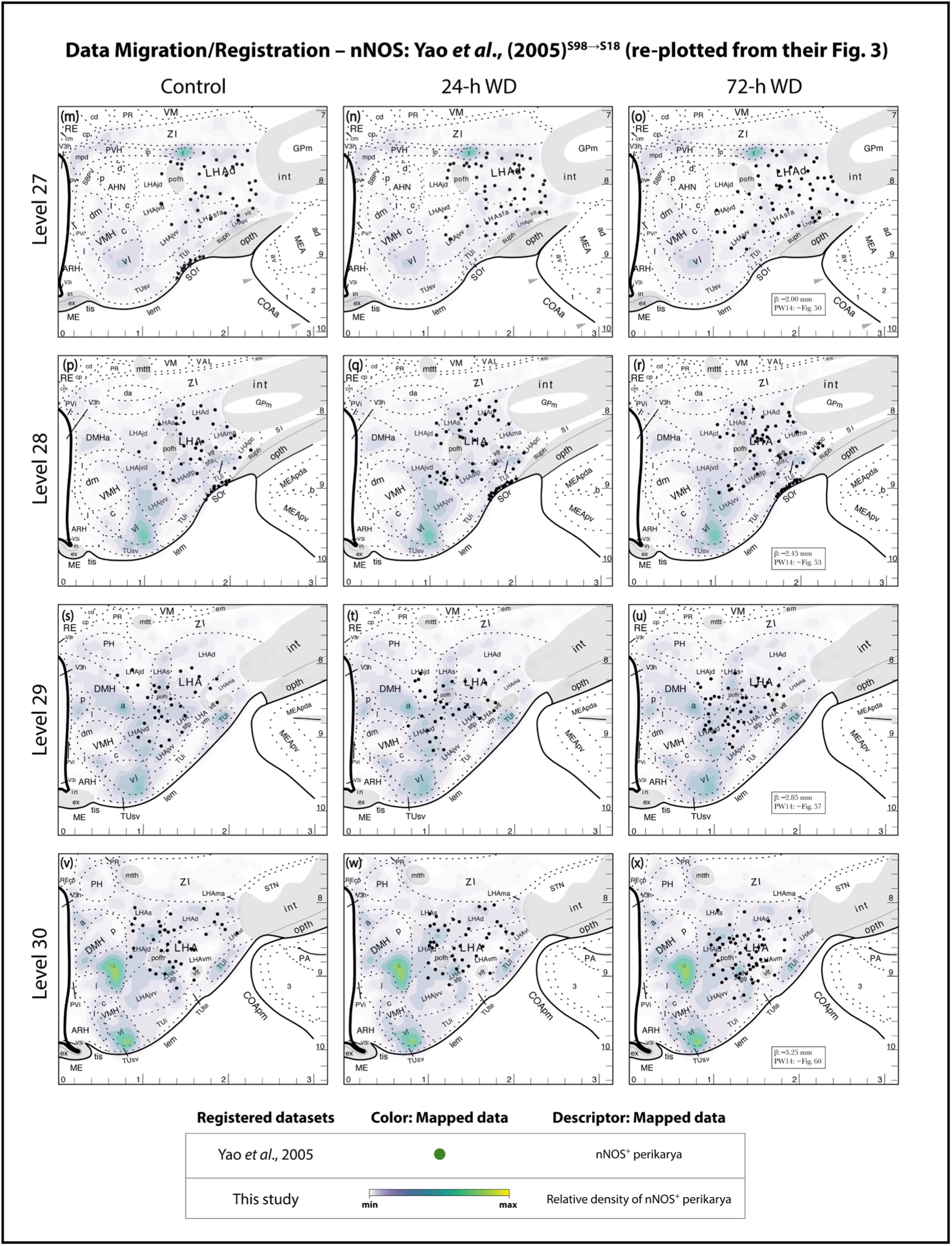
Example of data migrated and redrawn from Yao et al., (2005) into *Chemopleth 1.0* and placed in registration with the database’s relevant datasets.

**Figure 17.**
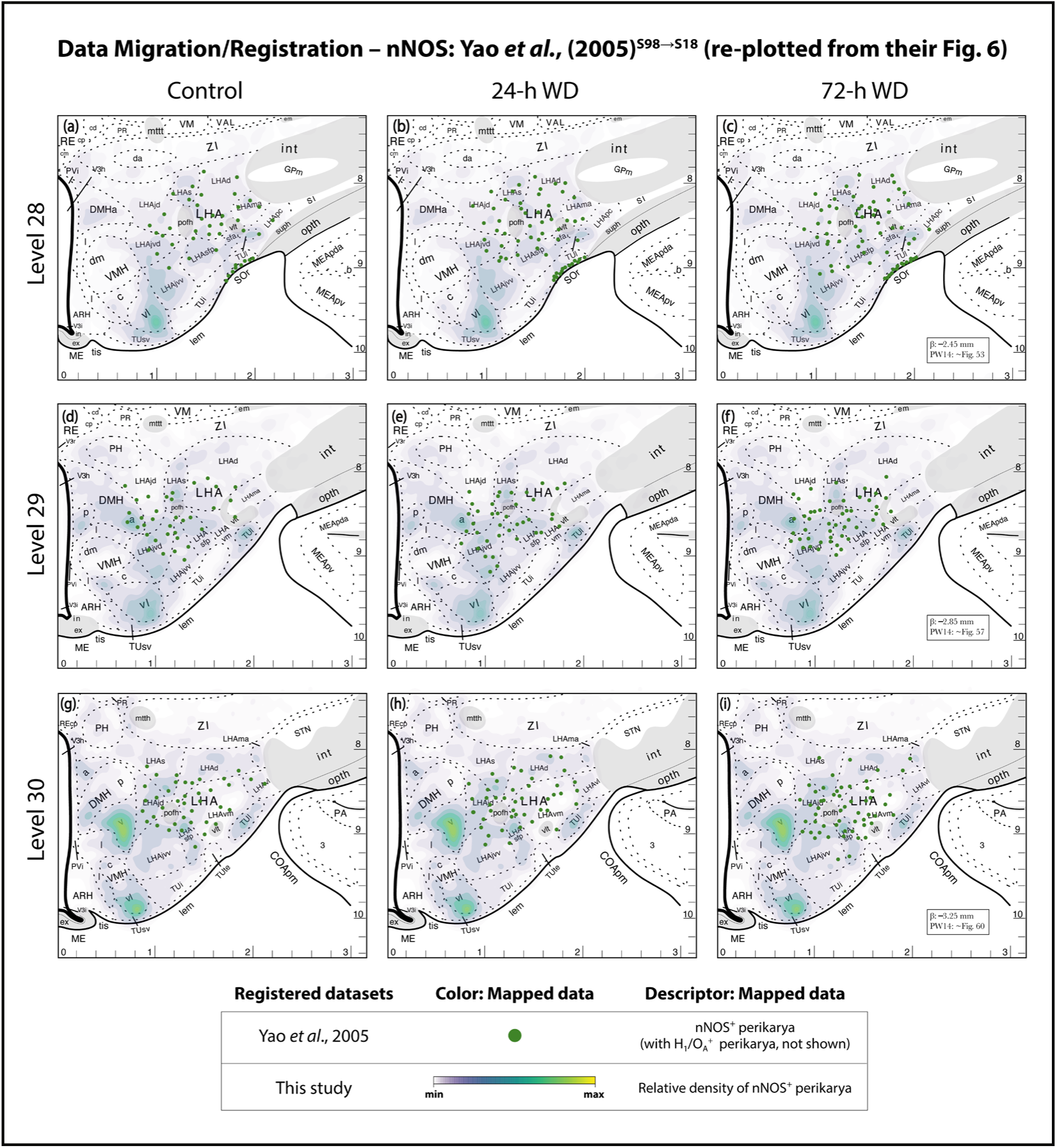
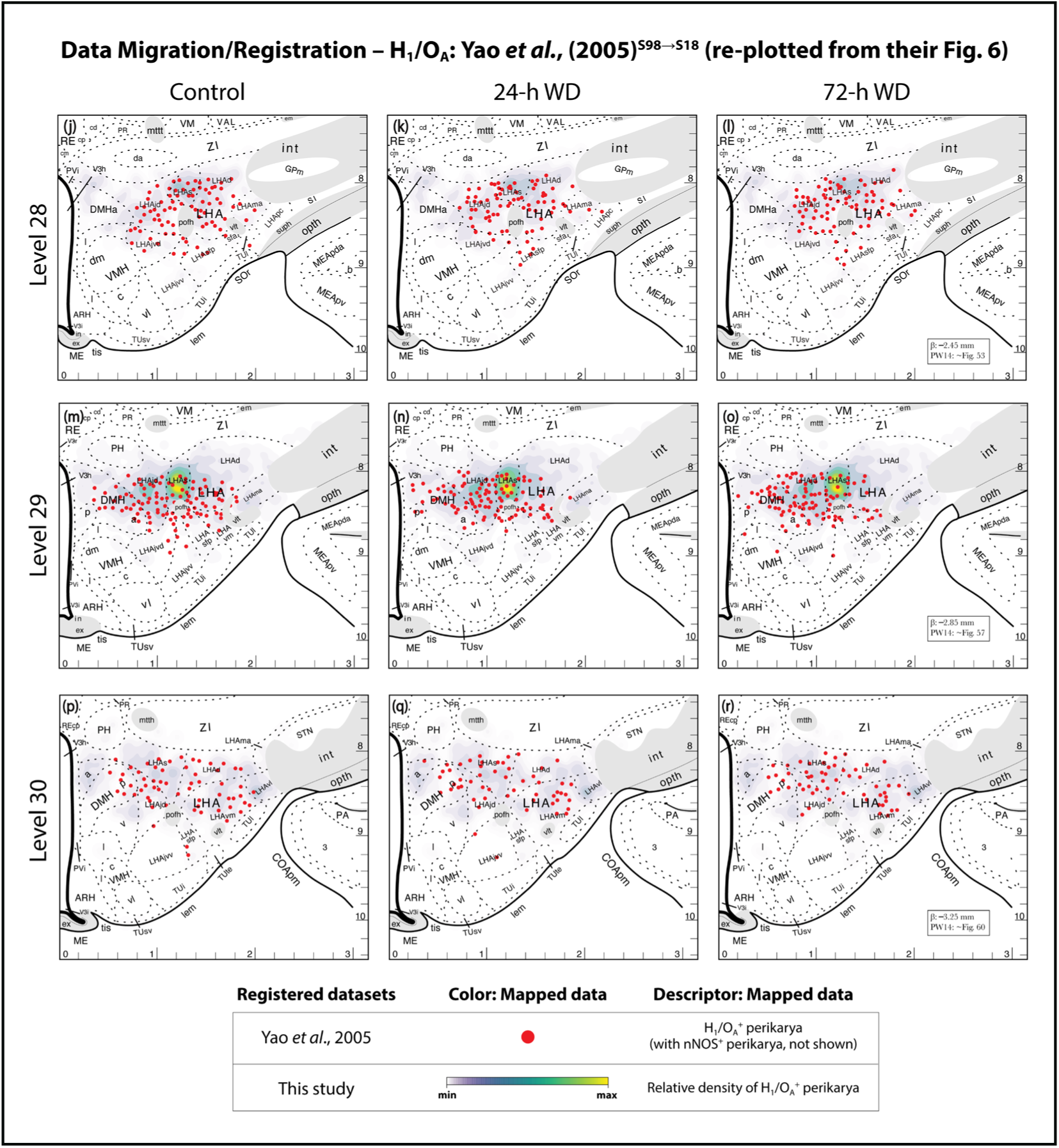
Example of data from Yao et al., (2005), redrawn onto BM4.0 templates to migrate them into *Chemopleth 1.0* and place them in registration with relevant datasets.

## 4 Discussion

In this study, we populate the base maps freely available from the *Brain* Maps 4*.0* rat brain atlas (BM4.0; Swanson, 2018) with new datasets generated in our laboratory to create *Chemopleth 1.0*, an interactive spatial database of chemoarchitecture for portions of the rat hypothalamus and zona incerta. *Chemopleth 1.0* data files (Navarro et al., 2025) are freely downloadable as SVG atlas files from the Zenodo data repository (https://doi.org/10.5281/zenodo.15788189) and can be opened using Adobe Illustrator (<u>Ai</u>) or the free software platform, Inkscape (www.inkscape.org). Each downloadable file provides a detailed view of the cell body and axonal fiber distributions of five neurochemical systems to aid in their targeted functional examination in stereotaxic space. This new resource allows the user to export cell and neurite data from the atlas in tabular and raster formats, compute on these data, and export data visualizations of spatially averaged “heat maps” derived from individual data maps from multiple subjects. The scripts for this workflow are available in <u>GitHub</u>. The file format of the maps produced from these scripts permit their re-importation into the native atlas environment. The database includes two types of these maps, one providing density estimates by cytoarchitectural region (choropleth maps) and the other providing density patterns that are computed across territory in a manner independent of the underlying cytoarchitectural boundaries (isopleth maps). Additionally, we introduce a new grid-based annotation system, based on the stereotaxic grid provided in the underlying atlas, to allow non-expert neuroscientists to map their own data within the atlas without needing training in detailed cytoarchitectural mapping methods. This grid system is also available in each downloadable *Chemopleth 1.0* data file as a digital overlay (layer). Finally, we provide guidance in the Results section of this report on navigating the <u>Ai</u> workspace to access and make sense of the spatial data contained in the database.

Below, we discuss the need for this database and then focus on its contents and design features. Throughout, we reflect upon the database’s design, environment, and assumptions, with discussion that draws upon research in geology, geography and geographical information sciences, the humanities, and the visual, literary, and performative arts. The net is deliberately cast wide to emphasize the shared ‘experimental entanglements’ (Fitzgerald & Callard, 2015) of our collective efforts to model, visualize, and make sense of complex, spatiotemporal data and to also learn how such tasks are carried out by those outside our immediate discipline.

### 4.0 : Why do we need *Chemopleth 1.0*?

Ultimately, the visual record of the chemical transmitter systems here, charted as they are within standardized maps with universal coordinates to target these systems, should prove useful for scientists studying these systems in detail. For certain applications, maps at this resolution may not be required, à la Borges (1964). However, for those with more pressing needs to search for finer-scale targets, maps such as the ones furnished in this database are likely as critical – and as rare – for localizing mesoscale brain circuits as similar records are for other poorly explored frontiers, such as the deep seafloor (Bell et al., 2025). Importantly, they help to overcome several limitations of current approaches to chemoarchitectural study, as described next.

#### 4.0.1 : Limitations of photomicrography for chemoarchitecture

Traditionally, photomicrography has provided rich evidence for chemoarchitectural relationships in the rat hypothalamus and zona incerta. For example, apart from single-molecule-directed studies, too numerous to cite here, there have been detailed photographic treatments of immunoreactive nerve cell populations for multiple neurochemical systems in individual hypothalamic regions (e.g., van den Pol, 1982; van den Pol, & Tsujimoto, 1985; Simerly et al., 1986; Simerly & Swanson, 1987; Lantos et al., 1995; Broberger et al., 1998; Elias et al., 1998; Abrahamson & Moore, 2001; Yao et al., 2005; Kerman et al., 2007; Hahn, 2010), and in some cases, afferent systems (e.g., Abrahamson & Moore, 2001; Polston & Simerly, 2003; Füzesi et al., 2007; Johnson et al., 2018). However, these studies are often limited by the lack of precise coordinate data for stereotaxic targeting and usually show documentation of their labeling from only one or a few subjects, making it unclear whether the spatial organization of the labeling patterns is truly representative for the animal taxon being studied. Often, it is not immediately clear if the planes of section for labeled tissues between studies examining the same chemical system are comparable, since only a few of these studies have used standardized maps to create cytoarchitecturally delimited chartings of immunoreactive patterns registered to a common plane of section of a reference atlas. Moreover, those that carefully register their labeling patterns to approximate stereotaxic coordinates (e.g., Paxinos et al., 2022) have not included spatially averaged maps of these systems from multiple subjects. *Chemopleth 1.0* provides standardized maps, including maps of spatially averaged patterns, across multiple subjects for several neurochemical systems in stereotaxic coordinate space.

#### 4.0.2 : Limitations of extant maps of molecular information

The limits of photographic documentation of neurochemical systems extends to their molecular analysis as well. In the past three decades, the identification of molecular constituents of hypothalamic and zona incerta systems has outpaced their spatial mapping. This identification has involved transcriptomic, proteomic or peptidomic analyses of sampled tissues from these regions (reviewed in detail by Khan et al., 2018a). Most of the published molecular studies of this kind have either been performed in laboratory mice, were not intended to (and therefore do not) provide actual *physically mapped* spatial information but only dimensionality-reduction visualizations (e.g., tSNE or UMAP plots), and/or have sampled denser nuclei at the expense of more diffusely organized areas (reviewed in detail by Khan et al., 2018a). In contrast, older, more concerted mapping efforts have provided maps of rat brain chemical systems, but these have largely done so as part of larger whole-brain survey efforts (this is exemplified by the excellent *Handbook of Chemical Neuroanatomy* series, many volumes of which remain out of print and are not yet digitally available at the time of this writing). These studies included the hypothalamus and zona incerta, but they often lack the spatial resolution or coordinate space to facilitate precise targeting of the chemical systems with virally directed tools; such tools aid in targeting neurochemical systems in the rat, which has few transgenic lines (discussed in the next section). Finally, most published reports of chemoarchitecture for the rat hypothalamus and zona incerta have focused on mapping cell bodies; only a few have mapped axonal fibers or terminal fields, the locations for which are helpful for targeting retrogradely transportable vectors, or to help determine the neurochemistry of the underlying fiber systems being affected by deep brain stimulation strategies (Zhang et al., 2024).

#### 4.0.3 : Limitations of current transgenic rat lines

The laboratory rat has been a mainstay of neuroscientific research for over a century (e.g., King, 1910; Huber, 1915; see Oglivie (2007) for a nuanced historical contextualization), with a rich scientific literature concerning neural substrates and their causal links to behavior and physiology. While leveraging this information affords scientists with immense benefits when probing deeper into structure-function relationships for the nervous system, there historically have been limitations in the genetic tractability of this model organism. This, however, appears to be changing dramatically as new programmable nuclease tools gain currency.

Thus, at the time of this writing, a few transgenic rat lines exist to aid in labeling and identifying specific chemoarchitecture in the rat brain (Witten et al., 2011; Bäck et al., 2019; Zallar et al., 2019; Iwasaki et al., 2023), and more lines are being generated, especially using CRISPR/Cas9 gene editing methods (reviewed by Ma et al., 2017). In most cases, however, detailed comparisons of transgenic neuropeptide expression patterns with wild-type controls are still lacking, and no detailed standardized atlas-based mapping for transgenically expressed rat chemoarchitecture has yet been reported. In the more rigorously characterized lines, such as the TH-Cre rat, unexpected germline recombination with a parental sex bias has been reported (Liu et al., 2016). This is also a wider problem for numerous transgenic lines of mice and zebrafish used in neuroscientific research (Luo et al., 2020), reinforcing the calls for observing careful breeding and genotyping procedures when using these important and valuable model organisms (Song & Palmiter, 2018). Maps of wild-type rat brain chemoarchitecture, therefore, will be important for validating transgenic patterns of expression, even as the models undergo refinement and improvement and newly revised datasets come online. *Chemopleth 1.0* provides a useful starting point to facilitate such comparisons against five different neurochemical systems in wild-type rats.

#### 4.0.4 : Limitations of automated and semi-automated mapping methods

The need for maps of wild-type rat chemoarchitecture is also being felt in the fields of machine learning and computer vision, where efforts to automate or semi-automate the mapping of chemoarchitectural systems will require deep learning algorithms to improve their performance by computing on enough training data that can serve as ground truth. While a small amount of histological data can be bootstrapped and/or resampled to increase the amount of training data (e.g., Straathof et al., 2020; Arnal et al., 2021; Arnal, 2022; Arnal & Fuentes, 2022; Carey et al., 2023), and for certain deep learning approaches a small training set of macroscale and mesoscale histological data (MRI images; whole-slide images of histology) may suffice for effective neural network performance (Zoghby et al., 2024; Nan et al., 2025), it is also evident from other studies that a large amount of such data can be a better means to achieve accuracy in computer-vision-based pattern recognition methods (Sun et al., 2017; Singh & Singh, 2021). At the time of this writing, mesoscale histological datasets of immunohistochemically defined chemoarchitectural elements in the brain, such as those we are providing here, have not yet been evaluated to determine the minimal effective amount of training data needed for robust pattern recognition to facilitate their mapping to a standardized atlas with a high degree of precision and accuracy. Ground truth will also be required to validate emerging approaches that harness spatial transcriptomic information for use by artificial intelligence to compute region boundaries based on the spatial clustering of transcriptomic markers in brain tissue (e.g., see Lee et al., 2025).

#### 4.0.5 : Limitations of current 3-D visualizations

Finally, current methods to label and visualize rat chemoarchitecture in 3-D (for example, by light sheet microscopic imaging) arguably require spatial validation from 2-D tissue sets produced by separate methodology (e.g., Quintana et al., 2024), particularly with respect to cytoarchitectural patterns, where Nissl stains have long served as a gold standard and for which compatible methods in cleared whole tissue samples are still undergoing refinements. Such validation needs also extend to antibody staining, since it is often the case that antibodies rigorously vetted to produce 2-D immunostained patterns in tissues are not always optimal or workable in antibody-labeling methods for cleared, whole-brain samples (Richardson et al., 2021). Consequently, newer as-yet-unvetted antibody labels may be used instead, simply because of their compatibility with the cleared tissues, but the labeling patterns they produce may fail to pass muster when compared to the more rigorous 2-D labeling. As refinements in antibody stability and penetration in cleared tissue samples continue to be reported (e.g., Lai et al., 2022; Gao et al., 2024), an understanding of the disposition of such labels in tissues from antibody reactions in 2-D sections ought to keep apace to ensure rigor and reproducibility in reported labeling patterns from these newer efforts. Benchmarks are being established to support this approach (e.g., Yau et al., 2023), and our dataset in *Chemopleth 1.0* can be considered as a type of benchmark for this purpose.

### 4.1 : Feature- and grid-based maps

In *Chemopleth 1.0*, we provide single- and multi-subject chemoarchitectural data visualizations for five neurochemical systems across eight *Brain Maps* atlas levels through the anterior and tuberal *hypothalamus (Kuhlenbeck, 1927)* and parts of the *zona incerta (>1840)*. Each atlas level from *Brain* Maps 4*.0* (Swanson, 2018) is represented by a single base map of the hypothalamus/zona incerta that can be opened in Adobe® Illustrator® (<u>Ai</u>) or in Inkscape. The atlas file consists of a stack of data layers placed over the base map as digital overlays. The data layers are labeled according to experimental subject and the neurochemical immunoreactivity patterns being shown, with sublayers sorted according to the type of data being shown (immunoreactive patterns for cell bodies, neurites, and appositions). In addition to base maps and evidence-based cytoarchitecture, the database also includes the contents described in the Results and discussed in greater detail in the next subsections.

#### 4.1.1 : Feature-based maps

The first major class of map included in the database is called *feature-based* because this type of map not only documents the features of the immunoreactive patterns we are seeking to map, but also because these patterns are drawn with respect to brain regional boundary assignments that are first drawn as a *base map* in the atlas from the cytoarchitectonic features of Nissl-stained tissue series serving as the reference (Fig. 1: see ‘Reference Space’). Indeed, both the boundaries and the immunoreactive patterns are basic features of the landscape and are manually drawn by observation of photomicrographic data or, as in the case of other disciplines (e.g., Hill, 1991), drawings from direct lens-based observations. These feature-based maps are for single subjects (see next section) but can be transformed into multi-subject consensus maps through spatial averaging (see *Section 4.2.1.2*).

##### 4.1.1.1 : Single-subject maps of data elements (cell bodies, neurites, putative appositions)

Strictly speaking, the mapped elements mark the locations of *profiles* of cell parts: perikaryal profiles, axonal or dendritic profiles, putative apposition profiles (Coggeshall & Lekan, 1996). The locations of the perikaryal profiles drawn over the base map are represented in 1:1 registration with the location of the profile observed in photomicrographs of the native immunostained tissue. In the case of a putative apposition, the drawn profile marked the location that was in registration with the location observed on the actual tissue slide as it was visualized under a ×100 oil objective lens through the *z*-axis by a trained student (Navarro, 2020; Peru, 2020). Thus, one circle glyph marks the position of one profile for each perikaryon or putative apposition. To the extent possible, this approach was also followed for immunoreactive neurite profiles but was not always maintained for especially dense regions of neurites, where it was difficult to distinguish individual elements from each other. In those instances, the overall trends of density and directionality were drawn to represent the overall spatial organization of the data overlaid on the base map.

##### 4.1.1.2 : Multi-subject maps

One way in which neuroanatomical mapping between subjects can be registered is using deformation or warping algorithms that normalize a subject’s brain anatomy to fit a standardized template. Such efforts require validation by comparing “before” and “after” dispositions of known landmarks within the tissue, as has been performed for human MRI images (e.g., Grachev et al., 1999) and for mouse and rat brains (e.g., Gaser et al., 2012; Sergejeva et al., 2015; Puchades et al., 2019; Carey et al., 2023). In such instances, the landmarks selected are identified using an expert human actor trained to recognize such landmarks, providing an expert-guided validation (Carey et al., 2023). An alternative approach, where the comparison across subjects occurs without warping but through structural analysis, has also been conducted by various groups and formally described (Riviére et al., 2022). We opted for a hybrid approach for *Chemopleth 1.0*; in this case, each subject was *first* mapped by an expert to the BM4.0 standardized atlas and *then* the final map from each subject for a given atlas level was spatially averaged to produce the two classes of multi-subject maps described below. With this approach, the need to warp was obviated by the direct intervention of an expert mapper to transfer neuroanatomical data to atlas templates with the aid of Nissl-based parcellation and manual registration.

##### 4.1.1.3 : Choropleth and isopleth mapping

For the choropleth maps, a decision was made to produce these by first setting provisional boundaries for the open vector graphics representing the brain regions. At the time of this writing, most of the brain regions represented in BM4.0 vector graphics templates are not closed polygons. Therefore, as noted in *Section 2.5.2*, we first closed these visually apparent vector patterns for the eight levels we analyzed in this dataset. We note in that section how these were provisionally assigned and that future refinements of the *Brain Maps* reference space will no doubt change these boundaries as more data collection and deliberation takes place. The boundaries, therefore, constitute mapped design features that will undergo revision over time and the density scoring that we performed based on their areal measures is also, therefore, provisional, insofar as the permanency of the agreed-upon boundaries remains the convention.

A new tool introduced in this database is the isopleth map. Like choropleth maps, isopleth maps denote differences in areal density for the cell populations being mapped across multiple subjects, with the concentrations in brighter or more saturated shading denoting “core” population densities, surrounded by lighter shaded, lesser saturated territories representing sparser densities of the cells. However, unlike choropleth maps, isopleth maps display density distributions that are not confined to cytoarchitectonic boundaries of the base map.

The choropleth and isopleth color patterns we used are consistent with older observations, using monochrome LCD monitors, that darker regions on lighter backgrounds, with dark or saturated shades representing greater abundance, is generally most effective in conveying distribution differences (McGranaghan, 1996). Color management of data layers in mapping (Brewer, 2013) also informed our decision to use specific color patterns for our maps. Apart from color, we opted for cartographic generalization and, therefore, the sizes of the glyphs and strokes used to represent perikarya and neurites, respectively, are decidedly not to scale.

#### 4.1.2 : Grid-based maps

##### 4.1.2.1 : Stereotaxic grid

The original hard copy edition of *Brain Maps: Structure of the Rat Brain* (Swanson, 1992) included a transparent overlay that could be aligned to each base map, and which bore calibration marks for the stereotaxic coordinates for the Paxinos and Watson (1986) stereotaxic coordinate system. Their system is considered the gold standard coordinate system for targeting probes to brain structures mapped in their atlas or – in the case of *Brain Maps* – any reference atlas that is produced in register with the plane of the Paxinos and Watson atlas. Since that first edition, the electronic atlas map files of *Brain Maps* reference atlas have each included this stereotaxic coordinate system as a data layer. The *Chemopleth 1.0* database retains this data layer of inferred stereotaxic coordinates, and we introduce a new, named coordinate system for this coordinate framework that allows grid-based mapping of data for the first time. *Section 4.3.4* describes this design feature in greater detail.

##### 4.1.2.2 : The FMRS System

We selected the acronym, FMRS, because it phonetically resembles the word *ephemeris*, which is the term for a table of positions of celestial objects (e.g., planets) that has been used by many civilizations and cultures since antiquity (Gingerich, 1997; also see *zij*: Kennedy, 1956; King and Saliba, 1987; see Banneker (1793) for a notable example of a published ephemeris). We use “FMRS” in the same way as “ephemeris” is used in a sentence (i.e., as a noun and not an adjective; thus “an FMRS”, not “an FMRS table”). Also, we use “FMRS” as both a singular and plural noun: “one FMRS”, “many FMRS”. We opted to use FMRS rather than ephemeris for two reasons: (1) the acronym emphasizes its roots as a tabulation in stereotaxic space and not celestial coordinate space (i.e., <u>F</u>requencies <u>M</u>apped with <u>R</u>eference to <u>S</u>tereotaxy); and (2) it does not dilute, expand, or confuse the use of ephemeris, as used in astronomy, but still pays tribute to it as essentially the same concept: *an annotated tabulation of positional information for objects in a coordinate space*. Usually, an ephermeris has been defined as a table more for predicted rather than observed locations of objects (*Oxford English Dictionary*, 2025). In our case, the starting point is a set of observations in one brain or a set of brains, which could eventually form the basis of predicted spatial locations as more such data are gathered in a standardized manner and aggregated. We envision the FMRS to evolve over time from a descriptive to predictive tabulation when more observations are brought into register with the same grid regions of the atlas.

The finite samples populating our FMRS provide it a meaning that closely hews to a word related to ephemeris, namely, “ephemeral”, which denotes “short-lived” or “momentary”. For our purposes then, ephemeris (and by extension, FMRS), emphasizes that the counts or frequencies recorded in each grid region represent those of a *single experiment or observation* in one subject within a circumscribed set of space-time conditions, and that these frequencies are being used by a single laboratory to model (and, ultimately, given more time and more data, to predict) such distributions as they may appear across subjects. Therefore, they should be treated as a limited representation of more generalized states or phenomena. In other words, FMRS notations are of *circumscribed experimental observations*. Much as positional information tabulated in a community-based (and mature) ephemeris might, early on, need to be corrected to account for local conditions where the initial observations were recorded from, so, too, are the frequencies binned to stereotaxic grid regions dependent on the laboratory making the observations, the condition of the experimental subject they are studying, the conditions and parameters of their labeling experiments, etc. Over time, we anticipate that such tabulations will become more accurate as more observers bring their data to bear on the modeled dataset. In *Section 4.6.3*, we discuss the advantages that the use of this grid-based annotation system may afford investigators seeking a general summary of which brain areas to target experimentally and how such a system is a straightforward way to map distributions without needing training or expert knowledge in feature-based (i.e., cytoarchitectonic) mapping.

### 4.2 : Legacy datasets and their critical role in community atlas building

In this study, we re-plotted and brought into register with our own datasets the spatial positions of three neuronal populations reported in two studies: Yao et al., 2005 (hypocretin 1/orexin A, nitric oxide synthase) and Kerman et al., 2007 (hypocretin 1/orexin A, melanin-concentrating hormone). A few points are worth noting regarding this registration.

#### 4.2.1 : Aligning earlier and later work within a ‘spatial commons’ reveals concordance

First, the alignment of these data with our own isopleth maps of these populations reveals a strikingly concordant overlap across our collective studies. This alignment is striking because the methods used to map these populations were not shared between those laboratories and our own, nor were the methods used to generate the histological staining patterns identical. The registration underscores how standardized atlasing of data provides a means to readily explore similarities and differences among experimental results generated by investigators separated in time and space and who may use resources and methods that are similar, but not identical (this is illustrated for <u>photographic data</u> – standardized for the same region of night sky – for the Eagle Nebula (M16/ NGC 6611) from the Hubble Space Telescope in 1995 and the James Webb Space Telescope in 2022; see NASA, 2025). Similarly, neuroscientists who “share the same spaces” and are essentially “performing the same movements” (Farriss & Hutchence, 1990), can operate out of isolation and into a shared intellectual commons where they can help build, refine, and interrogate a shared spatial model of the brain (discussed in Khan, 2013; also see Leergaard & Bjaalie, 2022). Indeed, apart from the Swanson laboratory’s own use of the *Brain Maps* reference space, many other labs report mapped data for the hypothalamus or zona incerta plotted in this space, including mRNA and/or protein expression patterns (Broberger et al., 1998; Kelly & Watts, 1998; Khan et al., 2000; Yao et al., 2005; Hahn, 2010; Gu et al., 2013), chemical/electrode lesions or electrode placements (Gallistel et al., 1996; Flores et al., 1997; Simmons et al., 1998; Radley, et al., 2006; Choi et al., 2007; Martínez et al., 2023), chemical microinjection sites (España et al., 2001; Khan et al., 2007; 2018b), cellular activation patterns (Gu et al., 2013; Zséli et al., 2016), neuroanatomical tract-tracer injection sites and tracer-labeled cell bodies and fibers (Jansen et al., 1995; Abrahamson & Moore, 2001; Leak & Moore, 2001; Geerling et al., 2003; Uschakov et al., 2007; Kerman et al., 2007; Negishi et al., 2024; 2025; Lukinic et al., 2025), and optogenetic fiber placements (Wood et al., 2019). It is hoped that *Chemopleth 1.0* allows for such a model to grow and evolve for the rat hypothalamus and zona incerta by collective efforts of the scientific community. Ultimately, the use of inference engines and artificial intelligence across such layered datasets could provide explicit linkages among them and aid in hypothesis generation for future data-constrained experiments, as has been recently shown in archaeological research (Casini et al., 2023), and of greater relevance, in linking structural brain variations with down-scale genetics and larger-scale autism phenotypes (Kundu et al., 2024). Applying deep learning tools to the use of these maps to guide experimental and therapeutic targeting of this chemoarchitecture could literally and conceptually refine such linkages.

#### 4.2.2 : Digital overlays are a form of shorthand citation and preservation of evidentiary trails

Second, the use of artificial intelligence notwithstanding, the digital overlays afforded by the Adobe Illustrator (<u>Ai</u>) environment (*see Section 4.3.1*) already reveal explicit evidentiary and historical relationships among disparate sets of data. Figure 18 provides a simple example of how overlays can provide a shorthand form of citation that links a newly reported dataset (for example, the distributions of MCH-immunoreactive neurons; present study) with previous efforts “within” the same reference space (identification of premotor-parasympathetic MCH-immunoreactive neurons; Kerman et al., 2007). By migrating and registering these older data with our newer isopleth maps of these immunoreactive cells, it becomes clear that the majority of the cells mapped by Kerman et al., (2007) fall within the sparser contours of our isopleths of MCH-immunoreactive neurons, such that they appear to be situated around a hotspot of “core” cells. If this registration models an actual spatial difference between these populations (which it predicts as a testable hypothesis) this would suggest connectional differences between LHAd “core” cells and those in the dorsal margin of the LHAd or in regions adjacent to the LHAd, such as the ZI dorsally and the LHAvm/vl ventrally (Fig. 15e, Fig. 18).

**Figure 18.**
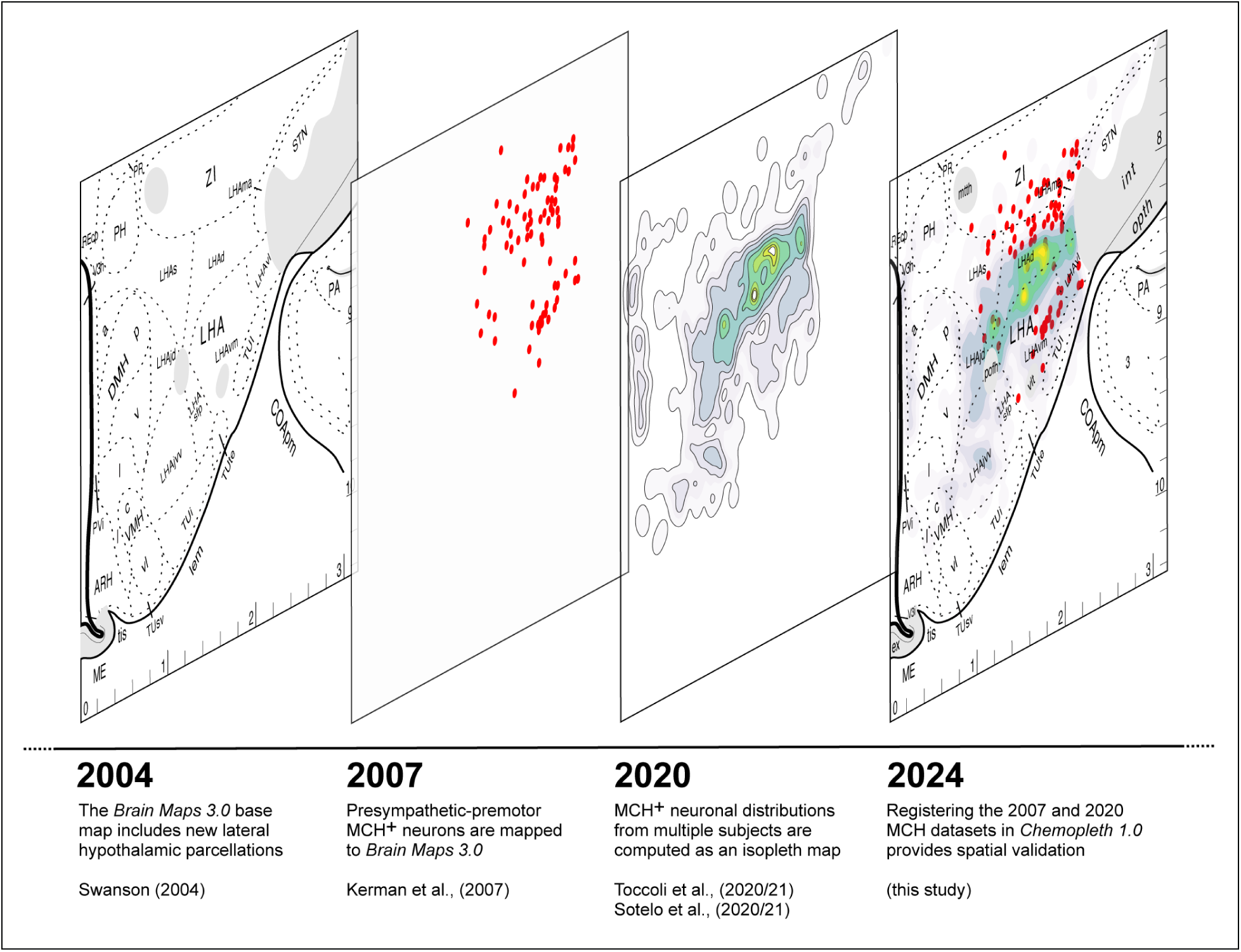
Community-based model building. An example of our data registration presented as a scientific chronology, illustrating a model-building process for the lateral hypothalamic area and zona incerta of the adult male rat in *Brain Maps 4.0* atlas reference space. In this example, the base map additions provided by Swanson (2004) allow finer-resolution mapping of traced MCH-immunoreactive populations in 2007 by Kerman and colleagues (presented as a separate data overlay in the database), which are supported by our successive isopleth analyses of multi-subject MCH-immunoreactivity patterns in 2020 (presented as a third overlay). The resulting model, shown at the far right, is, therefore, a community-built model of this chemoarchitecture. See text for details.

From the above example, it is clear that a prepared digital overlay of another laboratory’s data registered to a shared reference space is essentially a form of data curation from the literature that speeds up the process of understanding how such data relate to one’s own in that space. This approach of registering published datasets to these BM4.0 base maps is in keeping with the standard set by Swanson (2018) for the atlas itself, ever since its first edition (Swanson, 1992), where an appendix was included which relates most brain regions and areas to the published literature. The atlas is essentially a hypothesis (L. W. Swanson, personal communication to A. M. Khan), with each region’s boundaries undergoing refinements as more evidence furnished in the steadily accumulating scientific literature is brought to bear on that specific feature. This approach – that of linking structural features of the atlas maps to the published literature – also more formally underlies Swanson’s published Foundational Model of Connectivity (FMC), which states as one of its core principles that it is based on evidence not authority (Swanson & Bota, 2010).

In many ways, then, by linking our spatial datasets to the published evidence in the literature and citing those previous scientists who have furnished this evidence, we explicitly build upon those findings as we disseminate our own to the next group of readers who, in turn, may build upon what we have reported (see Burns et al., (2003) for an early model of this concept). This process is akin to the transfer of knowledge of spatial location in Polynesian cultures from one generation to the next in “a never-ending chain” (Foa’i & Miranda, 2016), only the difference is that their transfer is in oral, not written, form (Thompson, 2019). It also echoes the concept of ‘time-binding’ furnished by Korzybski (1950), as articulated by abstract artists such as Charles Biederman, where succeeding generations of human beings retain and build upon the knowledge and understanding from previous generations of how visualization of complex forms can be achieved (see p. 32–33 of Biederman, 1948; this concept is also implicit in Feynman’s “atomic hypothesis” discussion (Feynman et al., 1963/1989, page 1-2)). Complex and highly specific forms in art and music often find recurrences in later works, whether it be Greek key motifs in art, symphonic sounds in classical music (e.g., compare 1:29–1:50 of Handel, 1741/2009, with the flash of sound between 0:19–0:23 of Mendelssohn, 1825/2005), or guitar riffs in songs (e.g., compare 0:48 of Bowie, 1971, with 6:19 of Burton et al., 1986); the re-embedding of these motifs in later works simultaneously traces evidentiary trails of their origins while at the same time redefining them for new audiences. Similarly, the linkage in time and space of different mapped datasets explicitly points to recurring structure/function relationships in them; this includes re-contextualization of old datasets in relation to new ones, as we discuss next.

#### 4.2.3 : Older data fitted to newer models re-contextualizes them

As an example of how data can be updated to fit new models, we note that the data maps in both Yao et al., (2005) and Kerman et al., (2007) were plotted in those studies using older editions of the *Brain Maps* atlases. Thus, Yao and colleagues used the second edition of the atlas (*Brain* Maps 2), and Kerman and colleagues used the third edition of the atlas (*Brain* Maps 3). Since all extant editions of the Swanson atlas use the same experimental subject (i.e., the underlying tissue serving as the foundation of the maps is identical), the data in one atlas can be readily migrated to a later version, which is simply a refinement of boundaries and spatial relations on the same base territories in the same subject (i.e., it is not a different brain). Based on this rationale, in the present study, we migrated and re-plotted their published data to the latest edition of *Brain Maps* (*Brain* Maps 4*.0*; Swanson, 2018). The specific atlas levels Kerman and colleagues mapped their datasets to did not undergo any major change in *Brain* Maps 4*.0*, making their migration from the third to the fourth edition simply a carryover of the data into a similar space. In contrast, the migration of the Yao et al., (2005) dataset from the second to fourth editions *re-contextualizes* those data. This is a consequence of the evidence-based revisions made to the second edition’s subregional parcellations of the lateral hypothalamic zone. These revisions were a result of more detailed observations performed by Swanson of the original Nissl-stained tissue set on which the atlas is based (the editions are all based on the same brain: a single, 315-g male Sprague Dawley rat; also see Swanson et al., 2005). These considerations lead us to consider next the importance of data layers in a temporal context.

#### 4.2.4 : Data layers can constitute a chronological record

Much as data layers have been proposed for use by geoscientists to mark chronological events in geospatial datasets (Langran & Chrisman, 1988; pp 4, 9–11; also see Langran, 1993; Davoine et al., 2015; Saint-Marc et al., 2014; 2017), including real-world geospatial processes (‘world-time’; i.e., when the geological process took place) and their records/reported findings (‘database-time’; i.e., when the observation regarding the geological process was recorded; again, see Langran & Chrisman, 1988), data overlays can provide the same advantages for brain mapping. As stated by Langran and Chrisman (1993) in their treatment of time in geospatial datasets, we propose that a similar system should be adopted for registering legacy datasets to current reference spaces, since this form of data migration is important to track and memorialize, especially in terms of ‘database-time’, i.e., between sets of recorded observations in neuroscientific history. Model-building will require multiple snapshots in database-time as the spatial model of the brain is further refined. A standardized atlas, which importantly, also ought to undergo revisions over time, can serve as such a model, and spatial mapping such as that presented here can be databased over time and various experimental observations so recorded. Moreover, the older time-stamped datasets provide essential failsafe information should the later datasets or their aggregate analyses fail to “carry over” any essential features from the previous datasets; parameters that may be overlooked in the newer data may be present in the original, providing missing validation or critical context (as brilliantly illustrated in Rønning, 2025).

In the case of our simple exemplars of legacy database registration (Yao et al., 2005; Kerman et al., 2007), we propose that a standard notation be used for data migration, and that this notation be included as a tag accompanying a given dataset. Thus, Yao et al., 2005 (based on *Brain* Maps 2) migrated to the current *Chemopleth 1.0* database (based on BM4.0) could be written in notation form as the following dataset name: Yao et al., (2005)^S98^ ^→^ ^S18^, where S98 and S18 denote two sets of database-time; namely, data mapped to Swanson atlas editions from 1998 and 2018, respectively (as proposed by us earlier: Khan et al., 2018b) and the arrow denotes migration between the two atlas editions. We have used this notation in Figures 15–17.

### 4.3 : Digital Overlays

#### 4.3.1 : Background and history

In neuroscience, certain fundamental issues associated with the visualization of neural elements in space naturally arose from the desire to examine, in 3-D, information that was present in 2-D tissue sections under the microscope. This was difficult with the first microscopes, which were monocular lenses with limited depths of field (Ford, 1985), but these eventually gave way to binocular lenses and stereoscopic views where images arriving at both eyes were merged or superimposed to produce greater depths of field. Wheatstone’s stereoscope (1838) catalyzed the development of binocular lenses in microscopes, and Cajal’s manual on color photography and reports in his *Trabajos* describe his use of stereoscopy to visualize the nervous system (Cajal, 1912; 1918; also see Santarén, 2014).

Paralleling this need was the corresponding need to represent and visualize drawn 2-D representations of tissue elements viewed through their depth, with early attempts involving manual reconstructions via semi-transparent overlays of paper (e.g., Streeter, 1905). This need also grew out of limitations imposed by the finite cache of labels one could employ to visualize tissues under the microscope: “the gains in brain are mainly in the stain” (Bloom, 1975). Sequential images of different structural information could be viewed simultaneously using film technology (e.g., Levinthal & Ware, 1972), which allowed multiple photographs on film to be combined into a composite image using optical printing, as pioneered, for example, by Richard Edlund for *The Empire Strikes Back* (Lucas, 1980; discussed during 25:25–28:03 in Kasdan, 2022; also see Fordham, 2023). The history of the use of transparent overlays has proven to be elusive in its transition from analog to digital formats (e.g., see Cloud, 2005). However, as with the mainstream film industry in the 1980s (e.g., see Rose, 2022), computer graphics applications for neuroanatomy (“computer-assisted neuroanatomy”; see Woodward et al., 1985) largely supplanted film technology with digital data overlays, which were applied to neuroscientific data that same decade (e.g., Toga et al., 1984).

#### 4.3.2 : Chemopleth 1.0 digital overlays

The *Chemopleth 1.0* database is based on a digital overlay system that is historically embedded within the <u>Ai</u> environment from the time of its early version in 1993 (Adobe Illustrator, version 3.1). The use of this overlay tool within the first release of the graphics files of the *Brain Maps* atlas makes this rat brain atlas likely the first to utilize digital overlay technology and atlas maps formatted in vector graphics. For this atlas, instantiations of a graphic user interface with digital overlay tools were reported by Dashti et al., (2001) and Burns et al., (2006). *Chemopleth 1.0,* in many ways, can be considered as a complementary resource.

Overlays allow for both raster- and vector-format data to be superimposable in the same reference space. Historically, as with geomorphological mapping, usually the raster data are photographs (e.g., aerial photographs), which can be placed in register with points, lines, and areas (polygons), which are all vector data that consist of the user’s superimposable annotations or interpretations of the raster data (e.g., feature labels of the landscape) (Smith, 2011). Here, our photomicrographs of Nissl-stained histological material, and those of the adjacent series of immunolabeled tissues from the same subject, were captured in raster format and placed as data layers, with vector-based maps of parcellations superimposed on the photos of immunolabeled data. The simultaneous visualization of multiple data sets in registration with one another affords the investigator with novel views and juxtapositions to fuel the discovery of new spatial relationships among elements of the visualized datasets.

### 4.4 : Significance of our mapping and spatial averaging methods

We sought to capture –– graphically, within our maps –– a representation of the variance that occurs when an expert mapper transposes spatial labeling information by eye from the native tissue to the digital atlas map. In our process, by the time the mapper sets out to perform such a task, they have become familiar with the spatial relationships of the landmarks and territories in relation to their datasets. This includes an understanding of the plane of section and orientation of the tissue (what we could consider here to be a “compass sense”, or sense of direction), and the nearest-neighbor relationships of the cytoarchitectonic boundaries of the tissue in relation to the labeling patterns they are observing within it (what we could consider here to be a local “map sense”, or sense of location). This information that may be encoded by the mapper’s own brain in rough parallel to their own sense of spatial reference, or “directional sense” (Marchette et al., 2014; Chadwick & Spiers, 2014; Burte et al., 2018). Thus, when the mapper is setting out to place the labeling patterns within the atlas reference environment, they are already at a stage where the variability in placing such patterns is constrained, where their accuracy in mapping is guided by the bounds imposed by the map and compass senses. The orientation of their vantage point, together with knowledge that they are near the relevant objects in the environment, afford them the correct “directional sense”, or “field of view” away from the microscope. This is aided by their canvas already being populated as a digital atlas template, containing silhouettes of the regions appearing in their Nissl-stained tissue. Their initial task, therefore, becomes selecting from the appropriate templates and then choosing the correct general location within a single template to begin transferring labeled patterns based on a study of the Nissl series. In most instances, a simple act of placing a filled circle or traced line over the picture of a cell body or neurite, respectively, within the Adobe Illustrator environment preserves its relationship to another labeled cell body or neurite (also see Lingley et al., 2018). In fewer instances, a one-to-one drawn representation of the elements was not attempted for those labeling patterns that were very dense, but their overall trends were captured.

### 4.5 : Design features of the grid-based maps

Grid-based mapping of the brain can be traced as far back as the invention of the stereotaxic instrument by Horsley and Clarke (1909) (also see Krieg, 1975). Swanson (2000) notes that a drawing of the brain by Eustachius in the 1550s (but published in 1714), included a pair of rulers, and is a notable early example of a set of coordinates placed in relation to drawn neuroanatomical features. Grids have clearly been used in brain mapping in various ways since these early efforts, some of which may have not been documented in our scientific literature. Here, our use of grids is based on a series of specific motivations. At the most general level, we sought to establish a conceptual framework for the datasets reported by our laboratory that allowed for the recording of spatial locations in atlas-dependent and -independent formats (i.e., locations referring to specific features or entities versus locations referring to an invariant grid or field; see Rigaux et al., 2002). This was born from the “need to compute across *spatial territory*, not just *named variables for territories*, because certain [immunoreactivity] patterns cut across known territories and spill into unlabeled portions” (Khan, unpublished laboratory blog post, 21 May 2020). The specific reasons for such a need, however, are described below in greater detail.

#### 4.5.1 : Mapping elements that do not conform to cytoarchitectonic criteria

As discussed in *Section 4.2.1.3*, many neurochemical systems in the hypothalamus and zona incerta do not follow along traditional patterns of cytoarchitecture based on Nissl parcellations. Thus, the assignment of a cluster of cells with a name and Nissl-delimited cytoarchitectural boundaries, populating the atlas as a ‘part’ of a catalog of ‘parts’, does not always consider distributions that traverse or skirt these limits. A grid bypasses such constraints by allowing for its named grid region to represent everything contained within it regardless of whether the patterns fall within or outside of named brain parts. Thus, ‘extra-connectomic’ information is also represented and accounted for.

#### 4.5.2 : Generalizes space without resorting to questions of contents or phenotypes

Grid maps are a useful means to spatially account for, in a systematized fashion, the locations of labeled elements for neuroanatomical experiments. Such maps thus avoid the complexities of contextualizing location with respect to local features in the tissue, apart from macrolevel landmarks such as white matter tracts or other gross anatomical features. In contrast, while feature-based maps of the brain have undergone refinements and modifications over their history, they are often also accompanied by healthy discussion and debate concerning their validity (e.g., debates concerning cortical parcellation and compartmentalization). This is no less true for the hypothalamus and zona incerta, the nomenclature and, indeed, basic organization of which remain vigorous topics of discussion (e.g., Croizier et al., 2015; Puelles & Rubenstein, 2015; Bedont et al., 2015; Kim et al., 2020). The advantage of grid-based mapping is that the essential zoning of labeled chemoarchitecture is independent of and indifferent to varying opinions, even evidence-informed opinions, regarding the location, organization and nomenclature of the brain regions within which such chemoarchitecture resides.

That being said, there is always the option of superimposing the grid regions in our database as a digital overlay over feature-based contours for cytoarchitectonic regions, and counting immunoreactive elements within each grid region, something we find useful as a hybrid approach, where a composite map (see Fig. 12, *top row;* also Fig. 14) that includes both features and grids allows for information from both to help orient the user to accurately map elements to the appropriate grid region. This has been carried out in certain early studies of chemoarchitectural density mapping. For example, Agnati et al., (1985) reported quantitation of perikarya displaying immunoreactivity for the glucocorticoid receptor, where the counts were binned on a grid superimposed over a map of the cytoarchitectural boundaries of the locus coeruleus and paraventricular hypothalamic nucleus in young and aged male rats (see their Fig. 11 on p. 104 and Figs. 14 and 15 on pp. 107– 108 of their study, respectively; also see Zoli et al., (1993), for a similar analysis for other chemical systems). The main difference in our approach is that the grid regions we use are larger in area and indistinguishable from the areas demarcated by the stereotaxic grid already present within the atlas file in a separate layer.

Thus, the *stereotaxic grid*, which has traditionally been used to aid neuroscientists in *targeting* the brain with probes for intracranial chemical or gene-directed manipulations (Khan, 2013), is now being utilized in this instance to *map* novel chemoarchitecture. At the same time, such mapping efforts – calibrated to what is traditionally a targeting system – automatically render the newly mapped chemoarchitecture amenable to such targeting when reported this way to the greater scientific community.

#### 4.5.3 : Democratization

We also pursued the development and use of a grid-based system as a complement to feature-based mapping because there are unique advantages afforded with each method, much as there are for music referenced as standard notation or as tablature (Pillsbury, 2006, p. xvi), physical processes described in analytic (i.e., Cartesian) vs. geometric (i.e., Euclidean) notation (Feynman, 1964; scrub to 8:53–10:18), longitude measured by timekeeping or astronomical methods (Forbes, 1966; Sobel, 1995), or programs written in machine vs. assembly vs. compiler languages (Hofstadter, 1979, pp. 290–296). In our case, as in all these cases, a key advantage is the larger accessibility and democratization of data reporting that the alternative provides (see also Ramsdale et al., 2017; Voelker & Ramsdale, 2019). Specifically, little training is required to use a grid-based system to locate and indeed, even to map, labeled elements of the brain in a standardized way. In fact, apart from a basic understanding of stereotaxic coordinates and the use of a brain atlas, a user of this database could, in principle, map directly onto digital templates of the general atlas section’s outline of the brain, without the patterns of parceled brain regions in view (i.e., with the visibility of that digital overlay or data layer toggled off). Much as creators of music and art do not necessarily require formal training to pursue and successfully create and produce their work and can “approach [the work] conceptually rather than technically” (Eno, 2024), so, too, can neuroscientists untrained in formal atlas-mapping techniques use grids to localize the general position of their labeled elements in an atlas model of the brain.

Thus, the grid should potentially open the atlas to those in the scientific community who potentially seek to incorporate their datasets into a larger community model being built around the *Brain Maps* reference atlas framework (e.g., Acedo Aguilar et al., 2025a; 2025b), but who do not themselves have the time and/or expertise to rigorously map their labeled elements in the tissue to the underlying cytoarchitecture. While we encourage the use of our grid system to map their experimental probe placements or labeling patterns to BM4.0 space, we do not expect at this nascent stage any but the most discerning or motivated of community members to use what we provide here to begin such a process. We consider this as but a starting point for establishing a more accessible platform where neuroscientists can physically map their own data to the *Brain Maps* atlas and thereby contribute to a larger body of work that models structure-function relationships for the brain in a shareable and registrable manner. Such efforts, which have early proofs-of-concept (Dashti et al., 2001; Burns et al., 2006), are also now underway (Acedo Aguilar et al., 2025a; 2025b).

The benefits of such democratization are potentially numerous. For one thing, scientists who have expert training in neuroanatomical techniques outside of mapping, particularly in performing intracranial probe placements, central injections, tract-tracing, viral vector delivery, microdialysis, optogenetics, etc., can now use a relatively straightforward means to map, at a coarse resolution of a 1.0 mm × 1.0 mm stereotaxic grid region, the locations of their probe placements. Usually, the practice of reporting stereotaxic coordinates has been to state in the methods section of papers those coordinates being used to *target* a previously defined set of features, not necessarily to *report* the location of new features. If this practice gains currency – that is, of using grids to *map* in addition to *target* – then, suddenly, all probe locations mapped to grid regions by non-expert mappers can be “brought into” the spatial model of a communal atlas reference space and juxtaposed with finer neuroanatomical features mapped by experts. As proof of concept, we recently described how our mapped electrode placements from a collaborative intracranial self-stimulation study are co-spatial with prior studies of this kind mapped to the same reference (see discussion in our study: Martínez et al., 2023), allowing for future comparisons of the studies to be made lawfully with respect to identical grid regions to which they are mapped.

For a future communal atlas of the kind we have outlined here, we envision some form of content curation where tiers of quality are assigned to datasets from non-experts vs. experts. Such a content curation system ought to include lessons learned from creators and/or evaluators of similar digital resources who have identified challenges and best practices, such as the Encyclopedia of Life Project (Rotman et al., 2012). It should also be able to draw from and contribute to data sharing services that have already been established for image and atlas data, such as EBRAINS (https://ebrains.eu; see Leergaard & Bjaalie, 2022). Additionally, the use of our reporting structure (i.e., the grid-based FMRS; *see Section 3.5.4.2*) can be envisioned to serve the needs of those performing microdissections or finer samplings of brain tissue from experimental subjects (e.g., laser-capture microdissection). FMRS annotations could help anchor dissected samples of tissue and their attendant genomic, transcriptomic, proteomic or peptidomic metadata to fixed locations in atlas reference space by grid region. Such approaches allow for the contextualization of such “-omics” information with neuroanatomic information in a more precise way (Khan et al., 2018a).

Grid mapping could also be synergized with extant efforts to use computer vision or machine learning to assist in stereotaxic targeting (e.g., Arefev et al., 2021), where grid coordinates for chemoarchitecture, reported together with their spatial relation to visible landmarks during surgery, could inform surgical or intracranial methods. Conversely, the careful reporting of final intracranial injection sites intended to target the hotspots of chemoarchitecture shown in our isopleth maps could help to refine these maps (Khan, 2013). Again, there are larger challenges that need to be overcome if this is to be a reality; in particular, there remains a reportedly low commitment to best practices in reporting stereotaxic information in published studies, which limits our knowledge of the rigor and reproducibility of the experiments described in those studies (De Vloo & Nuttin, 2019). In principle, while a more straightforward mapping of a subject’s brain chemoarchitecture could be achieved by creating a subject-specific atlas (Azimi et al., 2017), the advantages of mapping the chemoarchitecture to a standardized atlas afford the scientific community a means to contextualize their data more precisely with those of others, as we have shown in this study and discussed elsewhere (Khan, 2013; Khan et al., 2018a). Indeed, there have been documented attempts to reconcile disparate spatially documented datasets that could have benefited (and still can) from standardized mapping, such as in the cerebellum (Apps & Hawkes, 2009). The benefits of bringing in data to an atlas standard have also been apparent for human studies, as exemplified by the elegant registrations of new electrode placements to the MNI human stereotactic atlas, confirming and more precisely formalizing the existence of somatosensory and motor homunculi (Roux et al., 2018; 2020; see Schieber, 2018, 2020 for commentary) originally mapped as a drawing without the underlying cortical structure by Penfield and Boldrey (1937). Moreover, as deep brain stimulation strategies continue to be developed in rat models for establishing important feasibility for the treatment of various neurological disorders, including in the hypothalamus and zona incerta (Gouveiae et al., 2023; reviewed by Zhang et al., 2024), electrode placement maps could help to facilitate comparison of stimulation targets for preclinical rat models of disease (e.g., see Martínez et al., 2023), even as the localization of electrode placements in human subjects undergoes evaluation and refinement (e.g., Hyam et al., 2015; Bower et al., 2023).

### 4.6 : Future directions

Below, we note some paths forward for possible future iterations of this database.

#### 4.6.1 : Future directions in data layers and visualization

Colors could in the future be palletized for the user’s specific needs (Hodgson, 2016), or a code of colors could universally signify a set of semiotic meanings for the grids or isopleths, forming a sort of essential visual grammar, much as they did for Mondrian’s abstract grids (Mondrian, 1987; see Weber, 2018; Deicher, 2020). Color selection could no doubt be refined to accommodate more labels, especially with advances in multiplex immunohistochemistry (e.g., Maric et al., 2021). In principle, perikarya marked on our maps by a circle glyph could also be color-coded based on cell size and analyzed for spatial patterns, as has been performed for perikarya in the cat and monkey cortex to reveal novel substructure in cytoarchitectonic patterns (Lingley et al., 2018). Our grid-based visualizations could, in principle, also encode more information by alterations in the appearance of individual tiles, much as Vasarely’s works so brilliantly achieve (e.g., see Vasarely, 1959–1961, ‘<u>Supernovae</u>’; also see Vasarely & Joray, 1970; Wade, 1978), or modify the actual parameters of the grids themselves to create depth in digital overlay space, as so adroitly achieved in other media, such as that found in the works of Christiane Feser (see Feser, 2019, ‘<u>Felder 3 (Field 3)</u>’) or Charles Biederman (see Biederman, 1939, ‘<u>Work No. 3, New York,</u> <u>1939</u>’).

Another avenue of active exploration is the migration of the *Brain Maps* reference space to a 3-D model. Such migration efforts could instantiate the use of grids in 3-D as cubic grid regions (i.e., a lattice). At the time of this writing, the *Brain Maps* reference space has been placed in coarse register with at least one extant family of 2-D atlases (the Paxinos & Watson rat brain atlases; see Khan et al., 2018b) and with an early version of the 3-D Waxholm rat atlas model (Papp et al., 2014; 2015; Kleven et al., 2023a; registration performed by Bjerke et al., 2019). We envision further, more precise integration of the *Brain Maps* space with these valuable atlas reference spaces, as well as within a future model incorporating a new “atlas brain holotype” for the *Brain Maps* reference space itself (Quintana et al., 2024; Acedo Aguilar et al., 2025a; 2025b). This migration could also eventually extend to the datasets in *Chemopleth 1.0*. To this end, the representation of data in 3-D will need refinements, particularly with respect to experimentally derived spatial data, such as injection sites and probe placements, which could be embedded within voxel-based cubic grid regions, perhaps dovetailing with recently developed pipelines for object representation within the Waxholm atlas (Blixhavn et al., 2024; Puchades/Yates et al., 2025) and incorporating established tools and approaches for 3-D mapping and visualization of labeled axonal structures (e.g., see Friedmann et al., 2020) and vasculature (e.g., Yao et al., 2021).

#### 4.6.2 : Future applications and validations

Some final considerations before closing this report are ideas concerning future applications of our database and its further validation as it continues to be populated. We first consider how maps such as these improve understanding and recall of the brain’s spatial organization. There is intriguing evidence that grids in maps facilitate greater recall of features embedded in maps (Edler et al., 2014). It would be interesting to see if this observation holds true for maps of cytoarchitecture, and a test of such a hypothesis could be conducted within a teaching laboratory setting. Work performed in our Brain Mapping & Connectomics teaching laboratory for freshman undergraduates (D’Arcy et al., 2019; Khan et al., 2021) suggests that it could be an ideal setting for such a hypothesis to be tested by student volunteers, a possibility we are now exploring. Another idea we are developing is to use the art of printmaking in this teaching laboratory to help students realize the power of digital overlays, since this technique requires careful explorations of mixed media approaches to create “layers” of data to emphasize the overlay process used digitally for published drawings of brain regions, distinct cellular morphotypes, and axonal pathways (Ramirez & Khan, 2024). Printmaking and closely aligned artistic practices such as screenprinting provide concrete examples of how overlays in register can precisely align disparate objects in space, effectively compelling the perceiver to consider them together (see for example, Rashid Johnson’s *<u>Untitled Large Mosaic, 2025</u>*; also in: Marchesano et al., 2025; pp 86–87).

As for considerations of error and validation, grid overlays could also be used to specifically bring separate mapped elements into register in a manner that minimizes specific mapping errors. Such errors often arise from differences in resolution between the overlaid elements, or from roundoff errors generated by disparities between algorithms and finite-precision mathematical operations, as has been demonstrated in geographic information systems (Magalhães et al., 2015). The overall extent of error in mapping using digital systems, more generally, has been a subject of much discussion in cartography (e.g., Chrisman, 1982) and future work will help formalize the sources of error for atlas-based transfer of anatomical information from photomicrographs of histologically processed tissue sections (see the cogent discussion of this topic by Simmons and Swanson, 2009; also see the resource provided by Fuglstad et al., 2023). Aggregation of such data in the form of spatially averaged choropleth and isopleth maps merit additional considerations of error (e.g., MacEachren, 1982; 1985; Sun et al., 2015) which require further exploration for atlas-based mapping. Error correction and the speed of manual mapping of elements could, in principle, be improved or augmented by incorporating visual records of expert mappers’ physical movements as they recognize Nissl cytoarchitectonic patterns and perform manual mapping, the feasibility of which has been recently demonstrated for histological tissue sets reviewed by clinical pathologists who volunteer to have their eyes movements captured for use by machine learning algorithms (Nan et al., 2025). Along these lines, error correction of the maps compels us to consider, as its counterpart, validations of the ground truth underlying our mapping techniques, which could be more formalized in the future, since it is clear that “ground truth” is an idealistic goal that one moves toward rather than a reality that exists without any possibility of further refinement (Sanchez Castro et al., 2006).

Finally, we sought to create a database resource that complies to FAIR standards (Wilkinson et al., 2016; Sandström et al., 2022; Martone, 2024; *discussed in Section 3.1*). Towards that end, as we created this resource, we kept guidelines in mind that have been proposed for the development of biological databases (Helmy et al., 2016) and also anticipate that *Chemopleth 1.0* could help contribute to a standard markup language already developed at one scale for the *Brain Maps* atlas environment (Brown & Swanson, 2013), and perhaps for others in the future.

### 4.7 : Concluding Remarks

In this report, we have provided a description of a spatial database, called *Chemopleth 1.0* (available at: https://doi.org/10.5281/zenodo.15788189), which provides freely accessible *Brain* Maps 4*.0* digital atlas maps of five neurochemical systems in the rat *hypothalamus (Kuhlenbeck, 1927)* and *zona incerta (>1840)*, brain regions critically involved in controlling many survival functions. Mapping the spatial organization of cells and fibers in the brain is a fundamental challenge in neuroscience (Swanson, 2000; 2007; Milligan et al., 2019; Newmaster et al., 2022). Now in its fourth edition (Swanson, 2018), the *Brain Maps* atlas provides a rigorous anatomical reference for the rat brain and is a valuable resource for the neuroscience community, alongside other excellent brain atlases for the rat (e.g., Paxinos & Watson, 2014; Papp et al., 2014; 2015; Kleven et al., 2023a), which have been placed in basic register with the *Brain Maps* framework (Khan et al., 2018b; Bjerke et al., 2019). Our efforts to create a spatial database at mesoscale resolution complements efforts to create databases at macroscales based on standardized brain atlases (e.g., Nowinski et al., 1997; 2000; also see Nowinski, 2023), and are meant to be used for the same purpose as those are, particularly with respect to stereotaxic targeting (e.g., see Nowinski et al., 2000; also see the finer-scale mouse stereotaxic atlas of Feng et al., 2025). The work also complements efforts for mouse brain to create interactive frameworks to synergize disparate datasets at cellular resolution (Fürth et al., 2018; Y. Chen et al., 2019).

Ultimately, we envision that future numbered versions of *Chemopleth 1.0* will: (1) contain greater amounts of chemoarchitectural data registered to our existing collection of mapped datasets, adding to the database, for example, atlas maps currently being finalized for additional neuropeptides (e.g., Delgado et al., 2024); (2) incorporate more registered legacy datasets from the published literature (e.g., Watts et al., 1999; Swanson et al., 2005; Hahn, 2010); and (3) include *Brain Maps* base maps that are created for future revised versions of the core atlas. Apart from these naturally progressing improvements, we are eyeing the artificial intelligence (AI) landscape for geospatial/ archaeological analysis (e.g., the newly emerging field of GeoAI; see Kang, 2020; Kang et al., 2024; see Casini et al., 2023, for archaeological application) to help develop workflows that allow diverse spatial locations of corresponding space on digital overlays to be analyzed for cause-effect biological relationships and inferences drawn about possible interactions, as conceptualized over a decade ago by our laboratory (Fig. 8 in Khan, 2013). This will allow for multiscale integration of experimental metadata and would address some issues plaguing top-down and bottom-up approaches (D’Angelo & Jirsa, 2022). They will also bring quickly to the fore the larger challenge of making sense of all spatially registered data within one mapped location, finally shifting our attention in neuroanatomy from data gathering to data intelligence, as we finally “face the mother lode” (Gabriel, 2023a; 2023b): evidence collected from communal data, and finally organized as shared information (e.g., Dashti et al., 2001; Burns et al., 2003; 2006; Khan et al., 2006), waiting to be harnessed for medical treatments, drug discovery, multiscale views of the nervous system, and ultimately, perhaps a deeper, better-informed view of ourselves and our close relatives.

## Acknowledgments

### Funding

This work was supported by grants awarded to AMK from the National Institute of General Medical Sciences (NIGMS) of the National Institutes of Health (NIH) (GM109817 and GM127251) and a grant from the UTEP Research & Innovation Office to the Interdisciplinary Group for Neuroscience Investigation, Training, and Education (IGNITE) investigators (L. E. O’Dell, A. M. Khan, I. A. Mendez, B. Cushing, S. Iñiguez). Work performed in the Brain Mapping & Connectomics (BMC) Undergraduate Teaching Laboratory was conducted within UTEP PERSIST (Program to Educate and Retain Students in STEM Tracks), a training program funded by Howard Hughes Medical Institute grant 52008125 (PI: S. B. Aley, Co-PIs: L. Echegoyen, A. M. Khan, D. Villagrán, E. Greenbaum). VIN and AA were supported by Doctoral Excellence Fellowships. VIN, ART, and DS are fellows of the UTEP RISE (Research Initiative for Scientific Enhancement) program (NIGMS; R25GM069621; PI: R. Aguilera). AA was also supported by Dodson Research Grants from the UTEP Graduate School and by travel grants from the UTEP Graduate School and the College of Science. This work was also supported by the Border Biomedical Research Center, funded by the National Institute on Minority Health and Health Disparities (NIMHD; 2U54MD007592; PI: R. A. Kirken).

### Colleagues

The authors thank Patrick Hof, Larry W. Swanson, and the late Anthony N. van den Pol for their support and encouragement of this project. We also thank Suzana Herculano-Houzel and Melissa J. Chee for their feedback on portions of this study. We acknowledge Claire E. Wells, Christina E. D’Arcy, Briana Pinales, Nicole Dominguez, Richard H. Thompson, Georgina A. Lean, Kristen Pennington, Alexa Escapita, Paola Rojas, and our BMC students for early immunofluorescence efforts for this project. We are grateful to members of the UTEP Systems Neuroscience Laboratory, especially Anais Martinez, Christina E. D’Arcy and Kenichiro Negishi for valuable discussions. AMK thanks the following colleagues for their service on the dissertation committees of VIN, RP and AA: [VIN] Ian A. Mendez, Andrew M. Poulos, Alexander Friedman, and Manuel Miranda; [EP] Germán Rosas-Acosta, Vanessa H. Routh, Edward Castañeda, and Bruce S. Cushing; [AA] Jorge A. Muñoz. AMK expresses his deep gratitude to Vivian-Lee Nyitray for introducing to him Zen aesthetic and tradition, which have enriched theoretical underpinnings of this project.

### Writing spaces, art installations, live performances

AMK acknowledges several spaces, art installations and live performances influencing the ideas presented in this study including sites where notes for this manuscript were written over several years. These include *Nick Cave:Forothermore* by Nick Cave at the Museum of Contemporary Art, Chicago (Sep 2022); *Maya: The Exhibition* at the California Science Center (Jul 2023); the Robert and Arlene Kogod Courtyard at the Smithsonian American Art Museum/National Portrait Gallery, Washington, D.C. (Jun 2012; Nov 2023; Mar and Apr 2024); *i/o – The Tour*, concert performed by Peter Gabriel at the Kia Forum, Inglewood, California (Oct 2023); *Jennifer Guidi:And so it is* by Jennifer Guidi at the Orange County Museum of Art, Costa Mesa, California (Dec 2023); the Zen Court at the Japanese Garden, The Huntington Library and Botanic Gardens, San Marino, California (Feb 2024); *Where We Belong*, a play performed by Madeline Sayet at the Folger Theatre at the Folger Shakespeare Library, Washington, D.C. (Mar 2024); the Japanese Friendship Garden of Phoenix, Phoenix, Arizona (Jun 2024); *Eno*, a generative documentary directed by Gary Hustwit, screened at the Texas Theatre, Dallas, Texas (Jul 2024); the collections at the Dallas Museum of Art (Jul 2024) and the Museum of Fine Arts, Boston (Jul 2024); a live performance of Frederic Mendelssohn’s Octet in E major by the Camerata del Sol chamber orchestra at the Women’s Club of El Paso (Apr 2025), and *Brand X Editions: Innovation in Screenprinting* at The Philadelphia Museum of Art (Oct 2025).

### Trademarks and credits

Adobe Illustrator and Adobe Photoshop are either registered trademarks or trademarks of Adobe in the United States and/or other countries.

### Resource sharing

#### Data availability

Digital maps are available in Zenodo as individual atlas files in SVG format that can be opened (with layers preserved) in Adobe Illustrator or in Inkscape: https://doi.org/10.5281/zenodo.15788189. Note that one known issue for Inkscape at the time of this writing is that the syntax of layer names are slightly altered during import of the downloaded files and require manual correction. <u>Code availability</u>: All code is available in <u>GitHub</u>.

## Supporting information

Supplemental Methods and References

## Author Contributions

*Design and conceptualization of project*: AMK, VIN, AA, SB.

*Mapping of original histological data*: VIN, EP, ART, DS

*Mapping and registration of legacy data*: VIN, ART, DS

*Conceptualization of isopleth and choropleth maps*: AMK, AA, SB

*Conceptualization of grid-based annotation*: AMK

*Coding, tool development, and documentation*: AA with input from AMK, SB, OF

*Preparation of datasets within Chemopleth 1.0*: VIN, AA, AMK

*Grid-based tabulations*: SB

*Undergraduate supervision*: VIN, EP

*Graduate student supervision*:

Chair of VIN’s and EP’s dissertation committees: AMK

Chair and Co-Chair, respectively, of AA’s dissertation committee: OF, AMK

*Project Management*: AMK

*Funding acquisition*: AMK

*Conference presentations*: AA, AMK, VIN, OF

*Manuscript writing*: AMK, VIN, AA with feedback from the other co-authors

## Abbreviations

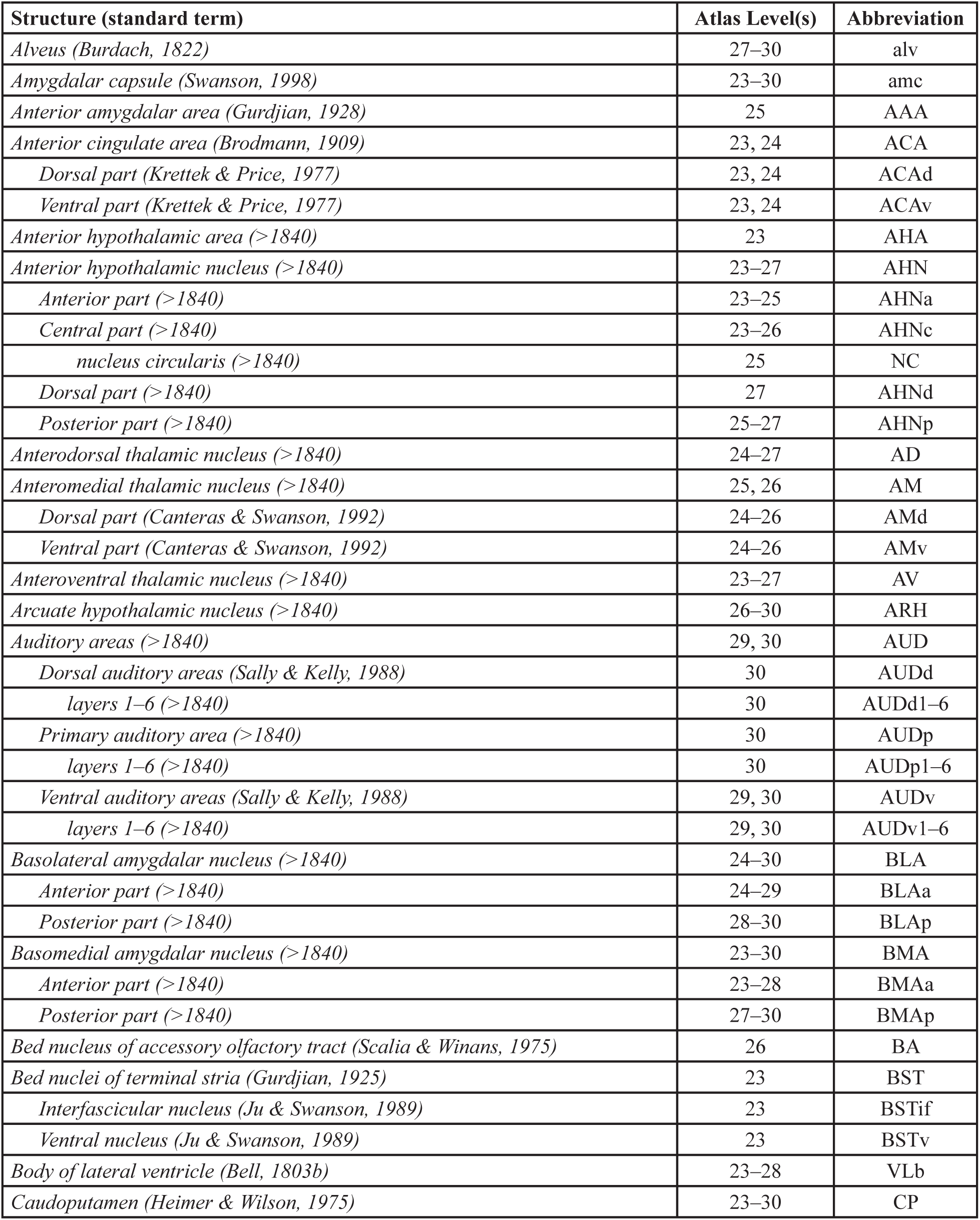
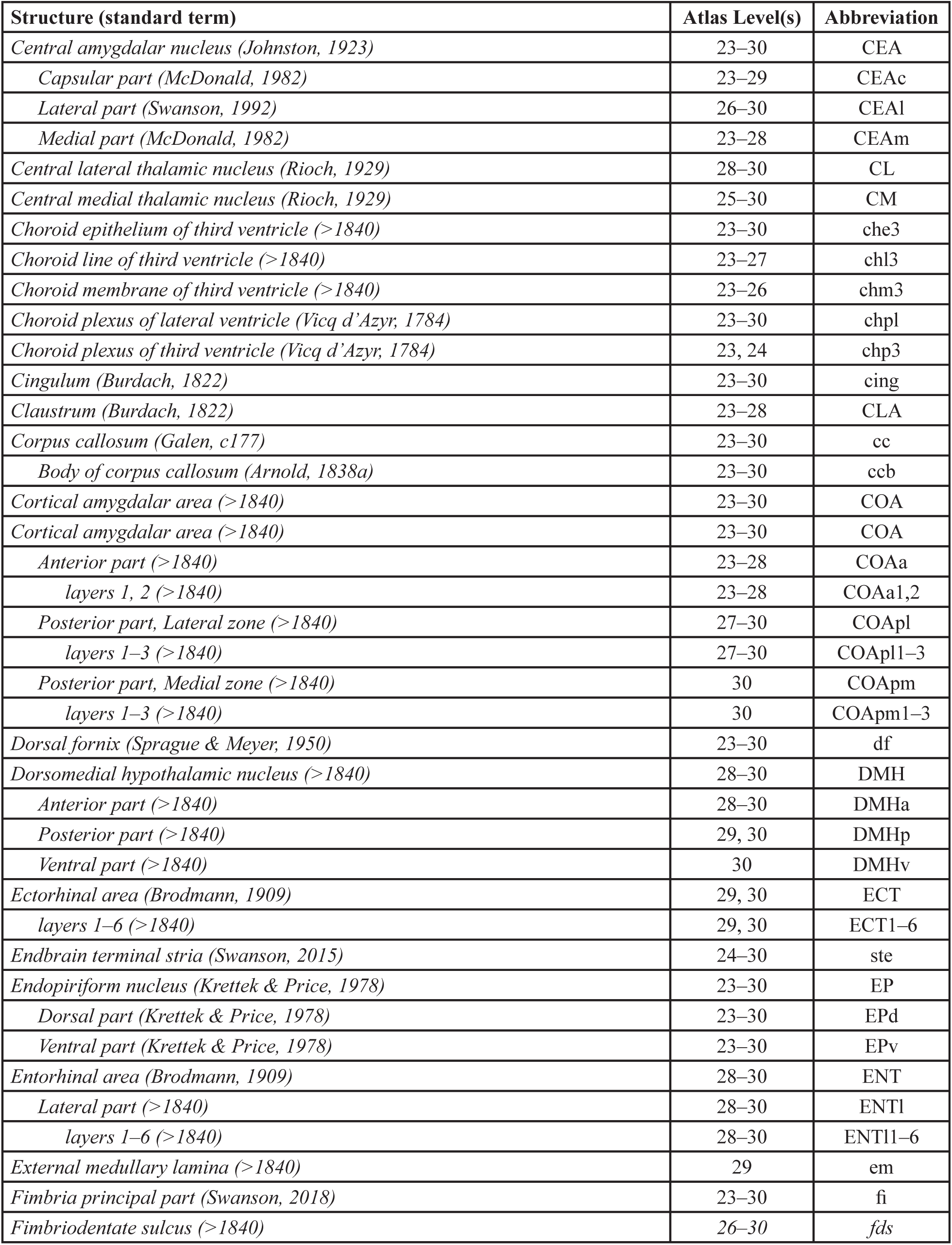
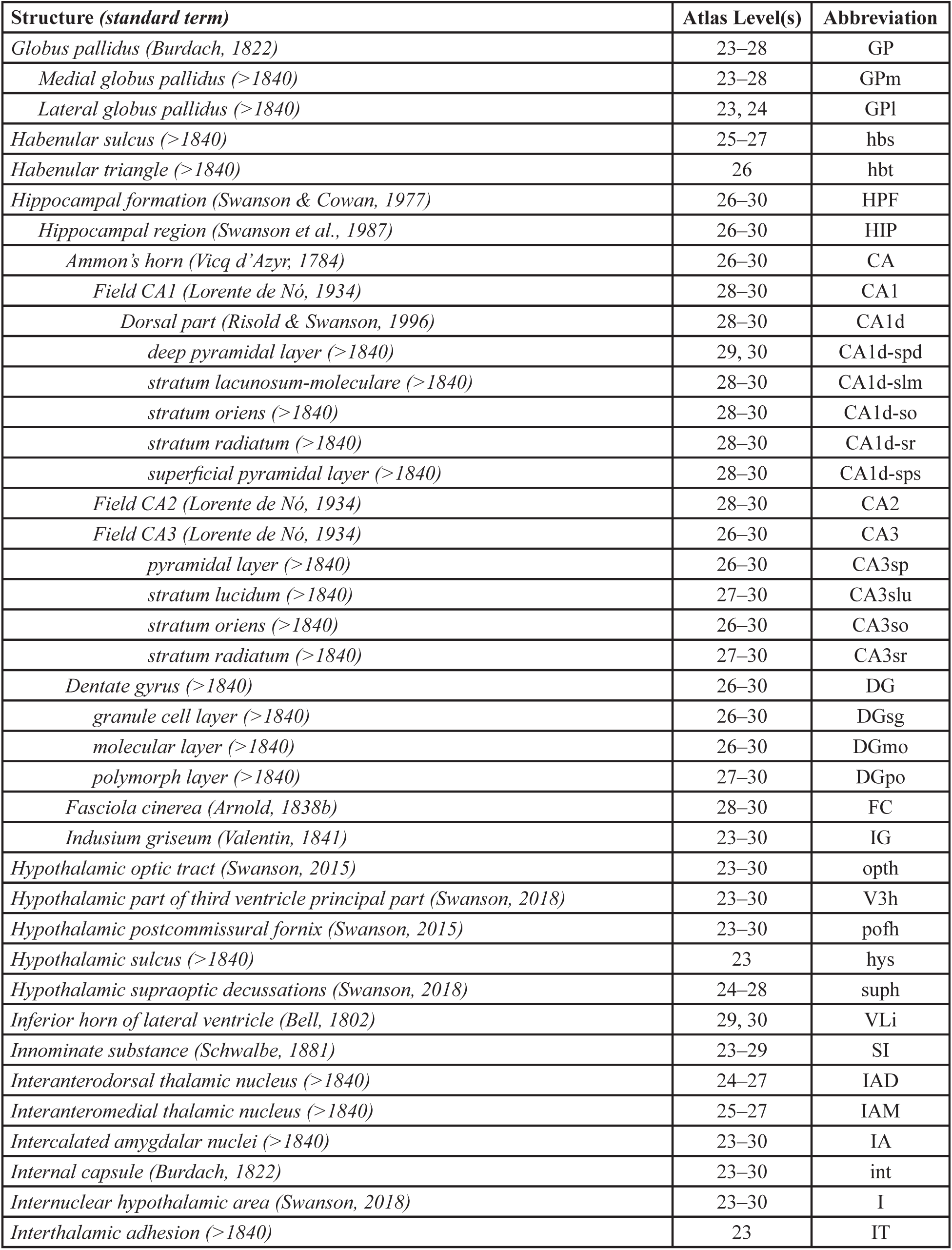
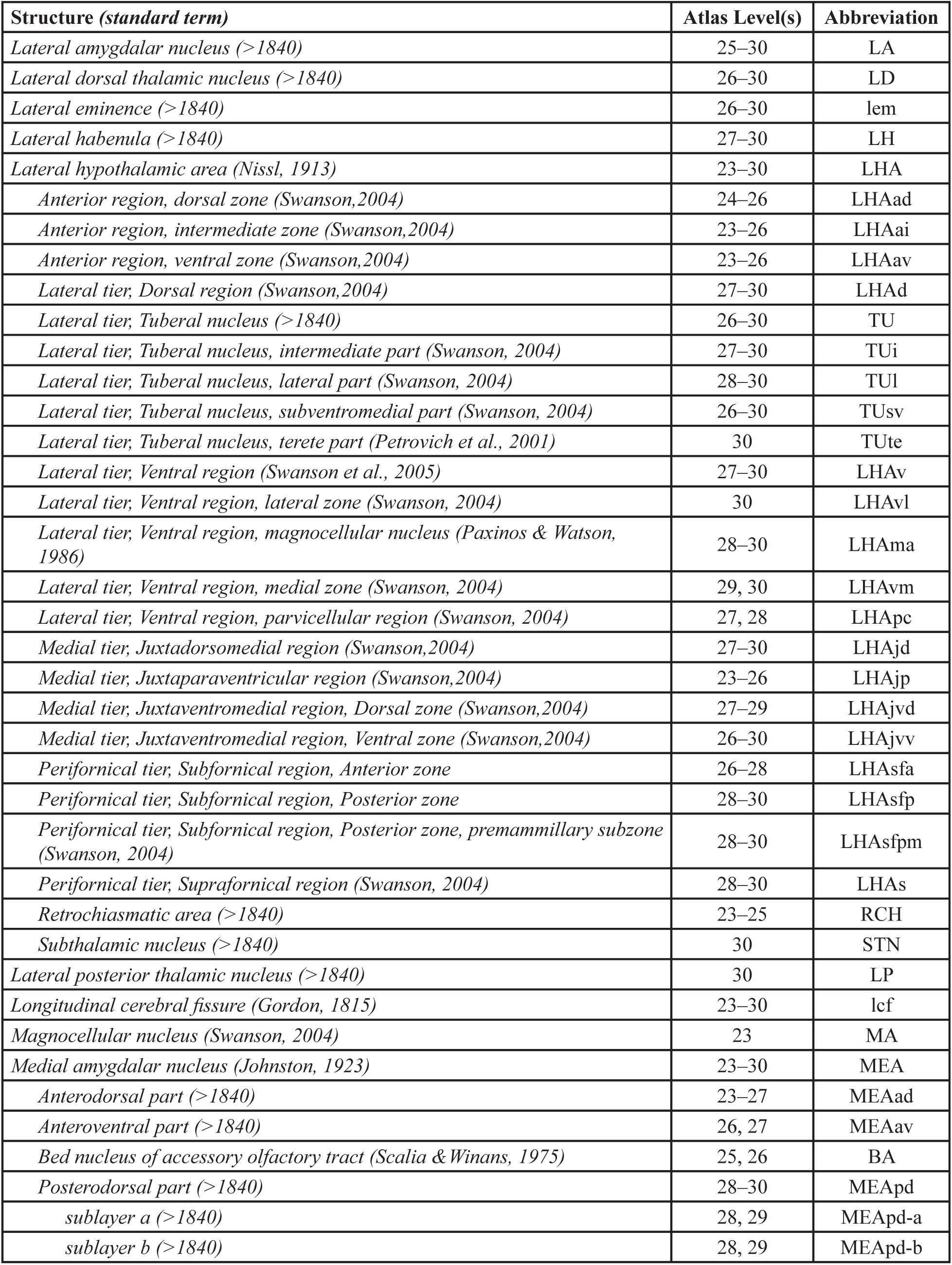
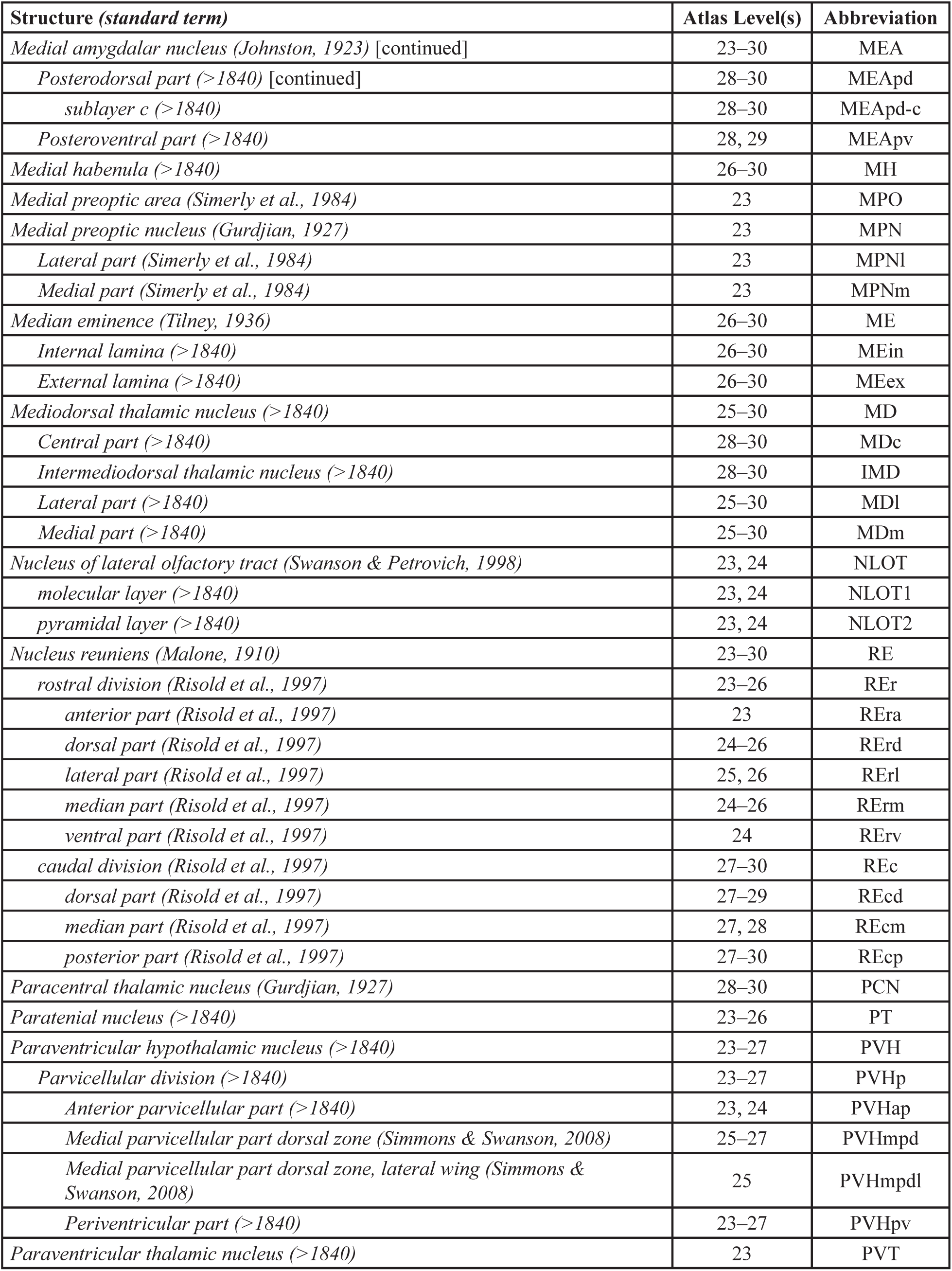
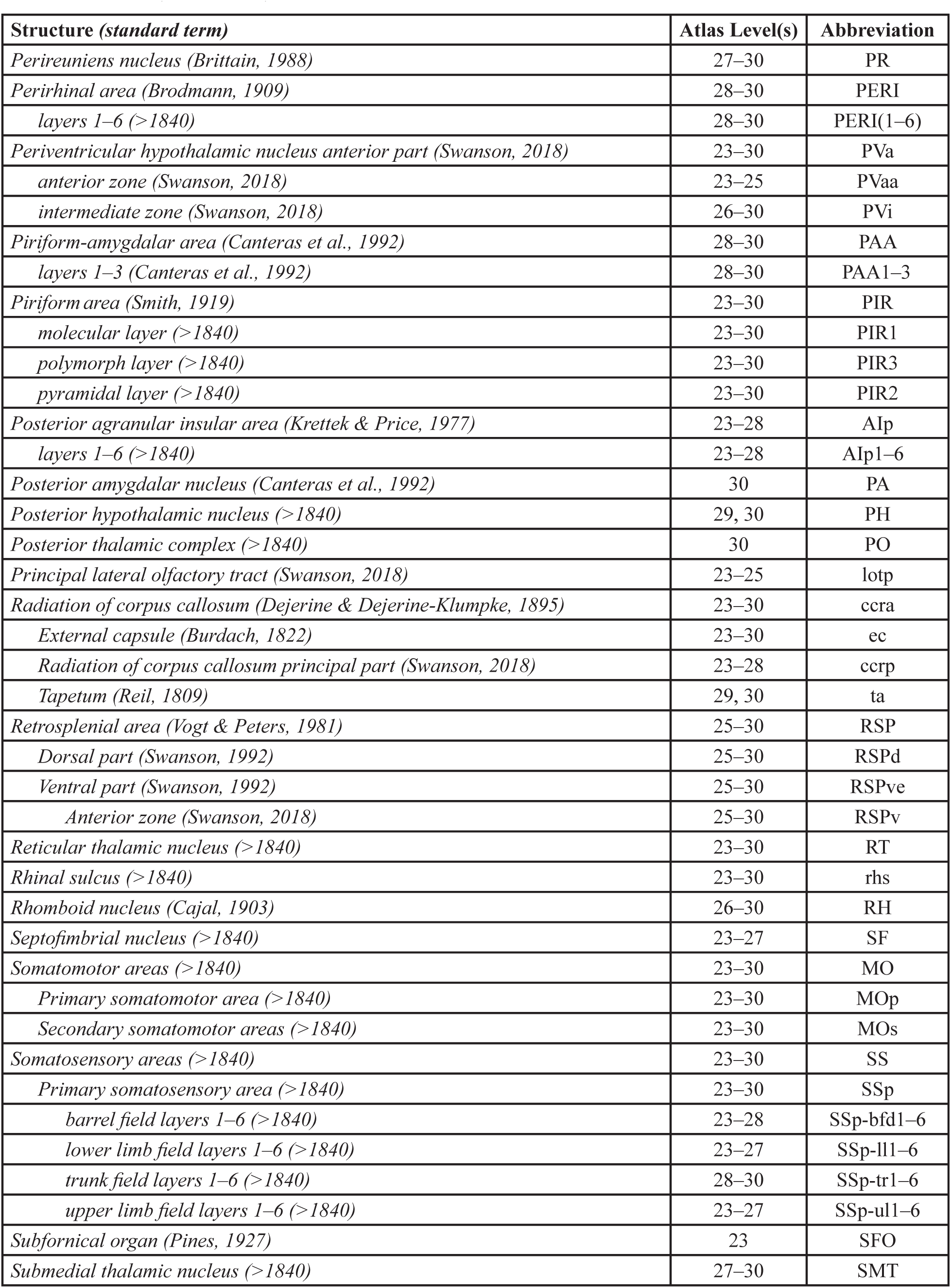
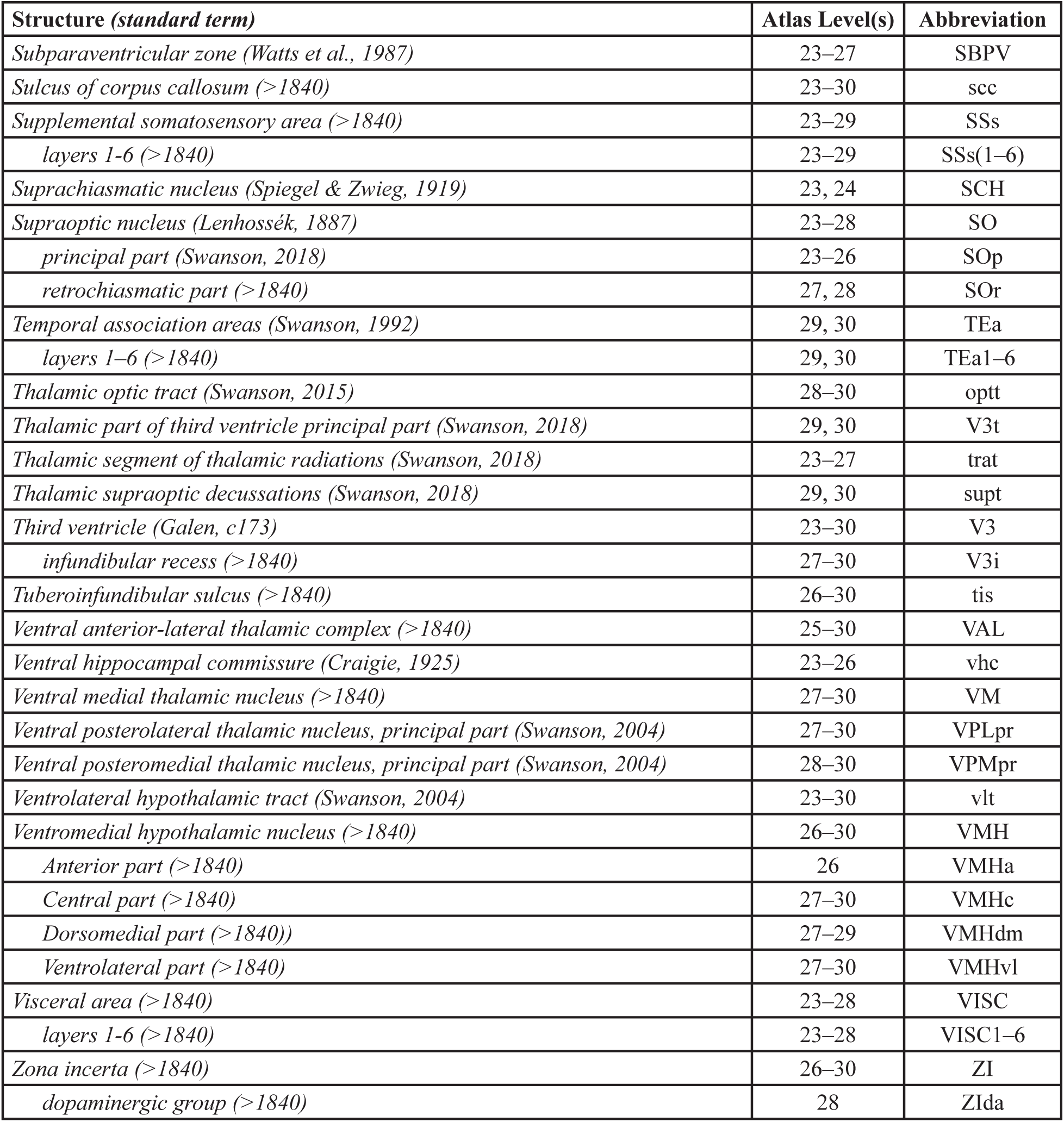

| Structure (standard term) | Atlas Level(s) | Abbreviation |
| --- | --- | --- |
| *Alveus (Burdach, 1822)* | 27–30 | alv |
| *Amygdalar capsule (Swanson, 1998)* | 23–30 | amc |
| *Anterior amygdalar area (Gurdjian, 1928)* | 25 | AAA |
| *Anterior cingulate area (Brodmann, 1909)* | 23, 24 | ACA |
| *Dorsal part (Krettek & Price, 1977)* | 23, 24 | ACAd |
| *Ventral part (Krettek & Price, 1977)* | 23, 24 | ACAv |
| *Anterior hypothalamic area (>1840)* | 23 | AHA |
| *Anterior hypothalamic nucleus (>1840)* | 23–27 | AHN |
| *Anterior part (>1840)* | 23–25 | AHNa |
| *Central part (>1840)* | 23–26 | AHNc |
| *nucleus circularis (>1840)* | 25 | NC |
| *Dorsal part (>1840)* | 27 | AHNd |
| *Posterior part (>1840)* | 25–27 | AHNp |
| *Anterodorsal thalamic nucleus (>1840)* | 24–27 | AD |
| *Anteromedial thalamic nucleus (>1840)* | 25, 26 | AM |
| *Dorsal part (Canteras & Swanson, 1992)* | 24–26 | AMd |
| *Ventral part (Canteras & Swanson, 1992)* | 24–26 | AMv |
| *Anteroventral thalamic nucleus (>1840)* | 23–27 | AV |
| *Arcuate hypothalamic nucleus (>1840)* | 26–30 | ARH |
| *Auditory areas (>1840)* | 29, 30 | AUD |
| *Dorsal auditory areas (Sally & Kelly, 1988)* | 30 | AUDd |
| *layers 1–6 (>1840)* | 30 | AUDd1–6 |
| *Primary auditory area (>1840)* | 30 | AUDp |
| *layers 1–6 (>1840)* | 30 | AUDp1–6 |
| *Ventral auditory areas (Sally & Kelly, 1988)* | 29, 30 | AUDv |
| *layers 1–6 (>1840)* | 29, 30 | AUDv1–6 |
| *Basolateral amygdalar nucleus (>1840)* | 24–30 | BLA |
| *Anterior part (>1840)* | 24–29 | BLAa |
| *Posterior part (>1840)* | 28–30 | BLAp |
| *Basomedial amygdalar nucleus (>1840)* | 23–30 | BMA |
| *Anterior part (>1840)* | 23–28 | BMAa |
| *Posterior part (>1840)* | 27–30 | BMAp |
| *Bed nucleus of accessory olfactory tract (Scalia & Winans, 1975)* | 26 | BA |
| *Bed nuclei of terminal stria (Gurdjian, 1925)* | 23 | BST |
| *Interfascicular nucleus (Ju & Swanson, 1989)* | 23 | BSTif |
| *Ventral nucleus (Ju & Swanson, 1989)* | 23 | BSTv |
| *Body of lateral ventricle (Bell, 1803b)* | 23–28 | VLb |
| *Caudoputamen (Heimer & Wilson, 1975)* | 23–30 | CP |
| *Central amygdalar nucleus (Johnston, 1923)* | 23–30 | CEA |
| *Capsular part (McDonald, 1982)* | 23–29 | CEAc |
| *Lateral part (Swanson, 1992)* | 26–30 | CEAl |
| *Medial part (McDonald, 1982)* | 23–28 | CEAm |
| *Central lateral thalamic nucleus (Rioch, 1929)* | 28–30 | CL |
| *Central medial thalamic nucleus (Rioch, 1929)* | 25–30 | CM |
| *Choroid epithelium of third ventricle (>1840)* | 23–30 | che3 |
| *Choroid line of third ventricle (>1840)* | 23–27 | chl3 |
| *Choroid membrane of third ventricle (>1840)* | 23–26 | chm3 |
| *Choroid plexus of lateral ventricle (Vicq d'Azyr, 1784)* | 23–30 | chpl |
| *Choroid plexus of third ventricle (Vicq d'Azyr, 1784)* | 23, 24 | chp3 |
| *Cingulum (Burdach, 1822)* | 23–30 | cing |
| *Clastrum (Burdach, 1822)* | 23–28 | CLA |
| *Corpus callosum (Galen, c177)* | 23–30 | cc |
| *Body of corpus callosum (Arnold, 1838a)* | 23–30 | ccb |
| *Cortical amygdalar area (>1840)* | 23–30 | COA |
| *Cortical amygdalar area (>1840)* | 23–30 | COA |
| *Anterior part (>1840)* | 23–28 | COAa |
| *layers 1, 2 (>1840)* | 23–28 | COAa1,2 |
| *Posterior part, Lateral zone (>1840)* | 27–30 | COApl |
| *layers 1–3 (>1840)* | 27–30 | COApl1–3 |
| *Posterior part, Medial zone (>1840)* | 30 | COApm |
| *layers 1–3 (>1840)* | 30 | COApm1–3 |
| *Dorsal fornix (Sprague & Meyer, 1950)* | 23–30 | df |
| *Dorsomedial hypothalamic nucleus (>1840)* | 28–30 | DMH |
| *Anterior part (>1840)* | 28–30 | DMHa |
| *Posterior part (>1840)* | 29, 30 | DMHp |
| *Ventral part (>1840)* | 30 | DMHv |
| *Ectorhinal area (Brodman, 1909)* | 29, 30 | ECT |
| *layers 1–6 (>1840)* | 29, 30 | ECT1–6 |
| *Endbrain terminal stria (Swanson, 2015)* | 24–30 | ste |
| *Endopiriform nucleus (Krettek & Price, 1978)* | 23–30 | EP |
| *Dorsal part (Krettek & Price, 1978)* | 23–30 | EPd |
| *Ventral part (Krettek & Price, 1978)* | 23–30 | EPv |
| *Entorhinal area (Brodman, 1909)* | 28–30 | ENT |
| *Lateral part (>1840)* | 28–30 | ENTl |
| *layers 1–6 (>1840)* | 28–30 | ENTl1–6 |
| *External medullary lamina (>1840)* | 29 | em |
| *Fimbria principal part (Swanson, 2018)* | 23–30 | fi |
| *Fimbriodentate sulcus (>1840)* | 26–30 | fds |
| *Globus pallidus* (Burdach, 1822) | 23–28 | GP |
| *Medial globus pallidus* (>1840) | 23–28 | GPm |
| *Lateral globus pallidus* (>1840) | 23, 24 | GPl |
| *Habenular sulcus* (>1840) | 25–27 | hbs |
| *Habenular triangle* (>1840) | 26 | hbt |
| *Hippocampal formation* (Swanson & Cowan, 1977) | 26–30 | HPF |
| *Hippocampal region* (Swanson et al., 1987) | 26–30 | HIP |
| *Ammon's horn* (Vicq d'Azyr, 1784) | 26–30 | CA |
| *Field CA1* (Lorente de Nó, 1934) | 28–30 | CA1 |
| *Dorsal part* (Risold & Swanson, 1996) | 28–30 | CA1d |
| *deep pyramidal layer* (>1840) | 29, 30 | CA1d-spd |
| *stratum lacunosum-moleculare* (>1840) | 28–30 | CA1d-slm |
| *stratum oriens* (>1840) | 28–30 | CA1d-so |
| *stratum radiatum* (>1840) | 28–30 | CA1d-sr |
| *superficial pyramidal layer* (>1840) | 28–30 | CA1d-sps |
| *Field CA2* (Lorente de Nó, 1934) | 28–30 | CA2 |
| *Field CA3* (Lorente de Nó, 1934) | 26–30 | CA3 |
| *pyramidal layer* (>1840) | 26–30 | CA3sp |
| *stratum lucidum* (>1840) | 27–30 | CA3slu |
| *stratum oriens* (>1840) | 26–30 | CA3so |
| *stratum radiatum* (>1840) | 27–30 | CA3sr |
| *Dentate gyrus* (>1840) | 26–30 | DG |
| *granule cell layer* (>1840) | 26–30 | DGsg |
| *molecular layer* (>1840) | 26–30 | DGmo |
| *polymorph layer* (>1840) | 27–30 | DGpo |
| *Fasciola cinerea* (Arnold, 1838b) | 28–30 | FC |
| *Indusium griseum* (Valentin, 1841) | 23–30 | IG |
| *Hypothalamic optic tract* (Swanson, 2015) | 23–30 | oph |
| *Hypothalamic part of third ventricle principal part* (Swanson, 2018) | 23–30 | V3h |
| *Hypothalamic postcommissural fornix* (Swanson, 2015) | 23–30 | poth |
| *Hypothalamic sulcus* (>1840) | 23 | hys |
| *Hypothalamic supraoptic decussations* (Swanson, 2018) | 24–28 | suph |
| *Inferior horn of lateral ventricle* (Bell, 1802) | 29, 30 | VLi |
| *Innominate substance* (Schwalbe, 1881) | 23–29 | SI |
| *Interanterodorsal thalamic nucleus* (>1840) | 24–27 | IAD |
| *Interanteromedial thalamic nucleus* (>1840) | 25–27 | IAM |
| *Intercalated amygdalar nuclei* (>1840) | 23–30 | IA |
| *Internal capsule* (Burdach, 1822) | 23–30 | int |
| *Internuclear hypothalamic area* (Swanson, 2018) | 23–30 | I |
| *Interthalamic adhesion* (>1840) | 23 | IT |

| Structure (*standard term*) | Atlas Level(s) | Abbreviation |
| --- | --- | --- |
| *Lateral amygdalar nucleus (>1840)* | 25–30 | LA |
| *Lateral dorsal thalamic nucleus (>1840)* | 26–30 | LD |
| *Lateral eminence (>1840)* | 26–30 | lem |
| *Lateral habenula (>1840)* | 27–30 | LH |
| *Lateral hypothalamic area (Nissl, 1913)* | 23–30 | LHA |
| *Anterior region, dorsal zone (Swanson, 2004)* | 24–26 | LHAad |
| *Anterior region, intermediate zone (Swanson, 2004)* | 23–26 | LHAai |
| *Anterior region, ventral zone (Swanson, 2004)* | 23–26 | LHAav |
| *Lateral tier, Dorsal region (Swanson, 2004)* | 27–30 | LHAd |
| *Lateral tier, Tuberal nucleus (>1840)* | 26–30 | TU |
| *Lateral tier, Tuberal nucleus, intermediate part (Swanson, 2004)* | 27–30 | TUi |
| *Lateral tier, Tuberal nucleus, lateral part (Swanson, 2004)* | 28–30 | TUI |
| *Lateral tier, Tuberal nucleus, subventromedial part (Swanson, 2004)* | 26–30 | TUsv |
| *Lateral tier, Tuberal nucleus, terete part (Petrovich et al., 2001)* | 30 | TUte |
| *Lateral tier, Ventral region (Swanson et al., 2005)* | 27–30 | LHA<sub>v</sub> |
| *Lateral tier, Ventral region, lateral zone (Swanson, 2004)* | 30 | LHA<sub>vl</sub> |
| *Lateral tier, Ventral region, magnocellular nucleus (Paxinos & Watson, 1986)* | 28–30 | LHA<sub>ma</sub> |
| *Lateral tier, Ventral region, medial zone (Swanson, 2004)* | 29, 30 | LHA<sub>vm</sub> |
| *Lateral tier, Ventral region, parvicellular region (Swanson, 2004)* | 27, 28 | LHA<sub>pc</sub> |
| *Medial tier, Juxtadorsomedial region (Swanson, 2004)* | 27–30 | LHA<sub>jd</sub> |
| *Medial tier, Juxtaparaventricular region (Swanson, 2004)* | 23–26 | LHA<sub>jp</sub> |
| *Medial tier, Juxtaventromedial region, Dorsal zone (Swanson, 2004)* | 27–29 | LHA<sub>jvd</sub> |
| *Medial tier, Juxtaventromedial region, Ventral zone (Swanson, 2004)* | 26–30 | LHA<sub>jvv</sub> |
| *Perifornical tier, Subfornical region, Anterior zone* | 26–28 | LHA<sub>sfa</sub> |
| *Perifornical tier, Subfornical region, Posterior zone* | 28–30 | LHA<sub>sfp</sub> |
| *Perifornical tier, Subfornical region, Posterior zone, premammillary subzone (Swanson, 2004)* | 28–30 | LHA<sub>sfp<sub>m</sub></sub> |
| *Perifornical tier, Supraforfornical region (Swanson, 2004)* | 28–30 | LHA<sub>s</sub> |
| *Retrochiasmatic area (>1840)* | 23–25 | RCH |
| *Subthalamic nucleus (>1840)* | 30 | STN |
| *Lateral posterior thalamic nucleus (>1840)* | 30 | LP |
| *Longitudinal cerebral fissure (Gordon, 1815)* | 23–30 | lcf |
| *Magnocellular nucleus (Swanson, 2004)* | 23 | MA |
| *Medial amygdalar nucleus (Johnston, 1923)* | 23–30 | MEA |
| *Anterodorsal part (>1840)* | 23–27 | MEA<sub>ad</sub> |
| *Anteroventral part (>1840)* | 26, 27 | MEA<sub>av</sub> |
| *Bed nucleus of accessory olfactory tract (Scalia & Winans, 1975)* | 25, 26 | BA |
| *Posterodorsal part (>1840)* | 28–30 | MEA<sub>pd</sub> |
| *sublayer a (>1840)* | 28, 29 | MEA<sub>pd-a</sub> |
| *sublayer b (>1840)* | 28, 29 | MEA<sub>pd-b</sub> |
| *Medial amygdalar nucleus (Johnston, 1923) [continued]* | 23–30 | MEA |
| *Posterodorsal part (>1840) [continued]* | 28–30 | MEApd |
| *sublayer c (>1840)* | 28–30 | MEApd-c |
| *Posteroventral part (>1840)* | 28, 29 | MEApv |
| *Medial habenula (>1840)* | 26–30 | MH |
| *Medial preoptic area (Simerly et al., 1984)* | 23 | MPO |
| *Medial preoptic nucleus (Gurdjian, 1927)* | 23 | MPN |
| *Lateral part (Simerly et al., 1984)* | 23 | MPNI |
| *Medial part (Simerly et al., 1984)* | 23 | MPNm |
| *Median eminence (Tilney, 1936)* | 26–30 | ME |
| *Internal lamina (>1840)* | 26–30 | MEin |
| *External lamina (>1840)* | 26–30 | MEex |
| *Mediodorsal thalamic nucleus (>1840)* | 25–30 | MD |
| *Central part (>1840)* | 28–30 | MDc |
| *Intermediodorsal thalamic nucleus (>1840)* | 28–30 | IMD |
| *Lateral part (>1840)* | 25–30 | MDl |
| *Medial part (>1840)* | 25–30 | MDm |
| *Nucleus of lateral olfactory tract (Swanson & Petrovich, 1998)* | 23, 24 | NLOT |
| *molecular layer (>1840)* | 23, 24 | NLOT1 |
| *pyramidal layer (>1840)* | 23, 24 | NLOT2 |
| *Nucleus reuniens (Malone, 1910)* | 23–30 | RE |
| *rostral division (Risold et al., 1997)* | 23–26 | REr |
| *anterior part (Risold et al., 1997)* | 23 | REra |
| *dorsal part (Risold et al., 1997)* | 24–26 | RErd |
| *lateral part (Risold et al., 1997)* | 25, 26 | RErl |
| *median part (Risold et al., 1997)* | 24–26 | RErm |
| *ventral part (Risold et al., 1997)* | 24 | RErv |
| *caudal division (Risold et al., 1997)* | 27–30 | REc |
| *dorsal part (Risold et al., 1997)* | 27–29 | REcd |
| *median part (Risold et al., 1997)* | 27, 28 | REcm |
| *posterior part (Risold et al., 1997)* | 27–30 | REcp |
| *Paracentral thalamic nucleus (Gurdjian, 1927)* | 28–30 | PCN |
| *Paratenial nucleus (>1840)* | 23–26 | PT |
| *Paraventricular hypothalamic nucleus (>1840)* | 23–27 | PVH |
| *Parvicellular division (>1840)* | 23–27 | PVHp |
| *Anterior parvicellular part (>1840)* | 23, 24 | PVHap |
| *Medial parvicellular part dorsal zone (Simmons & Swanson, 2008)* | 25–27 | PVHmpd |
| *Medial parvicellular part dorsal zone, lateral wing (Simmons & Swanson, 2008)* | 25 | PVHmpdl |
| *Periventricular part (>1840)* | 23–27 | PVHpv |
| *Paraventricular thalamic nucleus (>1840)* | 23 | PVT |
| *Perireuniens nucleus (Brittain, 1988)* | 27–30 | PR |
| *Perirhinal area (Brodmann, 1909)* | 28–30 | PERI |
| *layers 1–6 (>1840)* | 28–30 | PERI(1–6) |
| *Periventricular hypothalamic nucleus anterior part (Swanson, 2018)* | 23–30 | PVa |
| *anterior zone (Swanson, 2018)* | 23–25 | PVaa |
| *intermediate zone (Swanson, 2018)* | 26–30 | PVi |
| *Piriform-amygdalar area (Canteras et al., 1992)* | 28–30 | PAA |
| *layers 1–3 (Canteras et al., 1992)* | 28–30 | PAA1–3 |
| *Piriform area (Smith, 1919)* | 23–30 | PIR |
| *molecular layer (>1840)* | 23–30 | PIR1 |
| *polymorph layer (>1840)* | 23–30 | PIR3 |
| *pyramidal layer (>1840)* | 23–30 | PIR2 |
| *Posterior agranular insular area (Krettek & Price, 1977)* | 23–28 | Alp |
| *layers 1–6 (>1840)* | 23–28 | Alp1–6 |
| *Posterior amygdalar nucleus (Canteras et al., 1992)* | 30 | PA |
| *Posterior hypothalamic nucleus (>1840)* | 29, 30 | PH |
| *Posterior thalamic complex (>1840)* | 30 | PO |
| *Principal lateral olfactory tract (Swanson, 2018)* | 23–25 | lotp |
| *Radiation of corpus callosum (Dejerine & Dejerine-Klumpke, 1895)* | 23–30 | ccra |
| *External capsule (Burdach, 1822)* | 23–30 | ec |
| *Radiation of corpus callosum principal part (Swanson, 2018)* | 23–28 | ccrp |
| *Tapetum (Reil, 1809)* | 29, 30 | ta |
| *Retrosplenial area (Vogt & Peters, 1981)* | 25–30 | RSP |
| *Dorsal part (Swanson, 1992)* | 25–30 | RSPd |
| *Ventral part (Swanson, 1992)* | 25–30 | RSPve |
| *Anterior zone (Swanson, 2018)* | 25–30 | RSPv |
| *Reticular thalamic nucleus (>1840)* | 23–30 | RT |
| *Rhinal sulcus (>1840)* | 23–30 | rhs |
| *Rhomboid nucleus (Cajal, 1903)* | 26–30 | RH |
| *Septofimbrial nucleus (>1840)* | 23–27 | SF |
| *Somatomotor areas (>1840)* | 23–30 | MO |
| *Primary somatomotor area (>1840)* | 23–30 | MOp |
| *Secondary somatomotor areas (>1840)* | 23–30 | MOs |
| *Somatosensory areas (>1840)* | 23–30 | SS |
| *Primary somatosensory area (>1840)* | 23–30 | SSp |
| *barrel field layers 1–6 (>1840)* | 23–28 | SSp-bfd1–6 |
| *lower limb field layers 1–6 (>1840)* | 23–27 | SSp-ll1–6 |
| *trunk field layers 1–6 (>1840)* | 28–30 | SSp-tr1–6 |
| *upper limb field layers 1–6 (>1840)* | 23–27 | SSp-ul1–6 |
| *Subfornical organ (Pines, 1927)* | 23 | SFO |
| *Submedial thalamic nucleus (>1840)* | 27–30 | SMT |
| *Subparaventricular zone (Watts et al., 1987)* | 23–27 | SBPV |
| *Sulcus of corpus callosum (>1840)* | 23–30 | scc |
| *Supplemental somatosensory area (>1840)* | 23–29 | SSs |
| *layers 1–6 (>1840)* | 23–29 | SSs(1–6) |
| *Suprachiasmatic nucleus (Spiegel & Zwieg, 1919)* | 23, 24 | SCH |
| *Supraoptic nucleus (Lenhossék, 1887)* | 23–28 | SO |
| *principal part (Swanson, 2018)* | 23–26 | SOp |
| *retrochiasmatic part (>1840)* | 27, 28 | SOr |
| *Temporal association areas (Swanson, 1992)* | 29, 30 | TEa |
| *layers 1–6 (>1840)* | 29, 30 | TEa1–6 |
| *Thalamic optic tract (Swanson, 2015)* | 28–30 | optt |
| *Thalamic part of third ventricle principal part (Swanson, 2018)* | 29, 30 | V3t |
| *Thalamic segment of thalamic radiations (Swanson, 2018)* | 23–27 | trat |
| *Thalamic supraoptic decussations (Swanson, 2018)* | 29, 30 | supt |
| *Third ventricle (Galen, c173)* | 23–30 | V3 |
| *infundibular recess (>1840)* | 27–30 | V3i |
| *Tuberoinfundibular sulcus (>1840)* | 26–30 | tis |
| *Ventral anterior-lateral thalamic complex (>1840)* | 25–30 | VAL |
| *Ventral hippocampal commissure (Craigie, 1925)* | 23–26 | vhc |
| *Ventral medial thalamic nucleus (>1840)* | 27–30 | VM |
| *Ventral posterolateral thalamic nucleus, principal part (Swanson, 2004)* | 27–30 | VPLpr |
| *Ventral posteromedial thalamic nucleus, principal part (Swanson, 2004)* | 28–30 | VPMpr |
| *Ventrolateral hypothalamic tract (Swanson, 2004)* | 23–30 | vlt |
| *Ventromedial hypothalamic nucleus (>1840)* | 26–30 | VMH |
| *Anterior part (>1840)* | 26 | VMHa |
| *Central part (>1840)* | 27–30 | VMHc |
| *Dorsomedial part (>1840))* | 27–29 | VMHdm |
| *Ventrolateral part (>1840)* | 27–30 | VMHvl |
| *Visceral area (>1840)* | 23–28 | VISC |
| *layers 1–6 (>1840)* | 23–28 | VISC1–6 |
| *Zona incerta (>1840)* | 26–30 | ZI |
| *dopaminergic group (>1840)* | 28 | ZIda |

