## Supplemental Methods and References for "Chemoarchitectural studies of the rat hypothalamus and zona incerta. *Chemopleth 1.0* – A downloadable interactive *Brain Maps* spatial database of five co-visualizable neurochemical systems, with novel feature- and grid-based mapping tools"

[1]UTEP Systems Neuroscience Laboratory, [2]Department of Biological Sciences, [3]Border Biomedical Research Center, [4]Vision and Learning Lab, [5]Department of Computer Science, [6]PhD Program in Bioscience, [7]PhD Program in Computational Science, [8]RISE Program, [9]HHMI PERSIST Brain Mapping & Connectomics Undergraduate Teaching Laboratory,[10]Interdisciplinary Group for Neuroscience Instruction, Training, and Education (IGNITE), [11]Lower Brainstem Group, The University of Texas at El Paso, El Paso, TX 79968.

---

### *A. Supplemental Methods to Section 2*

#### *SM 2.1: Early lessons*

##### *SM2.1.1: Settling on a mappable labeling method*

This project has its antecedents in our student-driven efforts to characterize activation patterns for hypocretin/orexin-immunoreactive neurons (Nham & Khan, 2009a,b) and other cell types (Agostinelli et al., 2010; Agostinelli & Khan, 2011) in the rat lateral hypothalamic area (LHA) after glycemic challenge. A systematic photomicrographic analysis of several LHA tissue biomarkers was undertaken using immunofluorescence histochemistry (Pinales et al., 2012; 2013a,b; 2014; Pennington et al., 2014; Wells et al., 2014a,b) along with studies of putative synaptic interactions between labeled LHA systems (Escapita et al., 2015; Wells et al., 2015a–c). Additional student work continued within our two-semester Brain Mapping & Connectomics undergraduate laboratory course (D'Arcy et al., 2016; Flores-Robles et al., 2017; Burnett et al., 2018; Martinez et al., 2018; 2019), the learning outcomes and curriculum for which also have been reported (D'Arcy et al., 2019; Khan et al., 2021). However, fluorescence signal longevity and other challenges (also discussed in Wells, 2017) led us to realize that these patterns were best *mapped* using immunoperoxidase-based non-labile labeling methods that could be better controlled, signal-wise, across time (Guzman et al., 2019; Sotelo et al., 2019; Navarro et al., 2019; Peru et al., 2019; Toccoli et al., 2019; Vizcarra et al., 2019; Navarro, 2020; Peru, 2020; Sotelo et al., 2020/21; Toccoli et al., 2020/21).

##### *SM2.1.2: Settling on a feasible workflow for mapping density estimations*

Originally, for our immunohistochemical labeling patterns, we wanted to estimate densities of labeled elements within brain regions delineated on BM4.0 atlas maps. This led to an attempt to draw each vector-delimited area in BM4.0 base maps as a "shapefile" within a Geographic Information System (GIS) framework. Initial shapefiles were constructed in Mar 2020 by SB by exporting Ai files as .dxf AutoCad files for use in QGIS open-access software and plots of density distributions were created within the *R* framework (SB, unpublished laboratory blog post,

15 Apr 2020). Our initial efforts "were successful, as far as they went" (Hatton, 1984, p. 342). It soon became apparent, however, that the vector files from Ai needed to have closed polygons and exporting raster images of those regions would be more efficient than manually drawing shapefiles. It also became clear that we needed a tool "to take a published [BM4.0 atlas] map used by someone to populate their data and render it back into vector format so we can align [it] with current vector maps" (Khan, unpublished laboratory blog post, 21 May 2020). Therefore, we decided upon a workflow that would take vector-based shapes and *export them* for raster-based analysis and *return them* as vector graphics at the user level within the Ai graphical user interface. To achieve this, we settled upon the workflow described in the main paper.

*B. Supplemental References*


Agostinelli, L. J., Nham, T. A., Zobel, M., Michaels, J. M., & Khan, A. M. (2010). Distribution of neurons expressing nitric oxide synthase, acetylcholinesterase, and hypocretin/orexin in the rat hypothalamus: Relation to basal and stimulated levels of Fos and phospho-ERK. [Program #191.11]. *2010 Abstract Viewer/Itinerary Planner*. San Diego, CA: Society for Neuroscience. Online.

Agostinelli, L. J., & Khan, A. M. (2011). Distribution of neurons expressing nitric oxide synthase, acetylcholinesterase, and hypocretin/orexin in the rat hypothalamus: relation to basal and stimulated levels of Fos and phospho-ERK. *13th Annual Undergraduate Symposium for Scholarly and Creative Work*, University of Southern California, held on 13 April 2011.

Burnett, K. A. S., Pinales, B. E., Perez, E. J., Rodarte, D., Cardona, A. M., Galvan, K. J., Hernandez, G. G., Lezama, A. C., Lorenzana, K. T., Vasquez, A., Parada, P., Paz, J. I., Rascon, J., Thomason, R., Bautista, K., Barnes, J., D'Arcy, C. E., & Khan, A. M. (2018). High-spatial resolution mapping of anorexigenic neuropeptides expressed in the hypothalamus: A chemoarchitecture study in the adult male rat. Program No. 680.24. *2018 Neuroscience Meeting Planner*. San Diego, CA: Society for Neuroscience, 2018. Online.

D'Arcy, C. E., Martinez, A., Aranda, L. F., Cervantes, H. F. L., Chacon, L. E., Cordero, R. P., Fernandez, V., Garcia, G. A., Holguin, S., Jacquez, A., Miramontes, T. G., Montaño, B., Muñoz, P. C., Valenzuela, I. R., Yu, J. S., & Khan, A. M. (2016). Elaboration of hypothalamic chemoarchitecture of the adult male rat: A high spatial resolution mapping study of melanin-concentrating hormone, hypocretin/orexin, and calbindin immunoreactivities in multiple subjects. Program No. 453.07. *2016 Neuroscience Meeting Planner.* San Diego, CA. Society for Neuroscience, 2016. Online.

D'Arcy, C. E., Martinez, A., Khan, A. M., & Olimpo, J. T. (2019). Cognitive and non-cognitive outcomes associated with student engagement in a novel brain mapping and connectomics course-based undergraduate research experience. *Journal of Undergraduate Neuroscience Education*, *18*(1):A15–A43.

Escapita, A. C., Wells, C. E., & Khan, A. M. (2015). Chemoarchitecture of the mammalian hypothalamus: mapping of three peptidergic neuronal populations (α-MSH, nNOS, and MCH) in rat hypothalamus across multiple animals. *COURI Symposium Abstracts*, Spring 2015.

Flores-Robles, G., Negishi, K., Pacheco, R. A., Enriquez, A., Acevedo, E., Avila, B., Dominguez, E., Hernandez, E. E., Medina, A., Mejia, E., Novoa, M. A., Provencio, A. T., Renteria, F. D.,

Sifuentes, E., Tellez, Y., & Khan, A. M. (2017). Hypothalamic chemoarchitecture of the adult male rat: Further elaboration of results from a high spatial resolution longitudinal mapping study. Program No. 604.03. *2017 Neuroscience Meeting Planner*. Washington, D. C. Society for Neuroscience, 2017. Online.

Guzman, R., Peru, E., Navarro, V., Negishi, K., & Khan AM. (2019). Distributions of axons immunoreactive for α-melanocyte stimulating hormone and neurons immunoreactive for neuronal nitric oxide synthase in the hypothalamus of the adult male rat. *COURI Summer 2019 Symposium Abstracts*, ID=128, The University of Texas at El Paso, El Paso, Texas, USA, 3 August 2019.

Hatton, G. I. (1984). Hypothalamic neurobiology, In: *Brain slices* (R. Dingledine, ed.), pp. 341–374. New York: Plenum Press. https://doi.org/10.1007/978-1-4684-4583-1_14

Martinez, A., Barraza Escudero, L. M., Castro, D., Chavez, S. A., Coronado, M., Gallegos, S., Pineda Sanchez, A., Ruiz, M. S. P., Ruiz, V. G., Negishi, K., & Khan, A. M. (2018). Hypothalamic chemoarchitecture of the adult male rat: High spatial resolution mapping of copeptin, LIM homeobox 6, and melanin-concentrating hormone. Program No. 680.23. *2018 Neuroscience Meeting Planner.* San Diego, CA: Society for Neuroscience, 2018. Online.

Martinez, A., Navarro, V. I., Arias, K., Barnett, J., Carreon, A., Castaneda, B., Flores, M., Magadan, J., Heredia, D., Hooper, J., Lopez, T., Lozano, A., Mendez, S., Mercer, N., Munoz, A., Rosario Mojica, L., Sanchez, R., Sierra, K., Sotelo, D., Stevens, X., Toccoli, A., Verma, Y., Vizcarra, H., & Khan, A. M. (2019). The Hypothalamic Chemoarchitecture Project, Year 5: High-spatial resolution atlas-based mapping of neuronal populations expressing melanin-concentrating hormone, hypocretin/orexin, and calbindin in the hypothalamus of the adult male rat. Program No. 683.18. *2019 Neuroscience Meeting Planner*. Chicago, IL: Society for Neuroscience, 2019. Online.

Navarro, V. I. (2020). *An analysis and representation in an atlas reference space of putative appositions from neurons expressing alpha-melanocyte stimulating hormone onto neurons expressing hypocretin/orexin or melanin concentrating hormone.* Master's Thesis, Department of Biological Sciences, The University of Texas at El Paso. *Open Access Theses & Dissertations*, 3112. https://scholarworks.utep.edu/open_etd/3112

Navarro, V. I., Peru, E., Negishi, K., Ortega, M., & Khan, A. M. (2019). Distributions of immunoreactivities for hypocretin/orexin and neuronal nitric oxide synthase in the male rat hypothalamus: An analysis and representation in an atlas reference space. Program No. 149.18. *2019 Neuroscience Meeting Planner*. Chicago, IL: Society for Neuroscience, 2019. Online.

Nham, T., Khan, A. M. (2009a). Novel co-localization of phospho-ERK1/2 with the hypocretin/orexin system of the lateral hypothalamic area (LHA). *11th Annual Undergraduate Symposium for Scholarly and Creative Work*, University of Southern California, held on 15 April 2009.

Nham, T., Khan, A. M. (2009b). Identification of a novel population of neurons displaying immunoreactive phosphorylated MAP kinases in the lateral hypothalamic area: histochemical relationship to the hypocretin/orexin system. [Program #866.16]. *2009 Abstract Viewer/Itinerary Planner*. Chicago, IL: Society for Neuroscience. Online.

Pennington, K., Wells, C. E., & Khan A. M. (2014). Chemoarchitecture of the mammalian hypothalamus: Mapping of three peptidergic neuronal populations (α-MSH, nNOS, and MCH) in the lateral hypothalamic area and surrounding regions. *COURI Symposium Abstracts*, Spring 2014, The University of Texas at El Paso.

Peru, E. (2020). *Distributions of axons immunoreactive for alpha-melanocyte-stimulating hormone and neurons immunoreactive for neuronal nitric oxide synthase in the hypothalamus of the adult male rat: An analysis of interactions and their representation in an atlas reference space.* Doctoral Dissertation, Department of Biological Sciences, The University of Texas at El Paso. *Open Access Theses & Dissertations*, 3117. https://scholarworks.utep.edu/open_etd/3117

Peru, E., Navarro, V. I., Negishi, K., Guzman, R., & Khan, A. M. (2019). Distributions of axons immunoreactive for α-melanocyte stimulating hormone and neurons immunoreactive for neuronal nitric oxide synthase in the hypothalamus of the adult male rat: An analysis of interactions and their representation in an atlas reference space. Program No. 149.23. *2019 Neuroscience Meeting Planner*. Chicago, IL: Society for Neuroscience, 2019. Online.

Pinales, B. E., Dominguez, N., & Khan, A. M. (2012). Chemoarchitecture of the lateral hypothalamic area: Contiguous mapping and distribution of six biomarkers in the adult male rat. *COURI Symposium Abstracts, Spring 2012* (Paper 44), The University of Texas at El Paso.

Pinales, B. E., Rojas, P., Dominguez, N., Wells, C. E., & Khan, A. M. (2013a). Chemoarchitecture of the lateral hypothalamic area: determinations of forebrain and hindbrain contributions of the Neuropeptide Y system. *COURI Symposium Abstracts*, *Spring 2013*, The University of Texas at El Paso.

Pinales, B. E., Lean, G. A., Wells, C. E., Silva, N. D., Walker, E. M., Rojas, P., Dominguez, N., Martinez, A., Thompson, R. H., & Khan, A. M. (2013b). The Hypothalamic Chemoarchitecture Project: High resolution mapping of neuronal populations involved in pre-autonomic, neuroendocrine, and feeding control. Poster P301. *21st Annual Meeting of the Society for the Study of Ingestive Behavior (SSIB)*; New Orleans, LA; Jul 30–Aug 3, 2013.

Pinales, B. E., Wells, C. E., Pennington, K., & Khan, A. M. (2014). Identifying feeding control circuits in the brain by phenotyping, imaging and mapping the neurons of the lateral hypothalamic area in the adult rat brain. *2014 Annual Biomedical Research Conference for Minority Students (ABRCMS)*, San Antonio, Texas; Nov 12–15, 2014.

Sotelo, D., Guevara, A., Peru, E., Navarro, V. I., Toccoli, A., Szilvásy-Szabó, A., Magdolna, R. Y., Farkas, E., Guerra-Ruiz, J., Arzate, L. S., Negishi, K., Balivada, S., Arnal, A., Fekete, C., & Khan, A. M. (2020/21). Mesoscale and microscale interactions of fibers immunoreactive for α-melanocyte-stimulating hormone and hypocretin/orexin with somata immunoreactive for neuronal nitric oxide synthase: A representation in an atlas reference space of the adult male rat, with supporting ultrastructural evidence. Program No. 263.08. *Presented at the Global Connectome Meeting of the Society for Neuroscience*. Online, Jan 11–13.

Sotelo, D., Hooper, J., Vizcarra, H., Navarro, V., Martinez, A., & Khan, A. M. (2019). High-spatial resolution atlas-based mapping of melanin-concentrating hormone, hypocretin/orexin, and calbindin in the caudal hypothalamus of the adult male rat. *COURI Summer 2019 Symposium*

*Abstracts*, ID=87, The University of Texas at El Paso, El Paso, Texas, USA, 3 August 2019.

Toccoli, A., Arias, K., Navarro, V., Martinez, A., & Khan, A. M. (2019). The Hypothalamic Chemoarchitecture Project, Year 5: High-spatial resolution atlas-based mapping of neuronal populations expressing melanin-concentrating hormone, hypocretin/orexin, and calbindin in the rostral hypothalamus of the adult male rat. *COURI Summer 2019 Symposium Abstracts*, ID=68, The University of Texas at El Paso, El Paso, Texas, USA, 3 August 2019.

Toccoli, A., Arzate, L. S., Navarro, V. I., Peru, E., Sotelo, D., Szilvásy-Szabó, A., Magdolna, R. Y., Farkas, E., Guevara, A., Guerra-Ruiz, J., Negishi, K., Balivada, S., Arnal, A., Fekete, C., & Khan, A. M. (2020/21). An analysis and representation in an atlas reference space of synaptic interactions between alpha-melanocyte stimulating hormone-immunoreactive axonal fibers and neurons expressing hypocretin/orexin or melanin-concentrating hormone: light and electron microscopic evidence. Program No. 263.05. *Presented at the Global Connectome Meeting of the Society for Neuroscience*. Online, Jan 11–13.

Vizcarra, H., Ortega, M., Navarro, V., Peru, E., Negishi, K., & Khan, A. M. (2019). Distributions of immunoreactivities for hypocretin/orexin and neuronal nitric oxide synthase in the male rat hypothalamus: An analysis and representation in an atlas reference space. *COURI Summer 2019 Symposium Abstracts*, The University of Texas at El Paso, El Paso, Texas, USA, 3 Aug 2019.

Wells, C. E. (2017). *Method & madness – advanced techniques for characterizing rat hypothalamic chemoarchitecture and for modernizing legacy data*. Doctoral dissertation, Department of Biological Sciences, The University of Texas at El Paso. *Open Access Theses & Dissertations*, 584. https://scholarworks.utep.edu/open_etd/584

Wells, C. E., Pinales, B. E., Farkas, E., Lean, G. A., Walker, E. M., Rojas, P., Dominguez, N., Martinez, A., Fekete, C., Thompson, R. H., & Khan AM. (2014a). The Hypothalamic Chemoarchitecture Project: High resolution mapping of neuronal populations involved in pre-autonomic, neuroendocrine, and feeding control. *2nd Annual Interdisciplinary Research Symposium at UTEP*, April 24–25, 2014. The University of Texas at El Paso.

Wells, C. E, Pennington, K., & Khan A. M. (2014b). Hypothalamic chemoarchitecture in the adult male rat: Creating canonical atlas maps for three co-visualized neuronal populations and their fiber systems from a single brain. Program No. 256.17. *2014 Neuroscience Meeting Planner*. Washington, DC: Society for Neuroscience, 2014. Online.

Wells C. E., Pinales, B. E., Lean, G. A., Farkas, E., Pennington, K., Fekete, C., Thompson, R. H., & Khan, A. M. (2015a). Creating a multi-scale chemoarchitectural reference atlas of optogenetic targets in the mammalian hypothalamus. *Keystone Symposium on Molecular and Cellular Biology, 2015 Conference on Optogenetics*, March 12–16, 2015; Denver, Colorado.

Wells, C. E., Acosta, A., Aldrete, D., Carrion, A., Castro, L., Escapita, A. C., Espinoza, E., De La Fuente, K., Garrett, A., Gomez, A., Gomez, N., Hernandez-Casner, C., Luevano, M., Lopez, A., Martinez, D., Mendoza, E., Ortega, M., Perez, M., Rangel, E., Reza, E., Rivera, J., Roman, C., Rosas, A., Seade-Galindo, C., Teran, J., Unpingco, J., Valdez, S., & Khan, A. M. (2015b). Next generation interaction maps of hypothalamic circuits controlling survival behaviors. *Hypothalamic Circuits for Control of Survival Behaviors Workshop*; 2015 Sep 27; Janelia Farm Research Campus,

Ashburn, Virginia.

Wells, C. E., Acosta, A., Aldrete, D., Carrion, A., Castro, L., Escapita, A. C., Espinoza, E., De La Fuente, K., Garrett, A., Gomez, A., Gomez, N., Hernandez-Casner, C., Luevano, M., Lopez, A., Martinez, D., Mendoza, E., Ortega, M., Perez, M., Rangel, E., Reza, E., Rivera, J., Roman, C., Rosas, A., Seade-Galindo, C., Teran, J., Unpingco, J., Valdez, S., & Khan, A. M. (2015c). Hypothalamic chemoarchitecture in the adult male rat: Creating canonical atlas maps for co-visualized immunoreactive peptidergic neuronal populations (α-MSH, nNOS, MCH) and their fiber systems in multiple brains. Program No. 616.08. *2015 Neuroscience Meeting Planner*. Chicago, IL. Society for Neuroscience, 2015. Online.
